# Common mechanism of activated catalysis in P-loop fold nucleoside triphosphatases – united in diversity

**DOI:** 10.1101/2022.06.23.497301

**Authors:** Maria I. Kozlova, Daria N. Shalaeva, Daria V. Dibrova, Armen Y Mulkidjanian

## Abstract

Though P-loop fold nucleoside triphosphatases (also known as Walker NTPases) are widespread, their catalytic mechanism remains unclear. Based on a comparative structure analysis of 3136 Mg-NTP-containing catalytic sites, we propose a common scheme of activated catalysis for P-loop NTPases. In this scheme, a hydrogen bond (H-bond) between the strictly conserved, Mg-coordinating Ser/Thr of the Walker A motif ([Ser/Thr]^WA^) and the conserved aspartate of the Walker B motif (Asp^WB^) plays the key role. We found that this H-bond is very short in the structures with bound transition state analogs. Given that a short hydrogen bond (also known as a low-barrier hydrogen bond) implies parity of pK values of the H-bond partners, we suggest that the proton affinities of these two residues reverse upon activation so that the proton relocates from [Ser/Thr]^WA^ to Asp^WB^. The anionic [Ser/Thr]^WA^ alkoxide withdraws then a proton from the would-be nucleophile (either a water molecule or a sugar moiety in some P-loop kinases), and the nascent anion attacks the gamma-phosphate group. When gamma-phosphate breaks away, the trapped proton relays from Asp^WB^, via [Ser/Thr]^WA^, to beta-phosphate and compensates for its developing negative charge.

## 1. Introduction

Hydrolysis of nucleoside triphosphates (NTPs), such as ATP or GTP, by widespread P-loop fold nucleoside triphosphatases (also known as Walker NTPases) is one of the key enzymic reactions. P-loop NTPase domains (see Fig. 1 for their overview), which are coded by up to 20% gene products in a typical cell, drive the activity of rotary ATP synthases, DNA and RNA helicases, kinesins and myosins, ABC-transporters, ubiquitous translation factors, α-subunits of signaling heterotrimeric G-proteins and even oncogenic Ras-GTPases [1–19]. In the ECOD database [20], the topology-level entry “P-loop_NTPase” contains 193 protein families. In the Pfam database [21], the P-loop NTPase clan CL0023 contains 217 families. The main classes of P-loop NTPases were already present in the Last Universal Cellular Ancestor (LUCA) [9, 10, 17, 19, 22–27].

**Figure 1.**
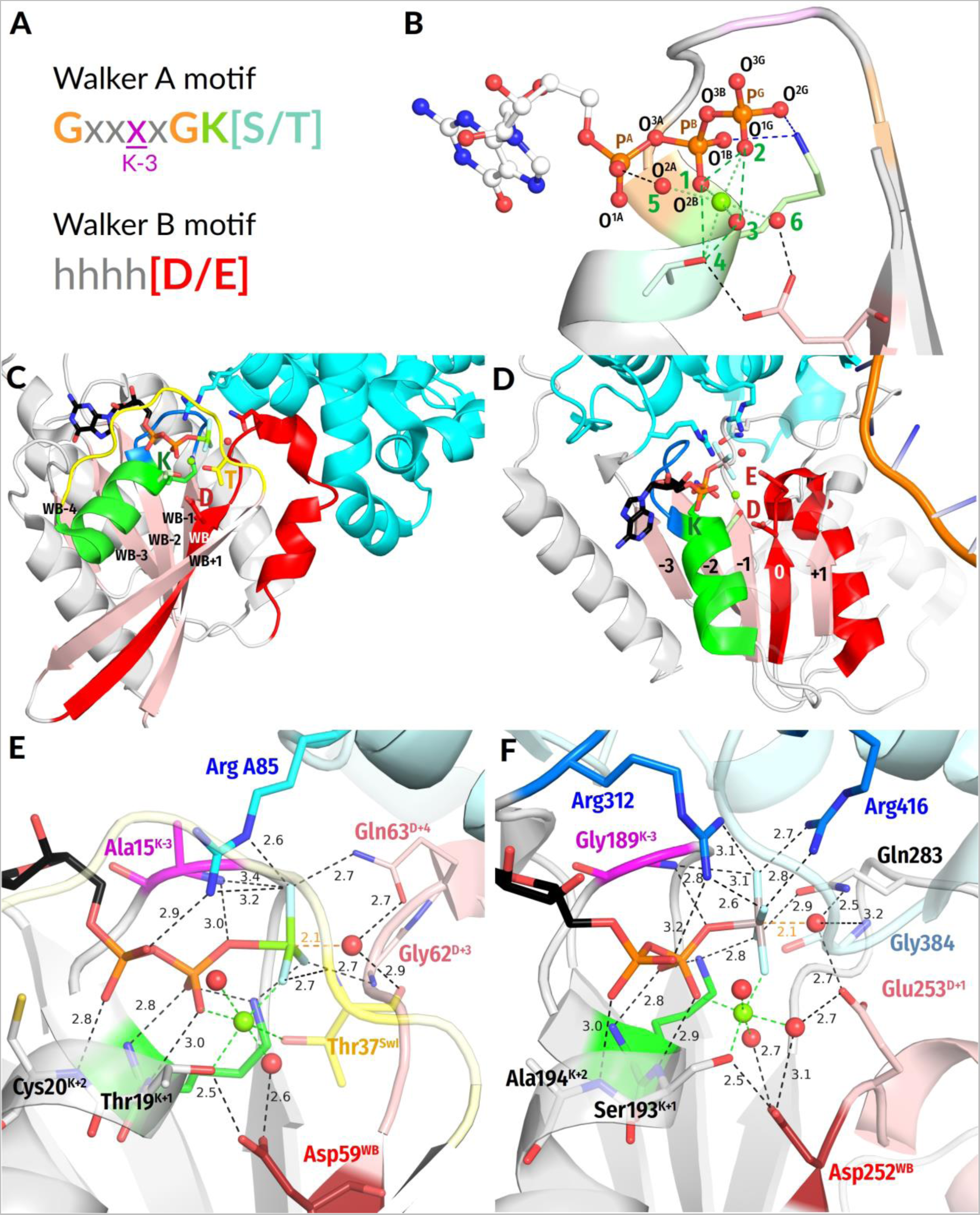
P-loop fold NTPases. **A,** conserved Walker motifs in P-loop NTPases; **B**, naming of atoms according to IUPAC recommendations for nucleoside triphosphates [33] and typical Mg^2+^ coordination; **C, D,** Typical folds of P-loop NTPases. **C**, GTPase Rho with its activator RhoGAP, (PDB ID 1OW3 [34]); **D**, Chikungunya virus nsP2 SF1-helicase (PDB ID 6JIM [35]). **E, F**, Close-ups of structures shown in **C, D** as examples of typical catalytic sites of P-loop NTPases. **E,** GTPase Rho with its activator RhoGAP and the transition state analog GDP-MgF_3_^-^, (PDB ID 1OW3 [34]); **F,** Chikungunya virus nsP2 SF1 helicase complexed with transition state analog ADP-AlF_4_^-^ (PDB ID 6JIM [35]). Color code: Polypeptide chains are shown as gray cartoons, β-strands, pink; α_1_-helix, green; P-loop, blue; Walker B motif and the following residues, red, Switch I loop, yellow, the activating partners are colored cyan. Nucleotides, their analogs, and important amino acid residues are shown as sticks, water molecules – as red spheres, Mg^2+^ ions are shown as lime spheres, sphere; the conserved Lys residue of Walker A motif is shown in green with the amino acid residue three residues before it highlighted in magenta; conserved Asp/Glu residue of Walker B motif is shown in dark red and the following residues are shown in pale red; Arg fingers are shown in blue/cyan. In amino acid residues shown as sticks, the oxygen atoms are colored red, and the nitrogen atoms are colored blue. In the MgF_3_^-^ moiety, the fluoride atoms are colored light blue. In the AlF_4_^-^ moiety, the Al atom is colored gray. All distances are given in ångströms.

The P-loop fold domain is a 3-layer αβα sandwich, see Fig. 1 and [10, 18, 28–31]. In small P-loop NTPases (Fig. 1C), the titular *<u>p</u>*hosphate-binding loop (P-loop) usually connects the first β-strand (β_1_-strand) with the first α-helix (α_1_-helix); the P-loop together with the two first residues of the α_1_-helix have the [G/A]xxxxGK[S/T] sequence, known as the Walker A motif [1]. This motif is responsible for the binding of the triphosphate chain and cofactor Mg^2+^ ion, see [1, 2, 32] and Fig. 1A-D. The Walker B motif *hhhh*D, where ’*h’* denotes a hydrophobic residue, is the other shared motif of P-loop NTPases, see Fig. 1A-F and [1, 2, 32]. The aspartate residue of the Walker B motif is thought to contribute to the stabilization of the Mg^2+^ binding site. In small P-loop NTPases, this residue is at the C-terminal tip (hereafter C-cap) of the β_3_-strand [9], just opposite of the α_1_-helix. A few P-loop NTPases have a glutamate residue in the respective position; we explicitly address these cases below. Still, throughout the manuscript, the conserved carboxylic residue of Walker B motif is denoted as Asp^WB^ for the sake of simplicity, see Fig. 1E-F. Asp^WB^ makes a hydrogen bond (H-bond) with the conserved Ser/Thr of the Walker A motif that follows the conserved Lys residue, see Fig. 1E and 1F where the respective residues are denoted as Thr19^K+1^ and Ser193^K+1^, respectively.

A specific feature of most P-loop ATPases is their activation before each turnover, so that the NTP hydrolysis proceeds in two steps. First, an ATP or a GTP molecule binds to the Walker A motif and attains a configuration with eclipsed β- and γ-phosphates, as enforced by the Walker A motif and the Mg^2+^ ion, see Fig. 1B and [31, 36–40]. Then the P-loop domain interact with its cognate activating partner, which could be a domain of the same protein, a separate protein and/or a DNA/RNA molecule. Upon this interaction, specific stimulatory moieties, usually Arg or Lys residues (“Arg/Lys fingers”, Fig. 1E-F), are inserted into the catalytic sites and promote the cleavage of γ-phosphate, see the accompanying article [41] and [4, 15, 16, 40, 42–49].

The energy of NTP binding drives the “closing” of the catalytic pocket, whereas the energy of hydrolysis is utilized for its “opening”; both these large-scale conformational changes can be coupled to useful mechanical work; they drive various cell motors, see, e.g. [4, 50–52].

Important hints for clarifying the catalytic mechanism of P-loop NTPases are provided by their structures with bound transition state (TS) analogs, such as NDP:AlF_4_^-^ or NDP:MgF_3_^-^ or NDP-VO_4_^-^ complexes [4, 34, 42, 43, 46, 48, 53–56]. The crystal structures with NDP:MgF_3_^-^ or NDP:AlF_4_^-^ bound (see Fig. 1E and 1F, respectively) revealed a “catalytic” water molecule W_cat_ near the analog of the P^G^ atom and almost in-line with its bond with the O^3B^ oxygen atom (see Fig. 1B for the atom names according to the IUPAC recommendations for nucleoside triphosphates [33]). In the NDP-VO_4_^-^ complexes, one of the four oxygen atoms of vanadate occupies the position of the catalytic water molecule [4, 55]. These structures, in support to earlier suggestions, indicate that the ultimate cleavage of γ-phosphate is triggered by the apical nucleophilic attack on the terminal phosphorus atom P^G^; the deprotonated state of W_cat_ (**OH^--^_cat_**) is thought to serve as a nucleophile in GTPases and ATPases, see Fig. 1E-F, Fig. S1 in Supplementary Materials and [15, 48, 56–63].

There is no consensus on how stimulatory moieties accelerate the NTP hydrolysis. It has been unclear whether they decrease the activation barrier of NTP hydrolysis by compensating for the developing negative charge on β-phosphate [62, 64] or by electrostatically stabilizing the TS [65] or by expulsing water from the catalytic pocket [66] or by some other impact.

It has remained also unclear how the interaction with the stimulating moiety initiates the deprotonation of W_cat_ and what is the fate of the proton that is released upon this deprotonation. The first TS-like crystal structures of small GTPases, such as shown in Fig. 1C, 1E, did not reveal potential bases among the ligands of W_cat_ [4, 42, 43, 53]. Thus, it was suggested that the proton is accepted directly by γ-phosphate [67–69]. In contrast, the crystal structures of many other P-loop NTPases showed that W_cat_ interacts with Glu or Asp residues; these “catalytic” residues were suggested to serve as proton acceptors in these NTPases, see e.g. Glu253 in Fig. 1F and [8, 54, 63, 70]. It remains unclear whether the proton is transferred from these bases to the triphosphate and, if so, how this transfer takes place.

In the accompanying article [41], by using cumulative computational approach, we inspected 3136 structures of Mg-NTP-containing catalytic sites of P-loop NTPases, identified the structures with inserted stimulator(s) and analyzed the patterns of stimulatory interactions. In most cases, at least one stimulator, by linking the oxygen atoms of α- and γ-phosphates, appears to twist γ-phosphate counter-clockwise by 30-40°; the rotated γ-phosphate is stabilized by a H-bond with the backbone amino group of the fourth residue of the Walker A motif. In rest cases, the stimulator(s) only engage(s) the γ-phosphate group and likely pull(s) or twist(s) it. The mechanistic interaction between the stimulator(s) and γ-phosphate appears to be a common property of all examined P-loop NTPases, which points to the stimulator-induced twist of γ-phosphate as the trigger for NTP hydrolysis.

Here we have combined manual identification of structural elements specific to major classes of P-loop NTPases with global computational analysis of H-bonding around Walker A and Walker B motifs in the same set of 3136 catalytic sites. By focusing on TS-like structures, we tried to capture the structural transitions involved in catalysis.

We have found that major classes of P-loop ATPases and GTPases share the following common structural features:

1. In most analyzed structures, a short H-bond (< 2.7 Å) connects Asp^WB^ and [Ser/Thr]^K+1^; this bond is extremely short (2.4-2.5 Å) in those structures that contain NDP:AlF_4_^—^ as a TS analog;
2. In TS-like structures of P-loop NTPases of all classes, except those of TRAFAC class, the W_cat_-coordinating “catalytic” Glu or Asp residue provides a proton pathway from W_cat_ to the nearest ligand of Mg^2+^.
3. The distance between neighboring ligands in the coordination shell of Mg^2+^ is 2.9-3.0 Å, which implies the possibility of proton exchange between all of them.

These common structural traits allowed us to specify a basic mechanism of stimulated catalysis for the whole superfamily of P-loop NTPases as follows:

i. interaction with the activating partner and twisting γ-phosphate by stimulator(s) appear to affect the properties of Mg^2+^ ligands including [S/T]^K+1^, so that, in the transition state, the [S/T]^K+1^ – Asp^WB^ H-bond shortens, the functional pKs of [S/T]^K+1^ and Asp^WB^ become leveled, and the proton relocates from [S/T]^K+1^ to Asp^WB^;
ii. the remaining proton vacancy at anionic [Ser/Thr]^K+1^ alkoxide is refilled by a proton that comes from W_cat_ (or a sugar moiety in some kinases), after which the nascent anion attacks γ-phosphate;
iii. upon the breakaway of γ-phosphate, the trapped on Asp^WB^ proton relays, via [Ser/Thr]^K+1^, to β-phosphate and compensates for its developing negative charge.

## 2. Results

To ensure detailed analysis and to cover as many structures as possible, we combined (i) the manual inspection of catalytic machinery in selected representatives from the major classes of P-loop NTPases (as described in Section 2.2.) with (ii) systematic computational analysis of all available structures of P-loop NTPases with bound Mg-NTP-like molecules (see Section 2.3 and the accompanying article [41]).

### 2.1. Generic designation of structure elements in P-loop fold NTPases

In the accompanying article [41], we have introduced and used a generic amino acid numbering for the conserved regions of P-loop NTPases. Hereafter we use the same nomenclature and number the amino acids residues of the Walker A, Switch I, and Walker B motifs (including functionally relevant residues following the Walker B motif) relatively to strictly conserved reference residues, namely Lys^WA^ (K^WA^) of the Walker A motif, Asp^WB^ (D^WB^) of the Walker B motif, and [Thr/Ser]^SwI^ ([T/S] ^SwI^) of the Switch I motif, respectively, as shown in Fig. 1 (see the accompanying article [41] for further details).

For ease of reference, we also introduce and use hereafter a novel generic numbering for the strands of the β-pleated sheet (for comparison, the conventional numbering of β-strands is shown in Fig. S2 for major classes of P-loop NTPases). Indeed, in small P-loop NTPases, the Walker B motif is at the C-cap of the β_3_-strand [9]. However, in other NTPases, it is at the C-cap of other β strands or even at the N-terminus [32]. Furthermore, larger P-loop NTPases can have additional β-strands which precede, in the amino acid sequence, the β_1_-strand that turns into the P-loop. This variable position of the Walker B motif is confusing. For instance, the Walker B motif is located at the C-cap of the β_2_-strand in the shikimate, gluconate, and adenosine phosphosulfate kinases [9]. The respective structures show that the conserved Asp^WB^ residue makes both canonical H-bonds with [Thr/Ser]^K+1^ and the Mg^2+^-coordinating water, respectively. And still, even recent review papers on these kinases indicate the Walker B motif in the β_3_-strand, where it is conspicuously absent.

Therefore, in this article, for simplicity, we call the β-strand, which is in-line with the triphosphate chain, “*Walker B strand*” or *WB-strand*. This strand is easy to find in a structure because it carries an aspartate or, rarely, a glutamate residue at its C-cap; the carboxyl of this residue is usually H-bonded to the [Thr/Ser]^K+1^ residue of the Walker A motif as shown in Fig. 1E-F. In addition, this carboxylic residue is usually preceded by four non-polar amino acids of the Walker B motif, see Fig. 1A and [1]. Other strands of the same β−pleated sheet, independently of their position in the amino acid sequence, can be numbered by their position relatively to WB-strand as WB-1, WB-2, or WB+1, WB+2 and so on, as shown in Fig. 1C. For the sake of brevity, “WB” could be omitted, and the strands could be numbered −1, −2, 0 (for the WB strand), +1, +2 and so on, as depicted in Fig. 1D, S3, and S4.

In those cases where the crystal structures of NTPases with their cognate activating partners are available, these partners, according to our observations, usually interact with a stretch of amino acids that follows the Walker B motif and approximately corresponds to residues from Asp^WB^+1 (aka D+1) to Asp^WB^+12 (aka D+12); this stretch is colored red in Fig. 1C-D or pale red in Fig. 1E-F. In the case of small NTPases, this region corresponds to their α_3_-helices (Fig. 1C). In some publications, the first few amino acids of this stretch are united with the Walker B motif into the “extended Walker B motif”. However, the amino acids beyond the conserved Asp^WB^ residue show no conservation throughout P-loop NTPases (see Fig. S3); hence, they make no conserved motif. By its shape, this stretch resembles a cock’s crest and usually rises above the neighboring β-strands as clearly seen in Fig. 1C-D. Therefore, hereafter we will name this structural element “*Walker B crest*” or *WB-crest*.

### 2.2. Catalytically relevant amino acids, stimulatory patterns, and activation mechanisms in different classes of P-loop NTPases

The P-loop NTPases are thought to form two divisions, namely the Kinase-GTPase division and ASCE (<u>A</u>dditional <u>S</u>trand <u>C</u>atalytic <u>E</u> (glutamate)) division, with both divisions containing several enzyme classes, see Fig. S2-S4 and [9, 10, 12, 14, 19].

For most classes of P-loop NTPases, we have selected representatives out of a set of proteins with available crystal structures in the Protein Data Bank (PDB) at www.rcsb.org [71, 72]. These structures are listed in Table 1. Each selected structure contains a Mg^2+^ cation and an NTP molecule or its analog bound; we preferred those structures that contain TS analogs. For these NTPases, after assessing the overall interaction with the activating partner, we checked the following features: (i) amino acids that coordinate the Mg-triphosphate moiety; (ii) positively charged, potentially stimulatory moiety(ies) inserted into the catalytic site during the activation (their cumulative analysis is presented in the accompanying article [41]); (iii) amino acids that interact with **W_cat_**; and (iv) other auxiliary amino acids that interact with oxygen atoms of the γ-phosphate group (or its analogs). The results of this manual analysis are described below and summarized in Table 1.

**Table 1.**
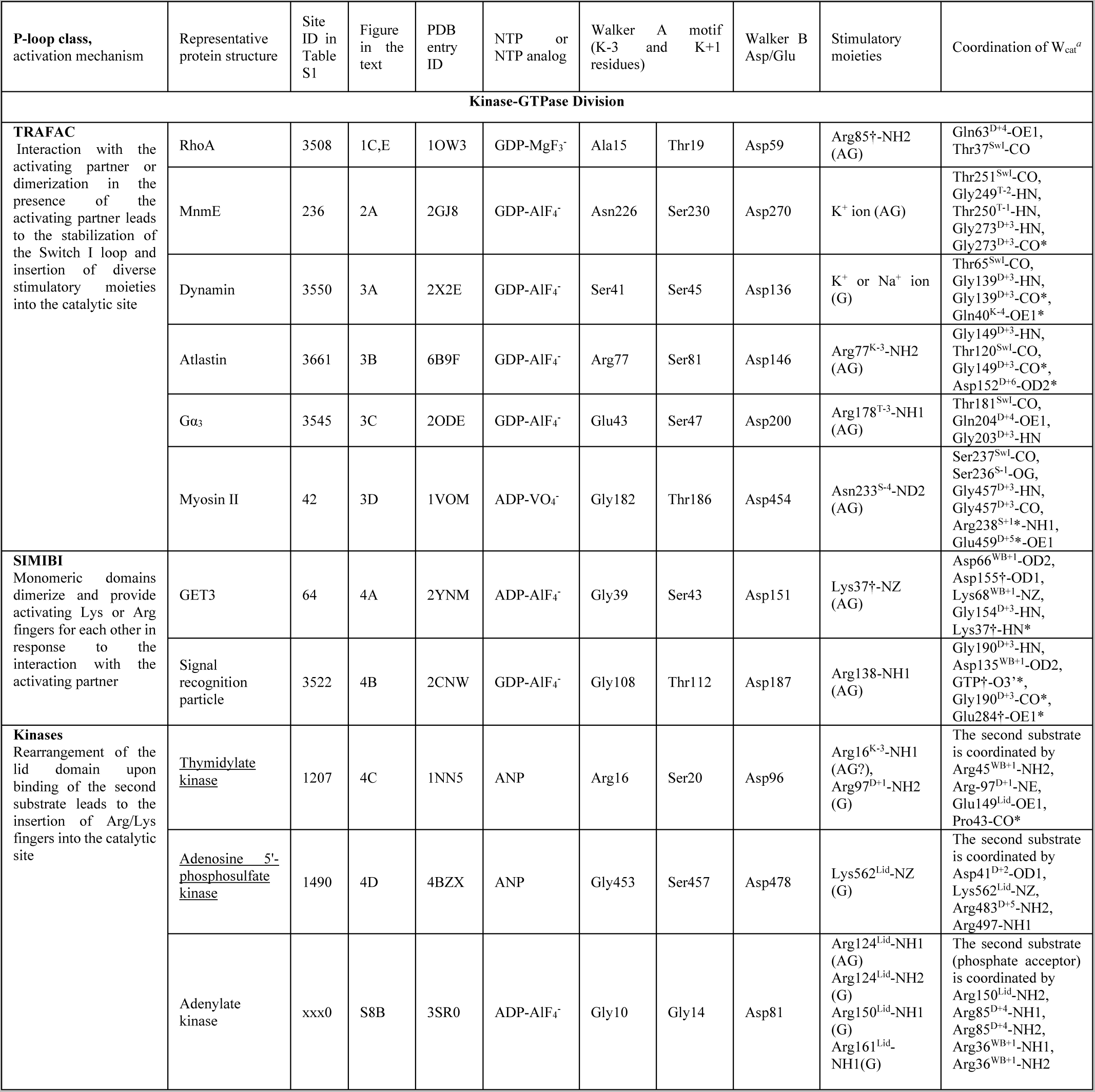
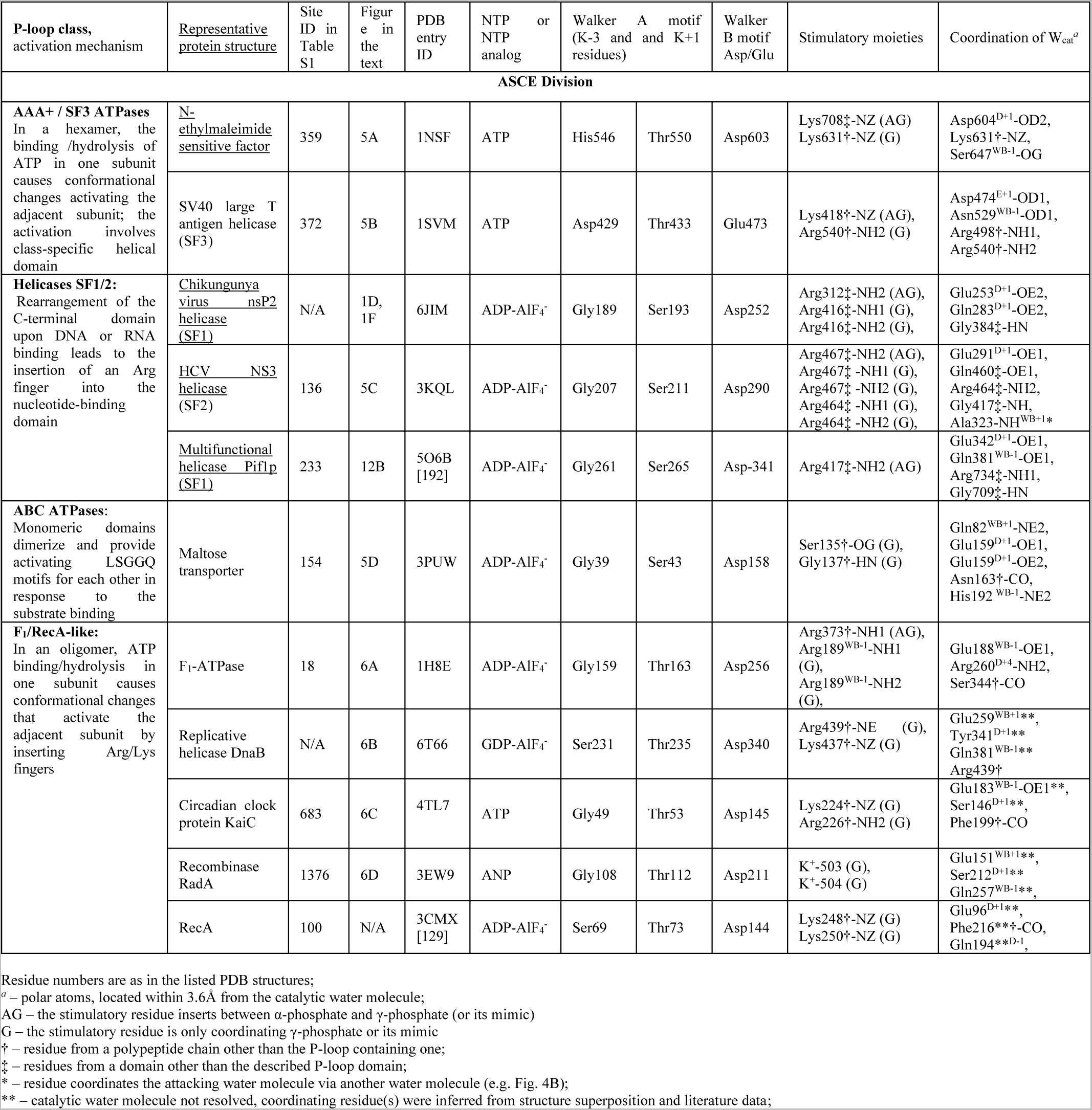
Structural traits of representative P-loop NTPases of different classes

#### 2.2.1. Coordination of the Mg-triphosphate moiety

First, representative structures from different classes were superposed with the structure of AlF_4_^—^-containing, K^+^-dependent GTPase MnmE (PDB ID 2GJ8, resolution 1.7 Å [46]), the activation mechanism of which we scrutinized elsewhere [40]. We aligned 20 amino acids of the ß_1_-strand, P-loop, and α_1_-helix with the corresponding amino acids 217-236 of MnmE. The whole shape of the P-loop found to be strictly conserved across all classes of P-loop NTPases (Fig. 2A). Accordingly, the binding mode of the triphosphate chain, which is described below, appears to be conserved throughout P-loop NTPases.

**Figure 2.**
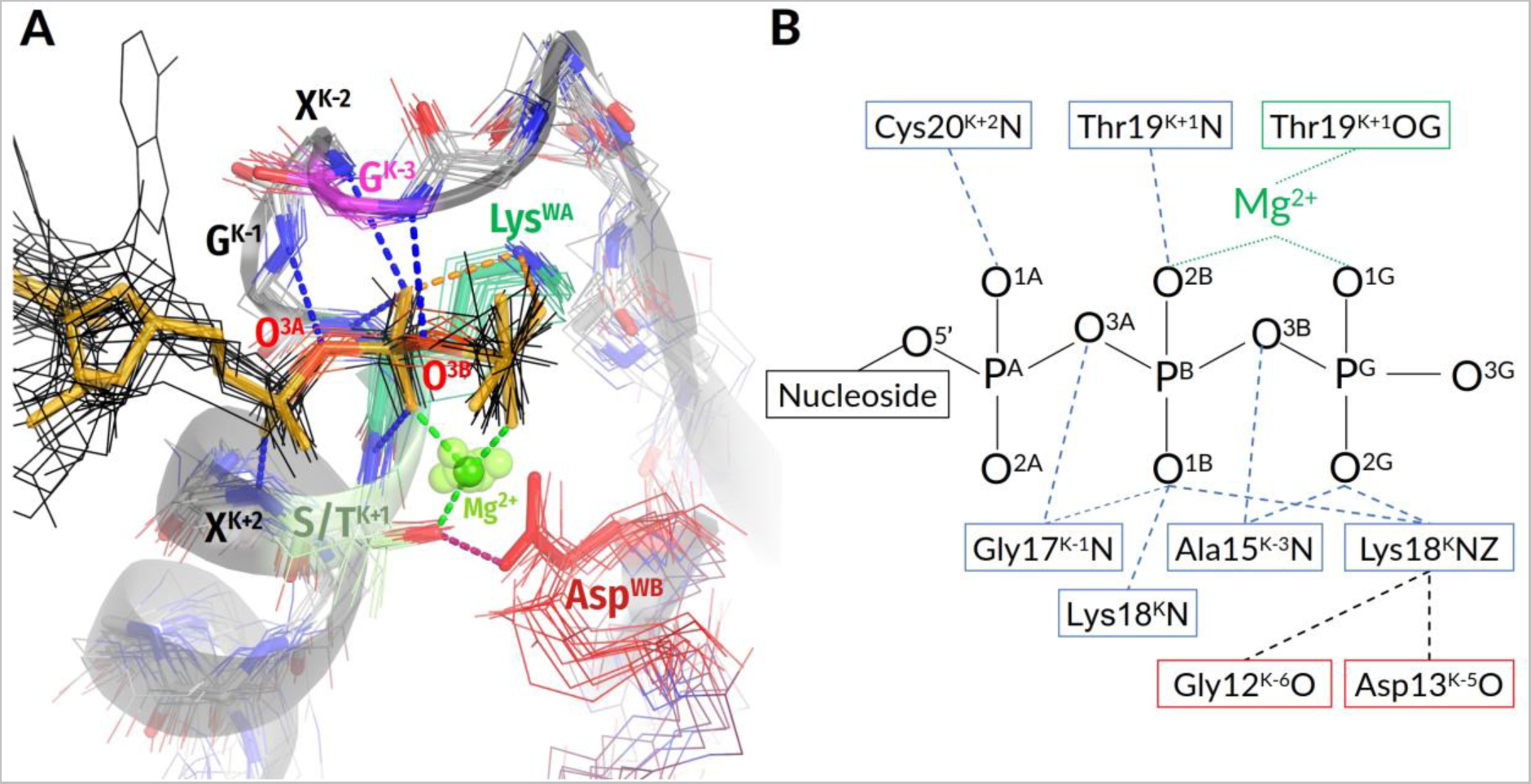
Conserved interaction between Walker A and Walker B motifs across classes of P-loop NTPases. **A.** Structures of proteins, representing different classes of P-loop NTPases and described in Table 1 are shown superimposed on the P-loop region of the MnmE GTPase with a TS-analog GDP:AlF_4_^-^ bound (PDB ID: 2GJ8 [46]). For the MnmE GTPase, the P-loop region is shown as a cartoon. The bonding pattern is shown only for the MnmE GTPase. For all the proteins, the last seven residues from the motif <u>GxxxxGK[S/T]x</u>, as well as the Asp residue of the Walker B motif and NTP analogs are shown by lines and colored as follows: Lys^WA^ in green, Ser/Thr^K+1^ in pale green, Gly/Ala/Asn^K-3^ in purple, Asp^WB^ in red; NTP molecules or their analogs are shown in black with bridging oxygen atoms in red. GDP:ALF_4_^-^ from the MnmE structure is shown in orange. Other protein residues are shown in gray with backbone nitrogen and oxygen atoms shown in blue and red, respectively. Mg^2+^ ions are shown as green spheres. The phosphate chain is involved in numerous bonds with backbone HN groups of Walker A motif residues (the bonds are highlighted in blue), conserved Lys^WA^ contacts O^2G^ and O^1B^ atoms (orange bonds) and O^2B^ and O^1G^ atoms are part of the Mg^2+^ coordination shell (green bonds). **B.** Coordination bonds and H-bonds between the Walker A motif and Mg-NTP moiety in the MnmE GTPase (PDB ID: 2GJ8 [46]).

In all manually inspected structures, the NTP molecule (or its analog) is bound to the P-loop of the Walker A motif in an extended conformation which is supposedly catalytically prone (Fig. 1E-F, 2A); elsewhere we showed that this conformation is similar to those of Mg-ATP and Mg-GTP in water in the presence of large monovalent cations such as K^+^ or NH_4_^+^ [40]. The configuration of the catalytic site is enforced by a plethora of conserved bonds that mostly involve the backbone amino group (hereafter **HN** groups) of the Walker A motif, see Fig. 1, 2A, B and [15, 31, 73]. The partial positive charges of these groups compensate for negative charges of the triphosphate chain oxygens. The structural elements responsible for stabilization of the triphosphate chain by amino acids of the Walker A motif are almost universally conserved (Fig. 2). The conservation of invariant residues in the [G/A]XXXXGK[T/S] motif has straightforward reasons: the Gly^K-1^ and Gly^K-6^ residues mark the beginning and end of the P-loop and enable the bending of the backbone; Gly^K-1^, in addition, electrostatically stabilizes the O^3A^ atom of α-phosphate. The Lys^WA^ residue interacts with O^1B^ and O^2G^ atoms and additionally appears to stabilize the P-loop by interacting with backbone carbonyl oxygens of K-5 and K-6 residues (Fig. 2).

In NTPases of all classes except several families of nucleotide monophosphate kinases, the Walker A and Walker B motifs are linked by a H-bond between the side chains of [Thr/Ser]^K+1^ and Asp^WB^ (Fig. 1E-F, 2A). This interaction is considered in more detail in Section 2.3.1. below where its comprehensive computational analysis is presented.

The same side chain of the conserved [Ser/Thr]^K+1^ serves as the ligand #4 of Mg^2+^. As shown in Fig. 1B, other strictly conserved ligands are O^2B^ and O^1G^ atoms of the triphosphate chain as ligands #1 and #2, respectively, and water molecules as ligands #5 (W5) and #6 (W6). The position #3 is taken either by water W3 or by diverse amino acid residues in different enzyme families, see [32] for details. The positions of Mg^2+^ ligands are similar over the entire superfamily of P-loop NTPases.

Fig. 2A, B show that the O^2A^ and O^3G^ atoms of the triphosphate are “free”, not bound to P-loop residues. Not surprisingly, in many TS-like structures, they interact with stimulators (Fig. 1E-F). For the MnmE GTPase, our molecular dynamics (MD) simulations showed that the insertion of a K^+^ ion and its simultaneous interaction with O^2A^, O^3B^ and O^3G^ atoms rotates γ-phosphate and leads to formation of a new H-bond between the O^2G^ atom and **HN** of Asn226^K-3^, see [40] and the accompanying paper [41]. Generally, the position of **HN**^K-3^ in the vicinity of the O^2G^ atom (or the corresponding atom of an NTP analog) is structurally conserved across P-loop NTPases, being determined by the highly conserved H-bond of **HN**^K-3^ with the bridging O^3B^ oxygen (Fig. 2). In manually inspected representative structures from Table 1, the distances between **HN**^K-3^ and the closest oxygen atom of γ-phosphate (or its structural analog) were shorter in the case of TS analogs. A comprehensive computational analysis of **HN**^K-3^-O^2G^ distances in all relevant PDB structures is presented in the accompanying paper [41].

#### 2.2.2. Catalytically relevant amino acids, activating partners, stimulatory patterns, and coordination of W_cat_

Other structural elements of P-loop NTPases, as well as their activating partners and stimulating moieties, vary among enzyme families; each class of P-loop NTPases is characterized by its specific constellation(s) of stimulators and W_cat_-coordinating residues. Since no detailed, comparative survey of catalysis-relevant structural attributes in all the multitude of P-loop NTPases is available, we provide here such a survey for the major classes of P-loop NTPases. Below we consider only those families and classes of P-loop NTPases for which the available structures allowed us to obtain unambiguous structural information. The results of this analysis are summarized in Table 1. Those classes of P-loop NTPases for which the structural information is ambiguous are discussed in the Supplementary File 1.

##### 2.2.2.1. Kinase-GTPase division

The Kinase-GTPase division unites three classes of NTPases: the TRAFAC (from *<u>tra</u>*nslation *fac*tors) class of translational factors and regulatory NTPases, SIMIBI (*si*gnal recognition, *<u>M</u>*inD, and *<u>Bi</u>*oD) class of regulatory dimerizing ATPases and GTPases, and the class of nucleotide kinases (Fig. S2, S3).

In the NTPases of **the TRAFAC class,** the α_1_-helix is followed by an elongated Switch I loop that is specific to the class. The elongated Switch I loop goes into the β-strand, which is antiparallel to all other β-strands of the core β-pleated sheet (Fig. 1C, S3). Apart from the TRAFAC class, other classes of P-loop NTPases have predominantly all-parallel core β-pleated sheets.

The Switch I loop contains the only strictly conserved [Thr/Ser]^SwI^ reference residue with its side chain coordinating Mg^2+^ as the ligand #3 (Fig. 1C, 1E, and S3). In most TRAFAC NTPases, the **HN** group of [Thr/Ser]^SwI^ forms a H-bond with γ-phosphate (Fig. 1E, 3). The backbone carbonyl group (hereafter **CO** group) of [Thr/Ser]^SwI^ interacts with the W_cat_ molecule seen in TS-like crystal structures (Fig. 1E, 3).

The TRAFAC class NTPases show a remarkably broad variety of stimulatory interactions; they are shown in Fig. 1C, 1E, 3, listed in Table 1 and described below.

***Monovalent cation-dependent NTPases***, which we have considered elsewhere [40], are usually stimulated by K^+^ ions, as the aforementioned GTPase MnmE, see Fig. 2, S3A, Table 1 and [46, 47]. The K^+^ ion in MnmE GTPase is coordinated by O^2A^, O^3B^, and O^3G^ atoms of the triphosphate chain, two **CO** groups of the K-loop, and the side chain of Asn^K-3^, see Fig. 2A, [40, 46] and the accompanying article [41].

In the unique eukaryotic protein family of dynamins, the NTP hydrolysis can be stimulated by either K^+^ or Na^+^ ions [74]. Here a Na^+^ or a K^+^ ion interacts only with the O^3B^ and O^3G^ atoms but does not reach the O^2A^ atom, see Fig. 3A, the accompanying article [41], and [40, 47] for details.

**Figure 3.**
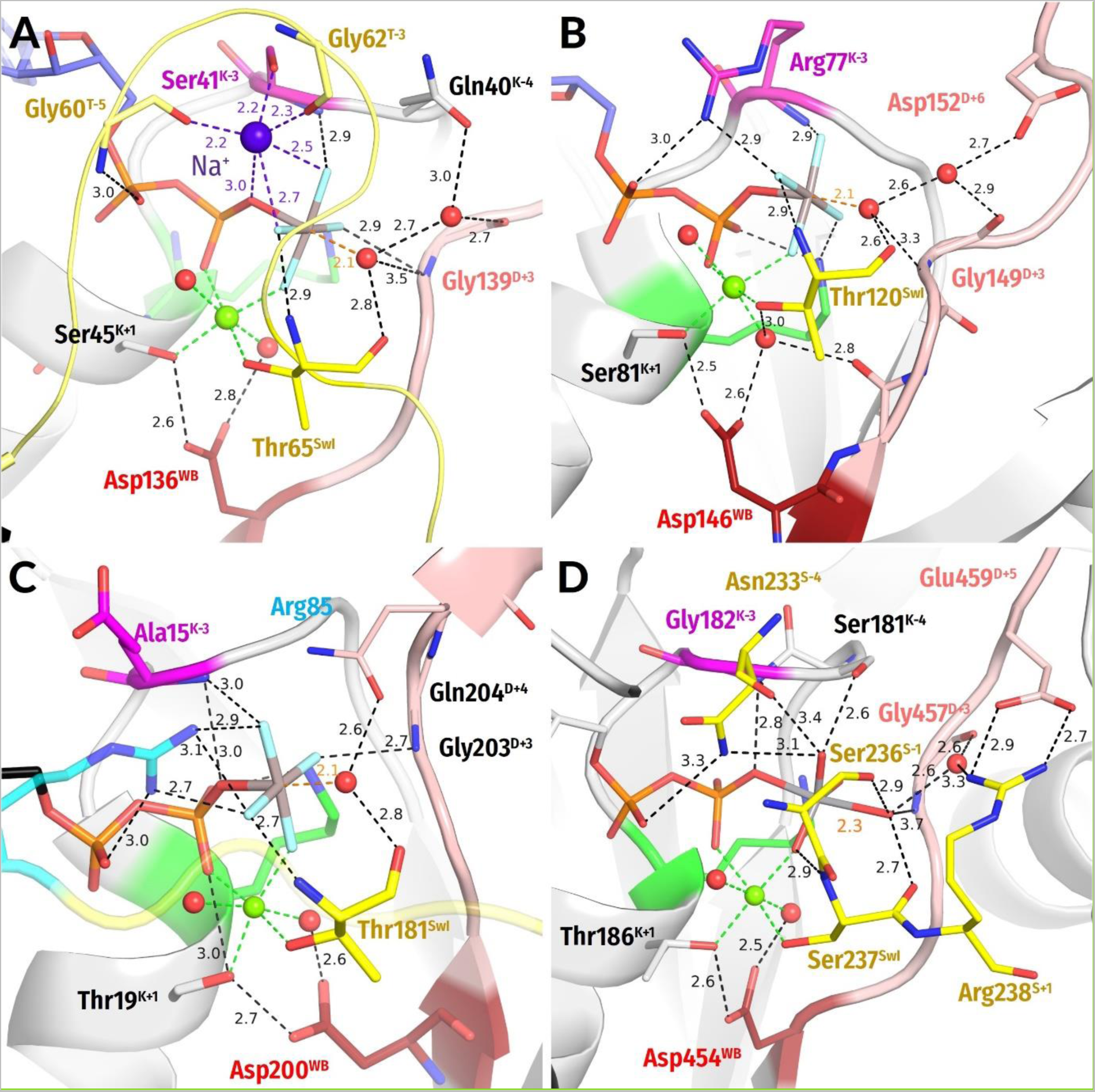
Representative NTPases of the TRAFAC class. The residues of Switch I/K-loop are shown in yellow, nucleotides, their analogs, and functionally relevant residues are shown as sticks, water molecules and cations are shown as spheres: water in red, Mg^2+^ in lime, Na^+^ in blue. Other colors as in Fig. 1E-F. All distances are given in ångströms. **A,** Dynamin (PDB ID 2X2E, [74]); **B,** Atlastin-1 (PDB ID 6B9F, [94]); **C,** Gα_12_ subunit of a heterotrimeric G-protein (PDB ID 2ODE, [86]); **D,** Myosin II, (PDB ID 1VOM [95]).

These K^+^/Na^+^-stimulated TRAFAC class NTPases belong to the family of HAS (hydrophobic aminoacid substitution) NTPases [75]. No hydrophilic side chains are present in the vicinity of Wcat; as indicated in Table 1, W_cat_ interacts only with the nearby atoms of the protein backbone, e.g. **CO**^SwI^, **HN**^SwI^, and **HN**^D+2^ in the case of the MnmE GTPase (see also the accompanying paper [41]). In dynamins (Fig. 3A), these are the side chain of Gln^K-4^, **CO**^SwI^ and **HN**^D+3^.

The titular family of ***translation factors*** unites ribosome-dependent GTPases directly involved in the translation, such as elongation factors EF-Tu and EF-G (Fig. S3B, Table 1). These GTPases showed K^+^-dependence both when studied in the absence of the ribosome [76, 77] and under conditions of protein synthesis [78–81]. Earlier, we suggested that these translational GTPases, alike K^+^-dependent GTPases, are stimulated by a K^+^-ion bound between the side chain of Asp^K-3^ and **CO** of Gly^T-2^ [82]. More recently, crystal structures of these GTPases indeed showed monovalent cations in the predicted position [83]. In the translational GTPases, W_cat_ is uniquely coordinated by the side chain of His^D+4^ residue in addition to **HN** of Gly^D+3^ and **CO** of Thr^SwI^; the side chain of His turns towards W_cat_ in response to the activating interaction of the WB-crest with the small ribosomal subunit and tRNA [84].

In ***GB1/RHD3-type GTPases*** (e.g. atlastins) the stimulatory Arg finger is in the K-3 position of the P-loop and links the O^2A^ and O^3G^ atoms of GTP when two protein monomers dimerize in response to the interaction with the activating partner (PDB ID 4IDQ [85], Fig. 3B). In this case, Arg fingers stimulate GTP hydrolysis in the very same P-loop domain they belong to. In the case of atlastins, W_cat_ is seen stabilized by **CO**^SwI^ and **HN**^D+3^ (Table 1, Fig. 3B).

In the GTPase domains of ***α-subunits of heterotrimeric G-proteins***, the intrinsic Arg finger, as provided by a family-specific insertion domain, links O^2A^ and O^3G^ atoms of GTP and is on a H-bond compatible distance from the O^3B^ atom, see Fig. 3C and [43, 86]. The W_cat_ is stabilized by Gln^D+4^ and **HN**^D+3^ of the WB-crest, as well as by **CO**^SwI^, see Fig. 3C, Table 1 and [43, 86].

The family of ***Ras-like GTPases*** named after its oncogenic members (from *<u>ra</u>t <u>s</u>arcoma*), is one of the best-studied groups in the TRAFAC class [15, 49, 50, 87]. In these proteins, the Switch I loop interacts with diverse physiological modulators of activity whereas the specific GTPase activating proteins (GAPs) bind to the WB-crest (also called Switch II in these proteins), see Fig. 1C, 1E, S3C and [15, 49, 88, 89]. As in α-subunits of heterotrimeric G-proteins shown in Fig. 3C, the NH2 group of the stimulatory Arg finger links the O^2A^ and O^3G^ atoms in most structures of Ras-like GTPases (Fig. 1C, 1E; see the accompanying article [41] for the discussion of apparent exceptions). The W_cat_ molecule is stabilized by the side chain of Gln^D+4^, **HN** of Gly^D+3^, and **CO**^SwI^ , see Table 1 and [15, 88, 90].

In P-loop NTPases of the ***kinesin and myosin families*** a Ser residue (Ser^SwI^) serves as a reference residue of Switch I; Ser^SwI^ provides its side chain oxygen atom to coordinate the Mg^2+^ ion as ligand #3 (Fig. 3D, S3D). In these proteins, the Asn^S-4^ residue inserts between α- and γ-phosphates [91–93], so that the side chain amino group of Asn^S-4^ links the O^2A^ and O^3G^ atoms, see Fig. 3D and the accompanying article [41]. Additional coordination of the γ-phosphate appears to be provided by the side chain of [Ser/Thr]^K-4^ of the P-loop and Ser^S-1^ of Switch I.

The **CO**^SwI^ group and, likely, the side chain of Ser^S-1^ form H-bonds with W_cat_. One more H-bond with W_cat_ is provided by **HN** of Gly^D+3^ (Table 1, Fig. 3D).

**The SIMIBI class NTPases** include ATPases and GTPases that dimerize upon interaction with the activating partner in such a way that catalytic sites of the monomers interact “face to face”, see Fig. S3E and [96, 97] for reviews. Each monomer inserts either a Lys (Fig. 4A) or an Arg residue (Fig. 4B) into the catalytic site of the other monomer [96, 97]. Many SIMIBI class ATPases and GTPases contain the so-called “deviant” Walker A motif KGGxGK[S/T] with an additional conserved Lys^K-5^ residue that is inserted into the catalytic site of the partner subunit in a dimer [97]. Lysine fingers, as used by SIMIBI proteins, form H-bonds with both O^2A^ and O^3G^ atoms (Fig. 4A).

**Figure 4.**
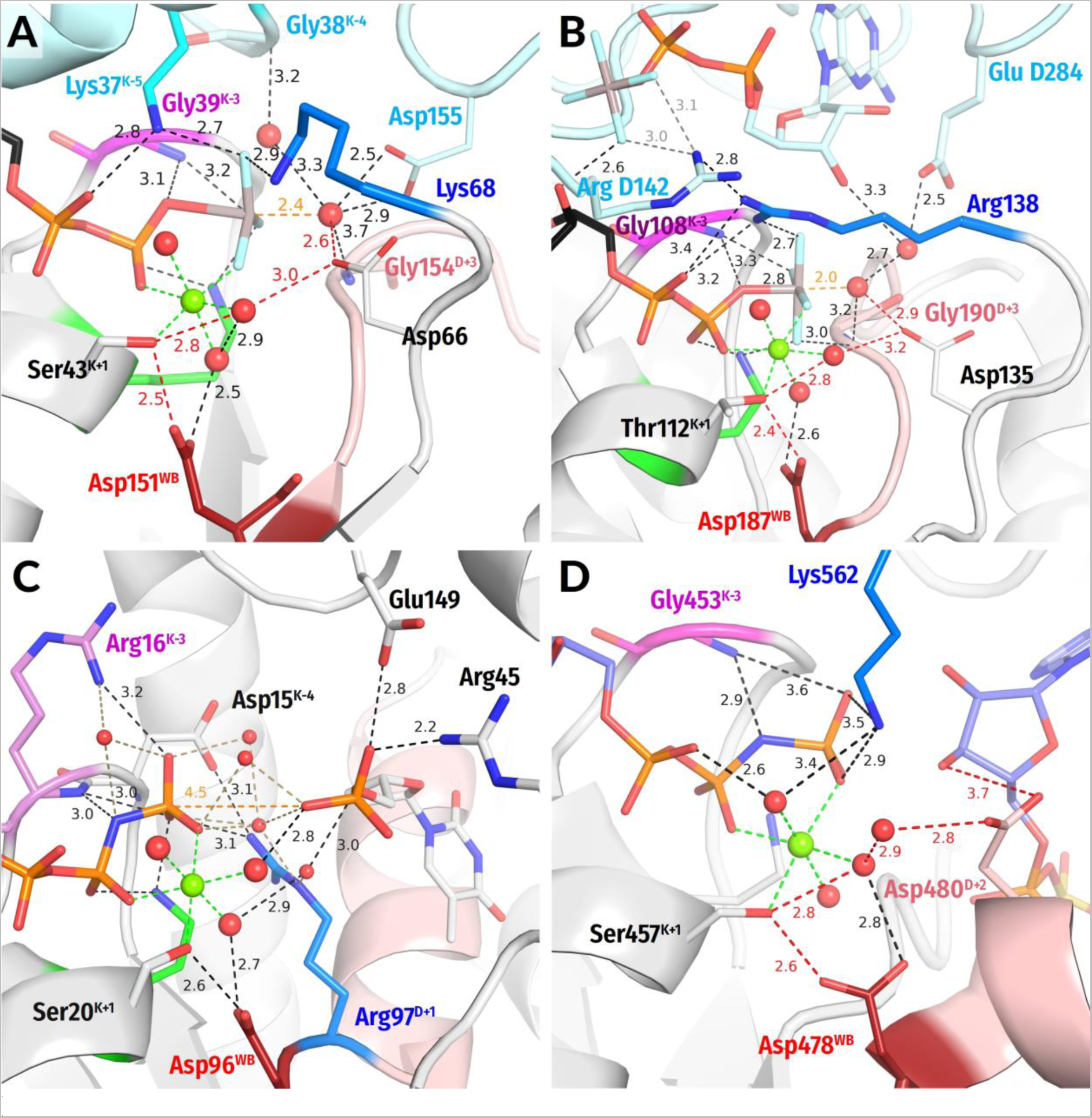
Representatives of the SIMIBI and kinase classes. The residues of the adjacent monomers in dimers are shown in cyan. Arg and Lys residues belonging to the same protein chain as Walker A motif, i.e. to the lid domains in kinases are shown in blue. Red dashed lines mark protonic connections between W_cat_ and Asp^WB^. Other colors as in Fig. 3. All distances are indicated in ångströms. **A.** ATP-binding component of the dark-operative protochlorophyllide reductase (PDB ID 2YNM, [99]). **B.** Signal recognition particle (FtsY/Ffh) complex (PDB ID 2CNW, [100]), **C**. Human thymidylate kinase (PDB ID 1NN5 [101]). **D**. Adenosine 5’-phosphosulfate kinase (PDB ID 4BZX, [102]).

Signal recognition particles (***SRPs***) and their cognate receptors ***(SRs***) stand separately within the SIMIBI class. Their GTPase domains form a pseudo-homodimer, where the two GTP binding sites interact face-to-face, but GTP hydrolysis occurs in only one of the two monomers. These proteins employ Arg residues that are inserted reciprocally so that the guanidinium group interacts with the α- and γ-phosphate of the GTP molecule bound by “its” subunit and the α-phosphate of the GTP molecule bound by the other subunit (Fig. 4B).

Many SIMIBI proteins have a Gly^D+3^ residue which appears to provide its **HN**^D+3^ for coordination of the γ-phosphate, similarly to Gly^D+3^ in TRAFAC class proteins, see Fig. 4A-B and [10]. Additional coordination of γ-phosphate is provided by amino acids located outside of the conserved motifs. Such residues are protein family-specific and are often introduced into the catalytic site from the adjacent monomer upon the interaction with the activating partner and dimerization (Fig. 4A-B).

The W_cat_ molecule in SIMIBI NTPases is routinely stabilized by **HN** of Gly^D+3^ and the Asp/Glu residue at the C-cap of the WB+1 strand; the carboxy group of this “catalytic” residue links W_cat_ with the Mg^2+^-coordinating W3 molecule (Fig. 4A-B). Also, the residues of the other monomer and even the ribose 3’ hydroxyl group of the “other” GTP molecule can contribute to the coordination of W_cat_, see Table 1, Fig. 4A-B. The NTP-bound SIMIBI dimer is believed to require the interaction with an activating protein (or RNA in SRP/SR complexes) to bring these specific “catalytic” residue(s) closer to the catalytic site where they can contribute to the H-bond network around W_cat_ [96, 98].

**Nucleotide Kinases** are ubiquitous enzymes that usually transfer a γ-phosphoryl residue from ATP to a wide range of “second” substrates, primarily small molecules [9, 103], see Fig. S3F. The key roles in the catalysis by P-loop kinases are usually played by Arg or Lys residue(s) located in the so-called “LID” domain, a small helical segment that “covers” the catalytic site and often carries several positively charged residues [9, 103, 104], see Fig. 4C-D. In thymidylate kinases, these fingers are assisted by Arg^K-3^, similarly to atlastines, cf Fig. 3B and 4C. Thus, P-loop kinases do not need other proteins or domains to provide stimulatory Arg/Lys fingers. Instead, binding of their second substrate is enough to trigger the Lid domain rearrangement that results in the insertion of Arg/Lys finger(s). While Arg residues serve as stimulatory moieties in most kinases (Fig. 4C), a Lys residue appears to be involved in adenylylsulfate kinases (Fig. 4D) [105].

One of the stimulatory fingers usually inserts between the α- and γ-phosphates. In the case of the adenosine-5’-phosphosulfate (APS) kinase, the lid domain inserts a Lys finger that interacts directly with the γ-phosphate and is connected to α-phosphate via a water molecule (Fig. 4D). As in many other P-loop NTPases, the catalysis in kinases is usually assisted by auxiliary arginine and lysine fingers that position the interacting molecules and neutralize the negative charges of phosphate groups, see Fig. 4C-D, Table 1 and [56, 106]. Some families of kinases use also an Arg residue of the WB-crest, see Fig. 4C and [9].

In kinases, the hydrolysis of the phosphoryl donor molecule is mediated by the acceptor molecule, which, similarly to W_cat_ in other reactions of NTP hydrolysis, appears to initiate the nucleophilic attack and formation of the pentavalent intermediate. In nucleotide monophosphate kinases, the attacking group is a negatively charged phosphate moiety requiring no specific proton acceptor (Fig. 4C). In those kinase families, where the deprotonation of the attacking molecule is needed to produce a nucleophile, the [Asp/Glu]^D+2^ residue appears to serve as an immediate proton acceptor, see Fig. 4D and Table 1. This residue links the would-be nucleophilic group with the Mg^2+^ ligand in position 3, usually a water molecule (Fig. 4D).

##### 2.2.2.2. ASCE division

The NTPases of the ASCE division have all-parallel β-pleated sheets with an inserted, as compared to the sequences of the Kinase-GTPase division NTPases, β-strand in the WB+1 position, see Fig.1B, S2, and S4. In many enzyme classes of this division, further additional β-strands were identified, see Fig. S2, S4 and [14, 18, 19]. For coordination of W_cat_, in addition to class-specific residues, ASCE NTPases typically use a “catalytic” glutamate residue that either directly follows the conserved Asp^WB^ or is located at the C-cap of the WB+1 strand, similarly to SIMIBI NTPases. These two features define the name of this division: *<u>a</u>dditional <u>s</u>trand, <u>c</u>atalytic <u>E</u>* (ASCE) [10, 12, 107, 108]. According to current views [14, 18, 19] and the most recent phylogenetic scheme depicted in Fig. S2, the ASCE NTPases are divided into two clades that differ by the number of β-strands in their P-loop domains. The group of “middle-size” domains with up to five-six β-strands includes AAA+ ATPases, helicases of superfamily 3 (SF3), as well as STAND and KAP ATPases. The clade of “large” ATPases domains, with many β-strands, includes helicases of superfamilies 1 and 2 (SF1/2), ABC-ATPases, RecA/F_1_ ATPases, VirD/PilT-like ATPases, and FtsK-HerA-like ATPases.

**AAA+ ATPases** are *<u>A</u>TPases <u>a</u>ssociated with various cellular <u>a</u>ctivities*, where “+” stands for “extended”, see Fig. 5A and S4A. These enzymes contain an N-terminal P-loop domain and an additional α-helical C-terminal domain, see [6, 12, 13, 16, 45, 109, 110] for comprehensive reviews. The P-loop domain of the AAA+ ATPases carries conserved Arg/Lys residue(s) from the side that is opposite to the P-loop. The P-loop domains interact upon oligomerization (most often a hexamer is formed), so that the nucleotide-binding site of one subunit receives the Arg/Lys finger(s) from the neighboring subunit (Fig. 5A-B) and/or, in some protein families, an additional Arg/Lys residue from its own C-terminal helical domain (Fig. 5A), see [16]. One of the stimulatory residues interacts with γ-phosphate and is called “finger” whereas the other one occupies the space between α- and γ-phosphates and is called “sensor 2”. We could not find structures of AAA+ ATPases with bound TS analogs; based on available data on site-specific mutants [16], W_cat_ appears to be coordinated by the “catalytic” [Glu/Asp]^D+1^ residue, Arg/Lys finger, and, perhaps, Asn/Ser/Thr residue at the C-cap of the WB-1 strand (“sensor 1”, see Fig. 5A). As shown in Fig. 5A, Glu^D+1^ connects a would-be W_cat_ with Mg^2+^- coordinating W3 in the structure of N-ethylmaleimide sensitive factor (PDB ID 1NSF [111]).

**Figure 5.**
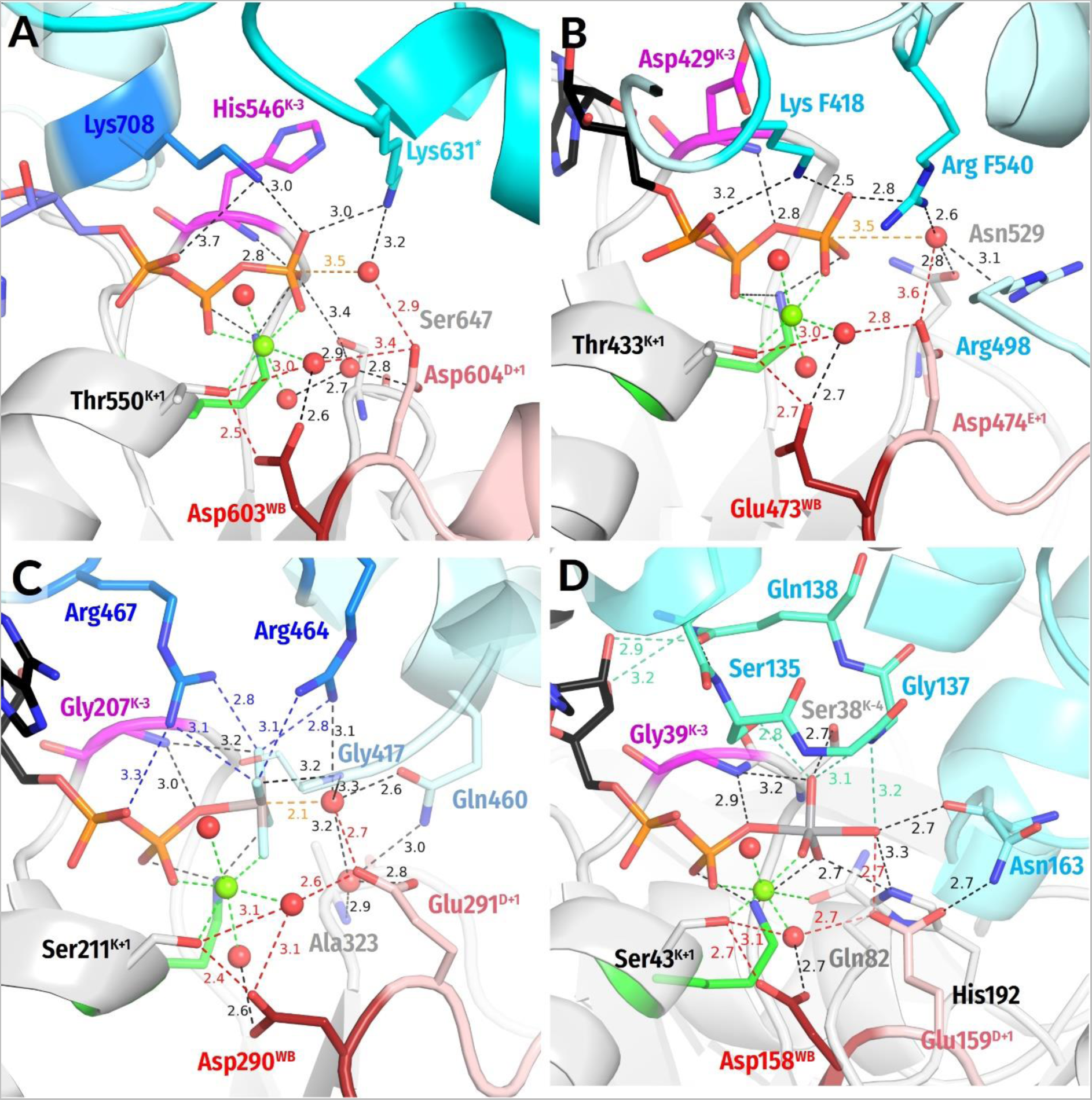
Representatives of AAA+ ATPases, SF3 helicases, SF2 helicases and ABC ATPases. The adjacent monomers and their Arg and Lys residues are shown in cyan, residues of adjacent monomers labelled in light blue. The Arg residues from the C-terminal helical domain of the same monomer are shown in deep blue (panel C). Red dashed lines mark protonic connections between W_cat_ and Asp^WB^. Other colors as in Fig. 3. All distances are given in ångströms. **A,** N-ethylmaleimide sensitive factor (PDB ID 1NSF [111]); **B,** Replicative hexameric helicase of SV40 large tumor antigen (PDB ID 1SVM, [114]); **C,** Hepatitis C virus NS3 SF2-helicase (PDB 5E4F [115]). **D,** Outward-facing maltose transporter complex with ADP:VO_4_^—^ bound (PDB ID 3PUV, [116]). The LSGGQ motif is shown in green-cyan, the VO_4_^—^ moiety by grey and red.

AAA+ NTPases are rather diverse, they can use further polar residues to interact with W_cat_ or γ-phosphate. While the Arg/Lys finger that interacts with γ-phosphate is usually present, the sensor 1 or sensor 2 residues are absent in some enzyme families, as reviewed in [6, 12, 13, 16, 45, 109, 110]. Also, the topology of domains that interact with the P-loop domain can vary. For instance, the helicases of superfamily 3 (SF3 helicases) have different topology of their C-terminal helical domain [13]. Their other specific feature is the presence of a Glu^WB^–Asp^E+1^ pair at the C-cap of the WB strand; the Asp^E+1^ residue builds a link to the Mg^2+^-coordinating W3 molecule (Fig. 5B).

**SF1/2 class helicases** are mostly monomeric or dimeric with each polypeptide chain containing two P-loop fold domains, see [3, 112] for reviews. While SF1 and SF2 helicases differ in the structural motifs that couple their DNA/RNA binding sites with ATP-hydrolyzing catalytic pockets, the pockets proper are quite similar (cf Fig. 1F and 5C, respectively). Both in SF1 and SF2 helicases, the ATP molecule binds to the functional Walker A and B motifs of the N-terminal domain; the C-terminal domain, although it has a P-loop-like fold, lacks the Walker A and B motifs. Following the interaction with an RNA or a DNA molecule, two Arg residues of the C-terminal domain are usually inserted into the ATP-binding site [113]. One of these Arg residues forms H-bonds with both α- and γ-phosphates, whereas the other Arg residue interacts with γ-phosphate (or its analog) and W_cat_ (Fig. 1F, 5C). The W_cat_ molecule is also coordinated by the “catalytic” Glu^D+1^ residue and class-specific Gln and Arg residues (Fig. 1F, 5C). The same Glu^D+1^ residue links W_cat_ with the Mg^2+^-coordinating W3 molecule (Fig. 1F, 5C, Table 1).

**ABC (*ATP****-**binding cassette*****) ATPases** are multidomain proteins that usually operate as homo- or heterodimers, see Fig. 5D, S4B and [117, 118]. Their ATP-hydrolyzing domains are usually used to drive large scale conformational changes, e.g. in membrane-embedded ABC-transporters or water soluble complexes involved in DNA repair or translation [19, 119, 120]. In dimers of ABC-NTPases, the nucleotide-binding sites of P-loop domains are located on the interface between the monomers, in the same way, as in dimers of SIMIBI NTPases, cf. Fig. 5D with Fig. 4A-B. Instead of an Arg or Lys residue, each monomer inserts a whole signature motif L<u>S</u>G<u>G</u>Q into the catalytic pocket of the other monomer (Fig. 5D, S4B), see also the accompanying article [41] for details.

The structures with TS analogs bound are available for the maltose transporter complex, see Fig. 5D, Fig. 7D in the accompanying article [41], and [116]. In these structures, the side chain of serine and **HN** of the second glycine residue of the signature motif interact with the O^3G^ atom of γ-phosphate. The side chain of serine is located between the α- and γ-phosphates, in the position of the Na^+^ ion in dynamin-like proteins, cf Fig. 5D with Fig. 3A.

Several amino acids commonly found in the catalytic sites of ABC transporters can stabilize W_cat,_ see Fig. 5D and Table 1. In the case of maltose transporter, these are a histidine residue at the C-cap of the WB-1 strand and Glu^D+1^; the latter, in addition, links W_cat_ with the Mg^2+^- coordinating W6 molecule. In the maltose transporter, the activating monomer in a dimer contributes to the coordination of W_cat_ by providing a backbone **CO** group of a residue that is located outside of the signature motif, see Fig. 5D and [116].

**RecA/F_1_ NTPases** class encompasses oligomeric ATP-dependent motors involved in homologous recombination and DNA repair (RecA and RadA/Rad51), catalytic subunits of rotary F/N- and A/V-type ATP synthases, helicases of superfamilies 4 and 5 (SF4 and SF5), as well as several other protein families [3, 107, 121]. In NTPases of this class, similarly to most AAA+ ATPases, the stimulatory moiety(is) is/are inserted in the catalytic site by the P-loop domain of the adjacent monomer (Fig. 6, S4C).

**Figure 6.**
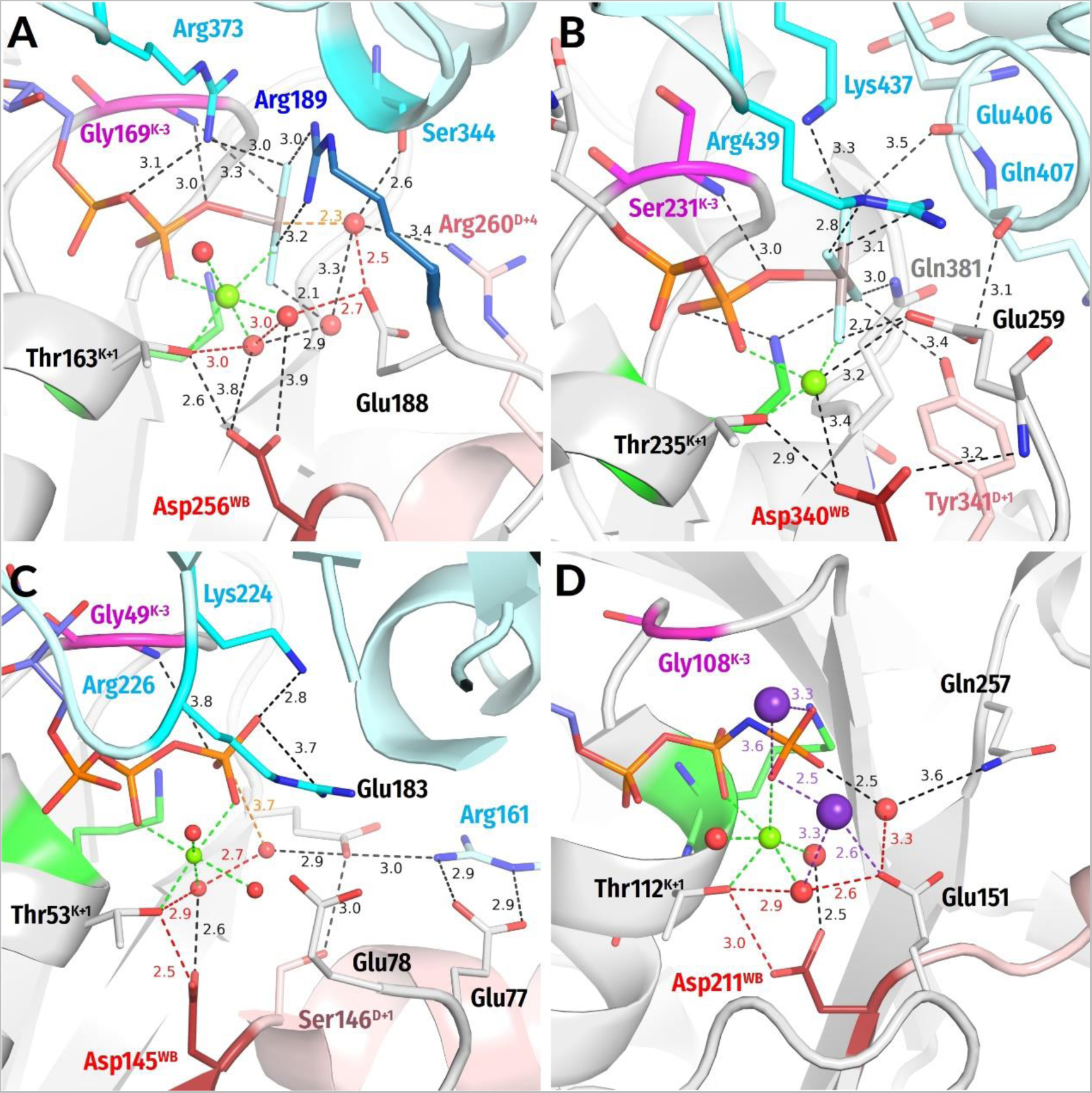
Representative proteins of the RecA/F_1_-like class of the P-loop NTPases. The adjacent monomers and their Arg and Lys residues are shown in cyan, K^+^ ions are shown as purple spheres. The Arg residue from the P-loop domain proper is shown in deep blue (panel A). Red dashed lines mark protonic connections between W_cat_ and Asp^WB^. Other colors as in Fig. 3. All distances are in ångströms. **A,** Bovine F_1_-ATPase (PDB ID 1H8E, [54]); **B,** DnaB replicative helicase from *Vibrio cholerae* (PDB 6T66 [123]); **C**, circadian clock protein KaiC (PDB ID 4TL7, [124]); **D,** RadA recombinase (PDB ID 3EW9, [125]).

Our structure analysis showed that the stimulation mechanism appears to differ between rotary F_1_-ATPases/SF5 helicases, on the one hand, and recombinases/SF4 helicases, on the other hand.

In the ***F_1_-ATPases***, the stimulatory Arg residue of the adjacent monomer interacts with both α- and γ-phosphates, whereas an additional, intrinsic Arg residue coordinates γ-phosphate, approaching it apically (see Fig. 6A, S4C and Table 1). A similar stimulation mechanism is realized in ***SF5 helicases***, see e.g. the Rho helicase (PDB ID 6DUQ [122]).

The W_cat_ molecule is seen in the catalytic apical position only in the ADP:AlF_4_^-^ -containing structure of the bovine F_1_-ATPase (PDB ID 1H8E, Fig. 6A) [54]. While in most ASCE ATPases the W_cat_-coordinating Glu^D+1^ residue directly follows the Asp^WB^ of the Walker B motif, the W_cat_-coordinating Glu188 of the F_1_-ATPase, similarly to SIMIBI NTPases, is at the C-cap of the WB+1 strand; within the ASCE division this feature is specific for the RecA/F_1_ class members [8]. This glutamate residue also links W_cat_ with the Mg^2+^-coordinating W3 molecule (Fig. 6A). In F_1_-ATPase, W_cat_ is also stabilized by Arg260^D+4^; this residue concurrently interacts with the activating neighboring monomer, which also provides a **CO** group to stabilize W_cat_ (Fig. 6A).

In ***RecA-like recombinases*** and ***SF4 helicases***, two stimulatory positively charged moieties interact only with the γ-phosphate group, in contrast to F_1_-ATPases. In several families, the adjacent subunit provides one Lys residue and one Arg residue that form a short KxR motif [126], with both residues reaching only γ-phosphate. These are, for instance, the bacterial helicase DnaB (Fig. 6B, see also [127]), circadian clock protein KaiC (Fig. 6C [124]), and gp4d helicase from the T7 bacteriophage (PDB ID 1E0J, [128]). In bacterial RecA recombinases, the adjacent monomer in the homooligomer provides two Lys residues which approach the phosphate chain laterally and interact with γ-phosphate as in the RNA recombinase of *E. coli* (see Table 1 and PDB ID 3CMX, [129]). In archaeal and eukaryotic RadA/Rad51-like recombinases, the positions of terminal groups of stimulatory Lys/Arg residues are occupied by two K^+^ ions, which appear to interact with γ-phosphate, see Fig. 6D and [126, 130].

In the case of RecA-like ATPases, the AlF_4_^-^-containing structures of the bacterial DnaB helicase of *Vibrio cholerae* (Fig. 6B), *Bacillus stearothermophilus* (PDB ID 4ESV, [131]) and RecA recombinase of *E. coli* (PDB ID 3CMX, [129]) are available. However, the apical water molecule is absent from these structures, the possible reasons of which are discussed in [127]. Therefore, the full set of W_cat_-coordinating groups remains unknown for RecA-like ATPases; the available structures imply the participation of glutamate residues at the C-terminus of the WB+1 strand (Fig. 6B-D), as in other RecA/F_1_ class enzymes. The same residue appears to link the would-be W_cat_ with the Mg^2+^-coordinating W3 molecule (Fig. 6B-D).

Generally, the H-bond networks around γ-phosphate and W_cat_ seem to be richer in RecA/F_1_-NTPases than in other classes of P-loop NTPases. Specifically, the coordination of W_cat_ may potentially also involve the D+1 residue of the WB strand, which is usually a H-bonding Ser/Thr/Asn/Tyr, as well as other polar residues at the C-caps of the WB-1 and WB+1 strands [112, 124, 132].

#### 2.2.3. Water molecules in the vicinity of the Mg^2+^ coordination shell

The W_cat_ molecule should be deprotonated on some stage of the catalytic cycle, see Fig. S1 and [15, 48, 56–59, 61–63, 133]. However, the pK_a_ value (hereafter pK for simplicity) of water is 14.0, which hinders the deprotonation both thermodynamically and kinetically.

The P-loop NTPases, in general, are not alone in their need to deprotonate a water molecule and to use the resulting hydroxyl for hydrolyzing the O-P bond. In various other, evolutionarily unrelated Mg-dependent enzymes, Mg^2+^ ions are thought to decrease the pK value of potential nucleophilic water molecules [60, 134–137]. In such cases, usually, the would-be nucleophilic water molecule belongs to the first or second coordination shell of the metal cation. In the case of P-loop NTPases, the Mg^2+^ ion is remote from W_cat_ and can incite its deprotonation only if there is a protonic pathway between W_cat_ and the coordination shell of Mg^2+^. Therefore, we manually searched for connections between W_cat_ (or the would-be W_cat_) and the Mg^2+^-coordinating ligands.

In many high-resolution structures of P-loop NTPases with bound substrate molecules or their analogs, water molecules are present in the vicinity of γ-phosphate and interact with the O^1G^, O^2G^, O^3G^ atoms of γ-phosphate, as seen in Fig. 7. In principle, a water molecule can interact with up to two oxygen atoms, which allows categorizing the γ-phosphate-bound water molecules as, for example, W_12_, W_13_, or W_23_. Most likely, one of these molecules becomes the W_cat_ upon activation. For instance, Fig. 7A shows a structure of RadA from *Pyrococcus furiosus* that was post-soaked with ATP, see PDB ID 4A6X and [138]. The structure clearly shows W_12_ and W_13_ and resolves two conformations of the “catalytic” Glu174 of the WB+1 strand. In both conformations, Glu174 interacts with W_13_, the apparent would-be W_cat_ in this enzyme. However, only in the Glu174A conformation, the side chain of Glu174 connects W_cat_ with the Mg^2+^-coordinating W3 molecule.

**Figure 7.**
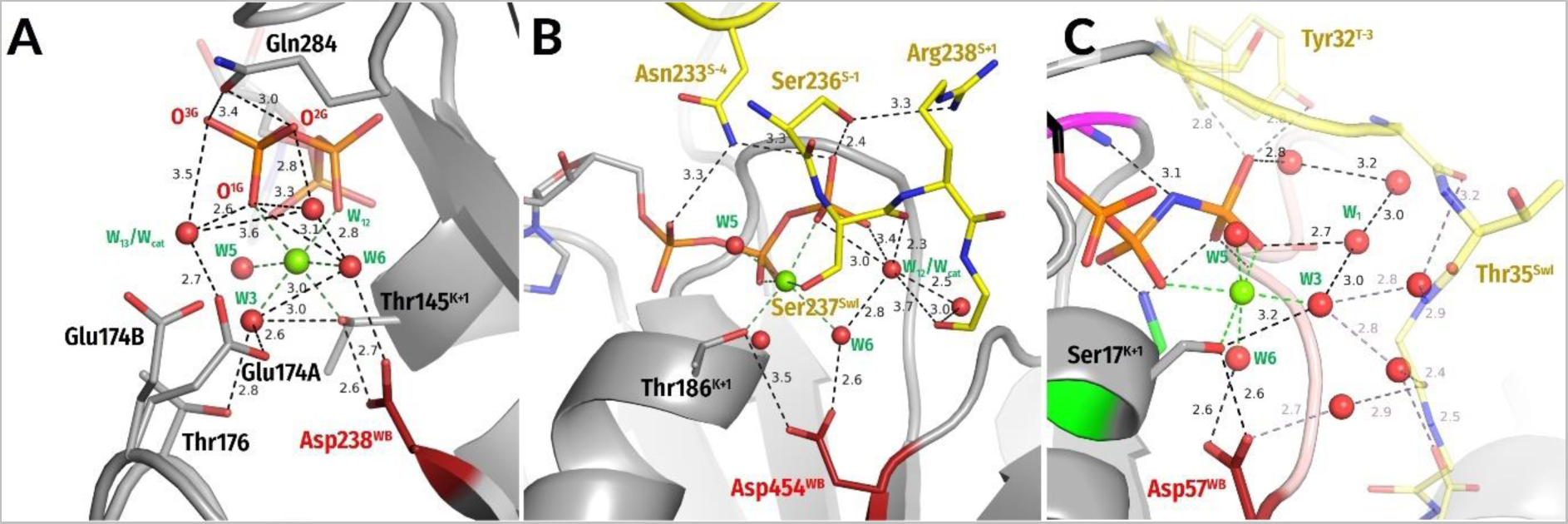
Networks of water molecules in the catalytic sites of P-loop NTPases. **A**, ATP-soaked ATPase domain of RadA from *Pyrococcus furiosus* (PDB ID 4A6X, [138]); **B**, ATP-soaked Myosin II from *Dictyostelium discoideum* (PDB ID 1FMW, [139]); C, human H-Ras GTPase (PDB ID 5B30 [140]). Colors as in figures 3-6.

Similarly, the “catalytic” Glu or Asp residues link W_cat_ with Mg^2+^-ligands #3 or #6 in TS-like structures of NTPases of all classes (except the TRAFAC class), as already repeatedly noted throughout the Section 2.2. The tentative proton pathways from W_cat_ towards the Mg^2+^-ligands are shown by red dashed lines in Fig. 4A-B, 4D, 5A-D, 6A, 6C-D.

In the case of TRAFAC class NTPases, which lack “catalytic” Glu/Asp residues, connections between W_cat_ and the Mg^2+^-coordinating ligands are not so evident. The water molecules around γ-phosphate appear to be highly mobile and hardly crystallizable. In the absence of TS-analogs, those water molecules that interact with γ-phosphate are usually resolvable in the structures only if constrained by additional interactions. For instance, in the ATP-soaked myosin structure (PDB ID 1FMW [139], see Fig. 7B), W_12_ is resolved because it is additionally bound to **CO** of Ser^S-1^. This W_12_ molecule is at a H-bond-compatible distance from the Mg^2+^-coordinating W6 molecule.

In TS-like structures of small GTPases of the TRAFAC class, the closest to W_cat_ Mg^2+^-coordinating ligand is [Ser/Thr]^SwI^ of Switch I at about 4-5 Å (Fig. 1E, 3A-D). In these proteins, however, the Switch I motif is very dynamic so that the protein fluctuates between “open” and “closed” conformations even with a bound substrate or its analogs [140–142]. While in the closed conformation [Ser/Thr]^SwI^ serves as ligand #3 for Mg^2+^, in the open conformation the whole Switch I moves away from Mg^2+^ and a water molecule serves as ligand #3 (Fig. 7C, S5B). Notably, Matsumoto and colleagues succeeded to control the open to close transition in H-Ras GTPases by changing the humidity [140]. The crystal structure of H-Ras that crystalized in the open conformation at low humidity (PDB ID 5B30) reveal an additional water molecule next to W3 (Fig. 7C). It is depicted as W_1_ in Fig. 7C and occupies approx. the same position as

W_12_ in the myosin structure in Fig. 7B. In the closed conformation (Fig. 1E, 3A-D), Switch I reaches Mg^2+^ and ousts W_1_ and W3; only W_cat_ is present in the TS-like structure (cf Fig. 1E and 7C). As argued in section 3.2.2.4 of the Discussion, one of the two waters seen in the “open” structure may turn into W_cat_ whereas the other may transiently form a proton pathway between would-be W_cat_ and the Mg^2+^-coordinating [Ser/Thr]^SwI^.

#### 2.2.4. Local molecular electric field in P-loop NTPases

In structures of P-loop NTPases that are shown in Fig. 1-7, the positive charged groups that stabilize the triphosphate chain, namely the Lys^WA^ residue, the backbone **HN** groups, the stimulatory moiety(ies), the auxiliary residues, and the positively charged N-terminus of the first α-helix are opposed by acidic residues that interact with W_cat_ either directly or via water bridges. Together with Asp^WB^, these catalytic acidic residues form negatively charged clusters. Fig. 8 shows, for diverse P-loop NTPases, these positively and negatively charged clusters, which should produce a strong local electric field, as argued in the Discussion section.

**Figure 8.**
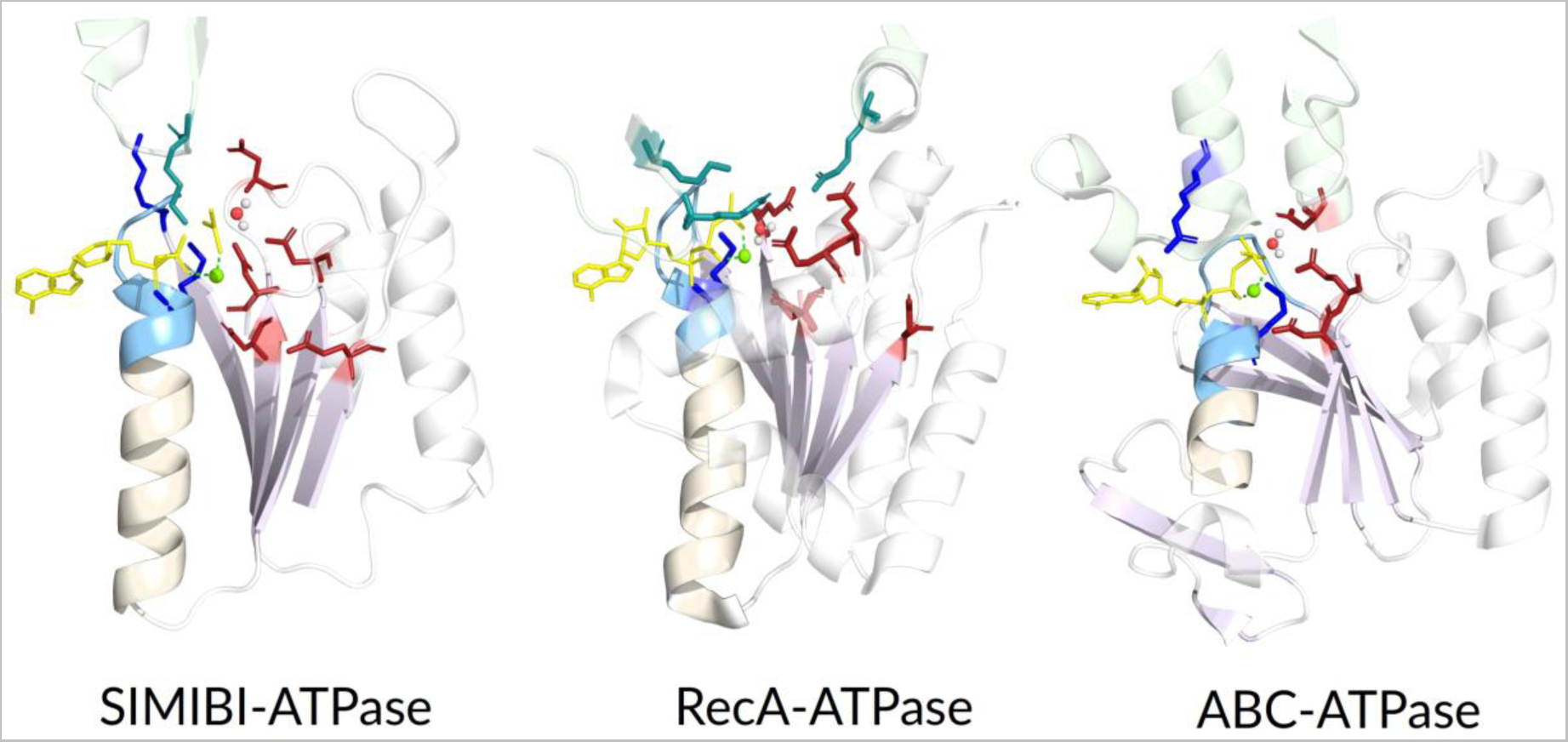
Uneven distribution of charged residues around nucleotide binding pockets in different classes of P-loop NTPases. Nucleotide analogs are shown in yellow, negatively charged residues are shown in red, positively charged residues are shown in blue and teal, the α_1_ helix is shown in beige with its N-terminus and the P-loop shown in light blue. **A**, SIMIBI class, light-independent protochlorophyllide reductase iron-sulfur ATP-binding protein chlL) (PDB ID 2YNM, [99]); **B**, RecA class, circadian clock protein kinase KaiC (PDB ID 4TL7, [124]); **C**, ABC-ATPase class, MBP-maltose transporter complex (PDB ID 3PUW,[116]).

### 2.3. Global structural analysis of hydrogen bonding between the Walker A and Walker B motifs in the whole set of P-loop NTPases with bound Mg-NTP complexes or their analogs

As described in the Methods section as well as in the in the accompanying article [41], we extracted from the Protein Data Bank (PDB) at www.rcsb.org [71, 72] those structures of P-loop NTPases that have substrates or their analogs bound together with a Mg^2+^ ion, yielding as many as 1383 structure entries with 3136 full-fledged catalytic sites (as of 11.0.2019; many of the structures contained several catalytic sites). The criteria for selection of full-fledged catalytic sites and the routine of their subsequent structural analysis are described and depicted in the Methods section of the accompanying article [41]. As discussed in Section 2.2 and depicted in Fig. 2, the shape of the P-loop and the spatial position of Asp^WB^ relatively to it are strictly conserved among P-loop NTPases. Building on that, we were able, here and in the accompanying article [41], to use the atomic coordinates of the Mg-triphosphate to pinpoint the catalytically relevant residues in an almost sequence-agnostic way, as described in the Methods sections of both articles. We applied this approach to perform analysis of nucleotide-binding sites in thousands of structures of P-loop NTPases. Relevant data on all these catalytic sites are cumulated in Supplementary Table 1 (hereafter Table S1).

Based on the type of the molecule bound, the catalytic sites could be sorted into four groups: 1043 sites contained native ATP/GTP molecules; 1612 sites contained bound non-hydrolyzable NTP analogs such as Adenosine 5′-[β,γ-imido]triphosphate, (AMP-PNP), Guanosine 5′-[β,γ-imido]triphosphate (GMP-PNP), Adenosine 5′-[*β*,*γ*-methylene]triphosphate (AMP-PCP), Guanosine 5′-[*β*,*γ*-methylene]triphosphate (GMP-PCP), Adenosine 5′-[γ-thio]triphosphate (ATP-γ-S), and Guanosine 5′-[γ-thio]triphosphate (GTP-γ-S); 234 sites contained NDP:fluoride complexes mimicking the substrate state, such as NDP:BeF_3_ and NDP:AlF_3_, and 247 sites were with TS analogs NDP:AlF_4_^-^(204), NDP:MgF_3_^-^ (10) and ADP:VO_4_^-^ (33).

#### 2.3.1. The H-bond between Asp^WB^ and [Ser/Thr]^K+1^ is shorter in the presence of transition state analogs

The H-bond between Asp^WB^ and [Ser/Thr]^K+1^ is special because it links the Walker A and Walker B motifs. Consequently, the length of this bond is a good measure of the catalytic site constriction. Upon our manual analysis, we noted that this H-bond is particularly short in the NDP:AlF_4_^-^-containing representative structures (see Fig. 1-6). Therefore, we measured the distances between the two residues to characterize their H-bonding in all the available structures of P-loop NTPases with bound substrates or their analogs, as described in the Methods section. The results of these measurements for all the analyzed structures are presented in Table S1. Fig. 9 summarizes the data on best resolved structures (with resolution of 2.5 Å or less). The data for the three TS analogs are plotted separately for the sake of clarity. We consider as short the H-bonds of < 2.7 Å [143]; the range of short H-bonds is highlighted by cyan. As seen in Fig. 9, the length of the H-bond between Asp^WB^ and [Ser/Thr]^K+1^ is, on average, < 2.7 Å in all groups of structures and < 2.6 Å in structures with ATP/GTP, nonhydrolyzable NTP analogs and NDP:MgF_3_^−^ or NDP:AlF_4_^-^ as TS analogs. Specifically, in NDP:AlF_4_^−^-containing structures the H-bond distance is on average as short as 2.5 Å. It is noteworthy that improper binding of AlF_4_^-^ can distort the coordination sphere of Mg^2+^, as documented in the accompanying article [41]. Therefore, for the plot in Fig. 9, we selected only the sites with a correct Mg^2+^ coordination; the cut-off for the distance between Mg^2+^ and [Ser/Thr]^K+1^ was 2.5 Å. When comparing only TS analogs, fluoride complexes were superior to vanadate complexes in their ability to constrict the catalytic site, as can be judged from the distributions of the Asp^WB^–[Ser/Thr]^K+1^ bond lengths (Fig. 9). From the same data it follows that the NDP:AlF_4_^-^ complexes “overperform” other metal fluorides complexes in the ability to constrict the catalytic pocket.

**Figure 9.**
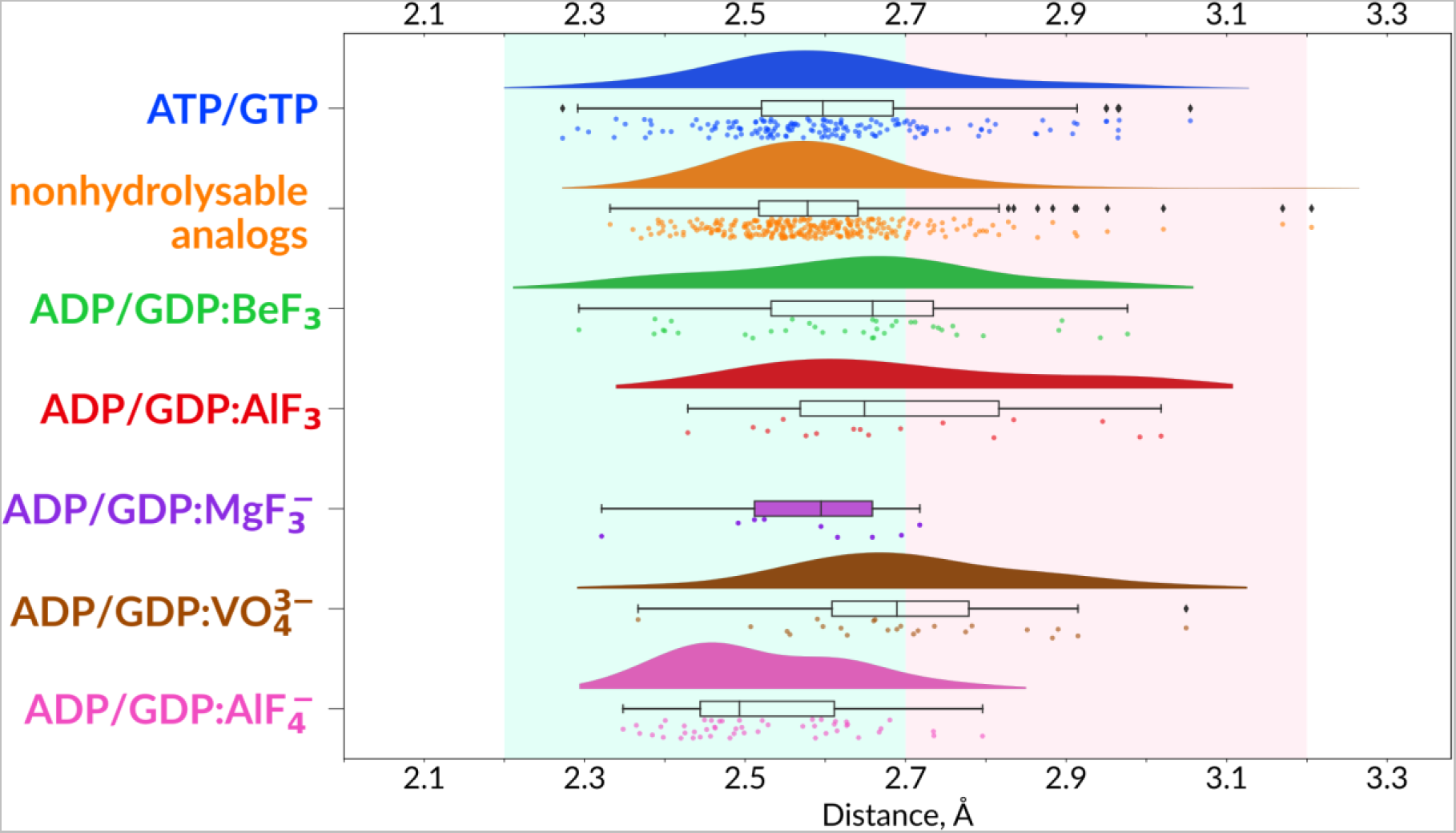
Distances between the side chains of [Asp]^WB^ and [Ser/Thr]^K+1^. For each type of complexes, distances are visualized as a kernel density estimate (KDE) plot, a boxplot and individual data points, each point representing one catalytic site in one structure. For ADP/GDP:MgF_3_^-^ complexes the density plot is not shown owing to the scarcity of data. Data are shown only for structures with resolution ≤ 2.5 Å (see Table S1 for the whole data set). The integrity of catalytic sites was checked by two criteria: the [Ser/Thr]^K+1^ – Mg^2+^ distance ≤ 2.5 Å and Mg^2+^ – Asp^WB^ distance ≤ 6 Å. The range of short H-bonds (2.2 Å-2.7 Å) is highlighted in cyan, the typical H-bond range (2.7 Å-3.2 Å) is highlighted in pale pink.

#### 2.3.2. The TS analog AlF_4_^-^ makes more bonds within the catalytic site than MgF_3_^-^ or AlF_3_

We hypothesized that the unique ability of AlF_4_^-^ to pull the catalytic site together may correlate with its ability to enter more bonds than MgF_3_^-^, AlF_3_ or BeF_3_. We tested this hypothesis by comparing the structures of the same activated complexes with bound NDP:AlF_4_^-^ and NDP:MgF_3_^-^/AlF_3_, respectively. We counted and measured all the bonds in catalytic sites for the two pairs of such structures. One pair of structures consisted of RhoA GTPases in complex with their activator RhoGAP and GDP:AlF_4_^-^ (PDB ID 1TX4, resolution 1.65 Å [144]) or GDP:MgF_3_^-^ (PDB ID 1OW3, resolution 1.8 Å [34]), respectively, bound. The other pair was made up of catalytic sites of bovine mitochondrial F_1_-ATPase with bound ADP:AlF_4_^-^ (PDB ID 1H8E, resolution 2.0 Å [54]) or ADP:AlF_3_ (PDB ID 1E1R, resolution 2.5 Å [70]), respectively. In the latter case, the fluoride derivative can be identified as a genuine ADP:AlF_3_ by the similarity of its geometry to that of the Mn-ADP:AlF_3_ complex in the Zika virus helicase (PDB ID 5Y6M [145]), see the accompanying paper [41] for details.

The findings are summarized in Supplementary Table S2 (hereafter Table S2), which shows that AlF_4_^-^ forms more bonds with surrounding residues than MgF_3_^-^ or AlF_3_. Specifically, AlF_4_^-^ usually makes two H-bonds, one moderate and one weak, with Lys^WA^, whereas MgF_3_^-^ or AlF_3_ make only one bond. In addition, AlF_4_^-^ makes two H-bonds with **HN**^K-3^, whereas MgF_3_^-^ or AlF_3_ make only one (cf Table S1 and S2). Furthermore, many of the AlF_4_^-^ bonds are shorter than those of MgF_3_^-^, see Fig. 9 and Tables S1 and S2. Apparently, the NDP:AlF_4_^-^ complexes have more potency to constrict he catalytic site than other TS analogs.

## 3. Discussion

Here and in the accompanying article [41], we reported the results of a comparative structural analysis of more than 3136 catalytic sites of P-loop NTPases with nucleoside triphosphates or their analogs bound. The aim of the analysis was to deduce the main structural features common to the catalytic sites of P-loop NTPases and use them to elucidate the mechanism(s) of stimulated NTP hydrolysis.

In our work, we used the tools of evolutionary biophysics, which assumes that certain structural elements are conserved by evolution because of their functional importance and that the degree of conservation of the structural elements involved should be taken into account when choosing from several possible biophysical mechanisms consistent with the structural data, see [126, 146–150] for some examples of this approach. In the case of P-loop NTPases, the *in silico* approach is particularly justified since the experimental investigation of all members of this vast superfamily would be unrealistic. In addition, P-loop NTPases are unique in that their catalytic repertoire is unusually narrow for such a vast superfamily [11], which increases the chances of finding a common catalytic mechanism.

### 3.1. Structure comparison of P-loop NTPases

#### 3.1.1. Constriction of the catalytic site in the transition state

The data presented here, as well as data from the accompanying article [41], show that binding of TS analogs leads to greater constriction of catalytic sites than binding of ATP, GTP, or their analogs. In some cases, this constriction can be revealed by comparing the structures of same P-loop NTPases with bound substrate/substrate analogs and TS-analogs, respectively (Fig. S5). In the presence of a TS analog, the WB-crest moves closer to γ-phosphate (Fig. S5). Furthermore, the Switch I of TRAFAC class NTPases undergoes dramatical conformational changes (see Fig. S5B, cf Fig 1E and 7C).

The constriction also manifests itself in the convergence of the Walker A and Walker B motifs. As it follows from Fig. 9 and Table S1, the most H-bonds between the side chains of Asp^WB^ and [Ser/Thr]^K+1^ residues can be categorized as short in those structures that contain ATP/GTP or their non-hydrolyzable analogs. Apparently, these short H-bonds form upon “closing” of the catalytic site in response to the substrate binding. Still, these H-bonds are even shorter in the presence of NTP:AlF_4_^-^ and NTP:MgF_3_^-^ (Fig. 9, Table S1), which indicates a further constriction of the catalytic pocket in the presence of these TS analogs.

Noteworthy, our structural analysis initially revealed Asp^WB^ – [Ser/Thr]^K+1^ distances of ≥ 4.0 Å in several NTP:AlF_4_^−^-containing structures (see Table S1). Further inspection of these structures, however, has shown that AlF_4_^−^ moiety is “improperly” bound; it makes two bonds with Mg^2+^ in all such structures but one (Table S1). When AlF_4_^−^ makes two bonds with Mg^2+^, the bond between Mg^2+^ and [Ser/Thr]^K+1^ can get lost and the whole catalytic site undergoes distortion, as detailed and illustrated in the accompanying article [41]. In these cases, (all marked pink in Table S1), the structure of the catalytic site cannot be considered as analogous to the TS.

After we excluded these cases, the Asp^WB^ – [Ser/Thr]^K+1^ distances found to be H-bond compatible in all TS-analog-containing structures (about 200) but one, which is depicted in Fig. S6 and discussed in its caption (Table S1). Hence, AlF_4_^−^-containing P-loop NTPases can be used as reliable models of transition states, in support of earlier suggestions [48, 53, 56, 151], provided that the “properness” of the interaction between AlF_4_^−^ and Mg^2+^ is checked in each particular case.

Earlier, our MD simulations of the MnmE GTPase showed that γ-phosphate, after its twisting by the stimulatory K^+^ ion, was stabilized by a new H-bond between NH^K-3^ and O^2G^ [40]. Our structural analysis, as presented in the accompanying article [41], strongly indicates that the counter-clockwise twist of γ-phosphate correlates with the formation of a new H-bond with O^2G^ indeed. Specifically, H-bond-compatible distances between **HN**^K-3^ and the nearest oxygen/fluorine atom are seen in most structures with TS analogs bound, see Table S1 and the accompanying paper [41].

Twisted γ-phosphates and H-bond compatible **HN**^K-3^ - O^2G^ distances are also seen in diverse NTP-containing NTPases which were crystalized in their pre-transition configurations because of mutated W_cat_-coordinating residues and/or their impaired interaction with W_cat_, see Table S1 and Fig. 3 in the accompanying paper [41].

In sum, as it follows from Fig. 9, S5, Table S1, and the data in the accompanying article [41], the structures with TS analogs bound have particularly short H-bonds between **HN**^K-3^ and O^2G^, as well as between Asp^WB^ and [Ser/Thr]^K+1^, which suggests an additional constriction of the catalytic site in the TS.

#### 3.1.2. Transition state analogs and energetics of P-loop NTPases

The observation of the shortest H-bonds within the catalytic sites in the presence of NDP:AlF_4_^-^ (see Fig. 9, Table S1 and S2) helps to clarify why these complexes are most potent functional TS-mimics in P-loop NTPases [34, 42, 43, 48, 49, 53, 54, 56, 90, 151–160]. Beginning in 1987, that is, even before the first GDP:AlF_4_^-^-containing structures were resolved in 1994 [42, 43], metal fluorides were shown to promote binding of various P-loop NTPases to their activating partners. Generally, many NTPases interacted with their activators only in the presence of metal fluorides [48, 49, 53, 56, 90, 151–154, 156, 161, 162]. Crystal structures of P-loop NTPases with NDP:AlF_4_^−^ or NDP:AlF_3_/NDP:MgF_3_^−^ complexes bound [34, 42, 43, 48, 54, 56, 90, 155, 160] show that metal-fluorides have planar (AlF_4_^-^ and MgF_3-_) or almost planar (NDP:AlF_3_) structures with W_cat_ in the apical attack position, see respective panels in Fig. 1-6 and the accompanying paper [41]. While the shape and electric charge of MgF_3_^-^ is the same as that of γ-phosphate in the anticipated TS (see Fig. S1), the shape of AlF_4_^-^ differs significantly, it is tetragonal instead of trigonal. Nevertheless, NDP:AlF_4_^-^ complexes are more potent functional analogs of the TS than NDP:MgF_3_^-^ or NDP:AlF_3_, as follows from their superiority in promoting binding of P-loop NTPases to their activators [48, 49, 53, 56, 90, 151–154, 156, 161, 162]. In addition, the distances between W_cat_ and the metal atom are shorter (2.0-2.1 Å) in the case of NDP:AlF_4_^-^ complexes than NDP:MgF_3_^-^ (approx. 2.5 Å) or NDP:AlF_3_ complexes (approx. 3.0 Å) see the accompanying article [41] and [48, 56]. The reason why NDP:AlF_4_^-^ performs better than other substrate and TS analogs in all these cases has remained obscure, see e.g. [151].

The reason could be, however, understood by considering the energetics of P-loop NTPases. Our earlier MD simulations have shown that the linking of α- and γ-phosphates by an inserted stimulatory cationic moiety is an endergonic reaction; about 20-25 kJ/mol appear to be needed to twist the γ-phosphate by straining its bonds with Mg^2+^ and Lys^WA^ [40, 163]. This activation barrier prevents haphazard NTP hydrolysis due to accidental insertion of a K^+^ ion or an Arg residue in the catalytic site, as argued elsewhere [40, 163]. These estimates from MD simulations corroborate earlier considerations of Warshel and colleagues that the stimulation of hydrolysis in P-loop NTPases by their cognate activators requires an input of about 20-25 kJ/mol of free energy, see e.g. [65, 133].

A specific problem of P-loop NTPases is the source of a such free energy input. In most enzymes, the free energy for lowering the activation barrier – by destabilizing the substrate-bound ground state and/or stabilizing the TS – is usually provided by substrate binding [164]. In P-loop NTPases, where the substrate binding step is separated from the ultimate catalytic step in most cases, the energy of substrate binding is partly used to bring the NTP molecule into its extended, catalytically prone conformation with eclipsed β- and γ-phosphates [36, 38, 165]. Another part of the binding energy is used to pre-organize the catalytic site, with the conformational changes that accompany catalytic site closure often being coupled with useful mechanical work, e.g. in myosins [92, 166]. While increasing the rate of NTP hydrolysis as compared to that in water (by five orders of magnitude in the case of Ras GTPase [39, 167]), the free energy of NTP binding to the P-loop is spent without achieving physiologically relevant hydrolysis rates. For the further acceleration of hydrolysis, additional source(s) of free energy is/are needed.

The source of additional free energy is evident in the case of ring-forming oligomeric P-loop NTPases, such as many AAA+ ATPases (see Fig. 5A and [109, 168]), helicases (see Fig. 5B-C and [14, 112]), or rotary ATPases/synthases of the RecA/F_1_ class (see Fig. 6A and [169–171]). In such complexes, the free energy for the cleavage of a bound ATP molecule in one monomer is provided by the binding of another ATP molecule to another monomer. Part of the substrate binding energy at this other site drives a molecular motion that is transferred, for example, to the Arg finger; in the case of rotational F_1_-ATPases, Paul Boyer called this kind of coupling “binding change mechanism” [171].

The same reasoning can be applied to ABC transporters (Fig. 5D), where ATP-hydrolysis appears to be coupled to the binding of the translocated molecule. Also, the activation of kinases (Fig. 4C-D) appears to be driven by binding of their phosphate-accepting substrates.

In other cases, however, P-loop NTPases, with an NTP molecule bound, do not bind further small molecules, but only the cognate activating partners, which are protein/RNA/DNA molecules (Table 1). These NTPases, per exclusionem, can harness only the free energy of binding to their activating partner(s). Indeed, there is evidence that the strength of such protein-protein binding does affect the catalytic activity of some P-loop NTPases [4, 172–174], making them easy to regulate via protein-protein interactions, as observed with small GTPases.

In all these cases, we are dealing with thermodynamic coupling where the free energy of binding is used for endergonic constriction of the catalytic site. Such a thermodynamic coupling is reversible by definition. Consequently, if we find a way to exergonically constrict the catalytic site, it will lead to the spontaneous assembly of the complex between P-loop NTPase and its cognate activator(s). Apparently, the binding of TS analogs such as NDP:AlF_4_^-^ or NDP:MgF_3_^-^, which form multiple strong bonds within the catalytic site, is an exergonic reaction that leads to a TS-like configuration of the catalytic site by “constricting it from the inside”, which can happen even in the absence of an activating partner (as in the NDP:AlF_4_^-^- containing separately crystallized dimer of GTPase domains of the MnmE protein (Fig. 1F in the accompanying paper [41]) or SIMIBI NTPases (Fig. 4A-B). If cognate activators are also present, the “from-the-inside constriction” drives their interaction with the NTPase domain yielding a full-fledged activated complex, as described in the literature [48, 49, 53, 56, 90, 151–154, 156, 161, 162].

If this assumption is correct, then the assembling “efficiency” of the TS analogs will depend on the amount of free energy that becomes available upon their binding. As it follows from Fig. 9 and Fig. 2 of the accompanying paper [41], fluoride complexes are superior to vanadate complexes in their ability to constrict the catalytic site. This is consistent with the fact that the more electronegative fluorine atoms form stronger H-bonds than oxygen atoms. Also NDP:AlF_4_^-^ complexes are superior to NDP:MgF_3_^-^ complexes (Fig. 9 and Fig. 2 of the accompanying paper [41]) because, as documented in Table S2, they enter into more bonds. Not surprisingly, NDP:AlF_4_^-^ complexes, which “raise” more binding energy despite their “unphysiological” geometry, are the most efficient in assembling activated complexes *in vitro*.

In sum, the extent of H-bonding of TS analogs within catalytic sites of P-loop NTPases (Fig. 9, Tables S1, S2 and Fig. 2 of the accompanying paper [41]) appear to correlate with their ability to induce self-assembly of activated complexes [48, 49, 53, 56, 90, 151–154, 156, 161, 162], which implies that these H-bonds stabilize the TS upon hydrolysis of ATP or GTP.

#### 3.1.3. Coordination of W_cat_ and auxiliary interactions

The W_cat_ molecule is usually stabilized by the combined action of evolutionary conserved and variable moieties. The conserved part is represented by residue(s) of the WB-crest and contains residues that bind either W_cat_ alone or W_cat_ and γ-phosphate, such as Gly^D+3^ and Gln^D+4^ of small GTPases (Fig. 3), or Asp^D+2^ of many kinases (Fig. 4 C-D), or Glu^D+1^ of many ASCE ATPases (Fig. 5). Variable residues are provided by the C-caps of the β-strands, family-specific protein loops, or even external activating proteins/domains, as documented in Table S1.

In the Kinase-GTPase division, not only the position but also the nature of W_cat_-coordinating residues varies between classes and families (Fig. 1E, 3, 4, S2); specifically, residues and/or **HN** groups in different positions of WB-crest are involved (Fig. 1E-F, 3, 4, Table 1). In most members of the ASCE division, W_cat_ interacts with the “catalytic” Glu^D+1^ residue of WB-crest (Fig. 5, Table 1). In Rec/F1 ATPases, the “catalytic” Glu residue is provided by the WB+1 strand and complemented by varying residues of WB-crest (Fig. 6, Table 1).

In addition, polar, W_cat_-stabilizing ligands may be provided by the activating partners. In some Ras-like NTPases, GAPs can provide both the Arg finger and the W_cat_-stabilizing Gln [175]. In SIMIBI NTPases, the “activating” monomer of the dimer provides not only the Lys finger, but also the Asp or a Glu residue that stabilizes W_cat_ (Fig. 4A-B). In maltose transporter ABC ATPase (Fig. 5D) and F_1_-ATPases (Fig.6A), the adjacent monomer that provides the stimulator also provides the backbone **CO** atom to coordinate W_cat_ (Fig. 5D, 6A). Such behavior, where both the stimulator and the W_cat_-stabilizing residue(s) are controlled by the interaction with the activating partner, might represent a kind of a “two-key mechanism” to better control and prevent accidental unwanted NTP hydrolysis.

In most cases, other auxiliary positively charged groups, such as **HN** groups and/or additional Arg/Lys residues, are involved in the coordination of the oxygen atoms of γ-phosphate and/or W_cat_ in addition to the “main” stimulator. This is observed in AAA+ NTPases (Fig. 5A), many helicases (Fig. 1F, 5B, 5C), ABC NTPases (Fig. 5D), hexamers of F_1_/RecA-like ATPase (Fig. 6), PilT-like proteins (Fig. SF1_2A in the supplementary File 1), see also the accompanying article [41]. These auxiliary residues, indicated for the various classes of P-loop NTPases in Tables 1 and S1, are poorly conserved even within individual families of P-loop NTPases.

The abundance and variety of such auxiliary interactions could be related to the energetics of P-loop NTPases, which was discussed in section 3.1.3. Many of H-bonding interactions between oppositely charged moieties of the P-loop domain and its activator, which often invoke residues of WB-crest, may provide the free energy needed for catalysis. If so, elimination – for example through mutations – of such interactions, even if they are not involved in the chemistry of catalysis, would still slow the enzyme reaction, which is consistent with some experimental observations [4, 49, 172]. Therefore, a case-by-case analysis is needed to understand whether specific auxiliary residues contribute directly to catalysis or only to the exergonic interaction of the NTPase domain with its activator.

#### 3.1.4. Common structural traits of P-loop NTPases

Comparative structural analyses, as performed here, in the accompanying article [41], and in our earlier paper [40], unraveled the following common structural features of P-loop NTPases:

i. in P-loop NTPases of different classes, the NTP molecule (or its analog) is bound by the Walker A motif in a same extended conformation (Fig. 2A); this conformation is likely to be catalytically prone [40];
ii. our structure analyses showed, in agreement with previous data (see e.g. [50, 176]) that the activating partner usually binds to the P-loop domain by making new H-bonds and salt bridges with the residues of the WB-crest, see Fig. 1C-D, S5. In all inspected structures for which TS-like structures are available, some of the amino acid residues that immediately follow Asp^WB^ (at the D+1 – D+5 positions of the WB-crest) are involved in catalytic interactions with W_cat_ (Fig. 1-7, S5, Table 1 and Table S1). The energy of binding of the activation partner seems to be used for pushing W_cat_ towards the P^G^ atom of γ-phosphate, constricting the catalytic site, and inserting the stimulatory moiety;
iii. in the accompanying article [41], we provide evidence that the common trait of all inspected stimulators is their mechanistic interaction with the oxygen atom(s) of γ-phosphate;
iv. comparing the structures with analogs of ATP/GTP and TS, respectively, we noticed that the binding of TS-analogues results in greater constriction of catalytic sites than the binding of ATP or GTP. The constriction manifests itself in the shorter distances between Asp^WB^ and [Ser/Thr]^K+1^ which are as short as 2.5 Å on average in the presence of ADP:AlF_4_^-^ (Fig. 1-6, 9). Also the distances between **HN**^K-3^ and the analogs of γ-phosphate are shorter in the TS-like structures, see Fig. 1-6 and the accompanying article [41].
v. In TS-like configurations of P-loop NTPases of all major classes, except the TRAFAC NTPases, the W_cat_-coordinating “catalytic” Glu or Asp residue links W_cat_ with Mg^2+^-ligands in positions 3 and/or 6, see Fig. 1E, 1F, 4-6, 7A.

These common structural traits, which are shared by P-loop NTPases of different classes, have been mostly overlooked so far. We believe that these features are of key importance for understanding the common mechanism of P-loop NTPases.

### 3.2. Common mechanistic traits in P-loop NTPases

#### 3.2.1. Background on the catalysis by P-loop NTPases

The cleavage of γ-phosphate by P-loop NTPases is thought to involve a nucleophilic attack of W_cat_/OH^-^ on the P^G^ atom (Fig. S1). However, it is yet unclear whether the reaction proceeds in two steps separated by formation of a metastable intermediate (Fig. S1A-B), or in one concerted transition with the pentavalent trigonal <u>b</u>i<u>p</u>yramidal (tbp) transition state, see Fig. S1C and [15, 48, 56–63, 133]. In the case of two-step mechanism, it is also unclear whether the reaction follows the dissociative S_N_1 pathway (Fig. 1SA) or the associative S_N_2 pathway (Fig. S1B). In the S_N_1 mechanism, the rate-limiting step is dissociation of the terminal O^3B^−P^G^ bond to form a metaphosphate intermediate, which interacts with OH^-^ yielding inorganic phosphate (P_i_), see Fig. S1A. In the S_N_2 mechanism, the rate-limiting reaction is the nucleophilic attack by OH^-^ on P^G^ and the formation of the pentavalent tbp intermediate, which then dissociates into NDP and P_i_, see Fig. S1B.

The research community is roughly evenly divided between proponents of these three mechanisms, see e.g. [63, 133, 177]. However, it is not clear whether the difference between the mechanisms is fundamental in this case. It was experimentally shown that the mechanism of ATP hydrolysis varies even in water; it changes from dissociative in pure water to associative in the presence of positively charged chelators for γ-phosphate [178].

Therefore, instead of focusing on these differences, we would like to emphasize the universally recognized common features of the reaction pathways shown in Fig. S1, namely:

1. in any model, the covalent bond between O^B3^ and P^G^ must be destabilized [57–59];
2. the γ-phosphate group undergoes a steric inversion when forming a covalent bond with the nucleophile, see Fig. S1 and [57]. Therefore, catalytic interactions should planarize the γ- phosphate [179, 180];
3. Negative charges of the oxygen atoms of γ-phosphate must be compensated to increase the electrophilicity of the P^G^ atom and make it prone to nucleophilic attack [15, 48, 56–59, 61–63, 133, 180];
4. Conditions in the catalytic site should enable deprotonation of W_cat_ by a suitable proton acceptor [57–61];
5. Upon dissociation of the O^B3^ – P^G^ bond, a large negative charge on the O^3B^ atom must be effectively compensated as it contributes significantly to the activation barrier [62, 64].

By correlating the results of our structural analysis with the literature, we outline below how P-loop NTPases may handle these five tasks.

#### 3.2.2. Main catalytic factors in P-loop NTPases

##### 3.2.2.1. Destabilization of the O^3B^ – O^3G^ bond

Even prior to activation, the NTP molecule is bound within the closed catalytic site in a conformation more extended compared to its conformation in water, as noted earlier for particular enzymes [39, 40, 180, 181]. As Figure 2A shows, this extended conformation of the bound NTP molecule is common to all major families of P-loop NTPases. In enzyme-bound NTP molecules, the β- and γ-phosphates are in an almost perfectly eclipsed conformation due to interactions with Mg^2+^ and Lys^WA^ (see Fig. 1D), in contrast to their staggered configuration in water [40, 181]. The repulsion of the eclipsed oxygen atoms has the potential to destabilize the O^3B^ – P^G^ bond [36, 62, 165, 182].

Our finding that stimulators always interact with γ-phosphate indicates that this interaction may contribute further to destabilization of the O^3B^ – O^3G^ bond. For example, the twisted γ-phosphate may become more eclipsed relative to the α-phosphate, so that the oxygen atoms of the α- and γ-phosphate repel each other, as suggested by Rudack and colleagues [183]. Moreover, any twisting or pulling γ-phosphate can destabilize the entire Mg-NTP system, inevitably disturbing the coordination sphere of Mg^2+^, since the O^1G^ atom of γ-phosphate is one of the Mg^2+^ ligands (Fig. 1D, 2).

##### 3.2.2.2. Planarization of γ-phosphate

Already in the absence of the stimulator, the oxygen atoms of the γ-phosphate are already “pulled up” to the β-phosphate by Lys^WA^, which interacts with O^1B^ and O^2G^, and by Mg^2+^, which interacts with O^2B^ and O^1G^, see Fig. 1B and [184]. The stimulatory moiety in the AG site further planarizes γ-phosphate by drawing the O^3G^ atom toward O^2A^ and enabling the interaction between **HN**^K-3^ and O^2G^ (Fig. 1-6, Table S1). Indeed, in the presence of stimulatory moieties – for example in the pre-TS structures shown in Fig. 3 of the accompanying article [41] – the O^3B^-P^G^-O^1G^, O^3B^-P^G^-O^2G^, and O^3B^-P^G^-O^3G^ angles are mostly less than the 109° expected for an ideal tetrahedron, see also [184].

Even when interacting only with γ-phosphate, the stimulator is often located between α- and γ- phosphates, as in dynamins (Fig. 3A) or ABC transporters (Fig. 5D) and can planarize γ-phosphate by tying O^3G^ to the “head” of the NTP molecule. For instance, the Na^+^ ion in dynamins, while not reaching the α-phosphate directly, is connected to it via two noncovalent bonds (Fig. 3A). The signature motif of ABC-NTPases is H-bonded via conserved Ser and Gln residues to the O2’ atom of the ribose (Fig. 5D).

Notably, the twisted or tilted γ-phosphate becomes more eclipsed relative to α-phosphate, but *less* eclipsed relative to β-phosphate [40], which might be a prerequisite for inversion of the γ-phosphate group (otherwise, in the case of an ideal eclipse, the oxygen atoms of β-phosphate would prevent the inversion of γ-phosphate).

##### 3.2.2.3. Electrostatic compensation of the negative charge of the phosphate groups

The positive charges of amino group of Lys^WA^, the Mg^2+^ ion, and several **HN** groups of the P-loop compensate for the negative charges of phosphate oxygen atoms (see Fig. 2B) so that electrons are “pulled away” from the P^G^ atom, see for example [36, 65, 176–178, 180, 183]. Notably, the phosphate chain “sits” on the last N-terminal turn of the α_1_-helix, which generally carries a dipole positive charge of about 0.5 [36, 185]. This positive charge should also contribute to electrostatic compensation. The positions of the groups involved are strictly conserved (Fig. 2), so that such a compensation is common to all major families of P-loop NTPases and may, at least partly, explain why the very binding of the GTP molecule to the catalytic site of Ras GTPase accelerated hydrolysis by five orders of magnitude, as compared to hydrolysis in water, even in the absence of an activating partner [39, 167]. In addition to these common positively charged moieties, class-specific auxiliary residues could be involved, as, for instance, “sensors 3” in some AAA+ ATPases.

On activation, the negative charges of the γ-phosphate oxygen atoms are additionally compensated by the positive charges provided by most stimulators, see Fig. 1-6 and the accompanying article [41]. The additional electrostatic compensation significantly increases the electrophilicity of the P^G^ atom. Rudack and colleagues showed in their QM/MM calculations that the insertion of the arginine finger alone increases the partial positive charge on P^G^ to 1.46 elementary charges [183]. The interaction of O^2G^ with **HN**^K-3^, as described in the accompanying article [41] and in [40] should further increase the positive charge on P^G^ in the pre-transition state.

Furthermore, Fig. 8 shows that the electrostatic potential at the catalytic sites of diverse P-loop NTPases is distributed unevenly, which has already been noted for particular enzymes of this family, see, e.g., [186]. The strength of the local electric field can be roughly estimated using the Coulomb’s equation as ∼ 10^8^ V/m under the very modest assumptions that (i) the resulting electric charge difference at a distance of 10 Å (between the acid residues and the P-loop) corresponds to one elementary charge and (ii) the effective dielectric permittivity of the catalytic pocket is about 10 [187, 188]. Such an electric field strength, albeit large, is compatible with those measured in the catalytic pockets of other enzymes [189]. Notably, the local electric field is directed approximately from the P-loop to Asp^WB^. Hence, the catalytic pocket is strongly polarized in P-loop NTPases, which also can contribute to catalysis, as discussed below.

Consequently, in those cases where the stimulator is positively charged, its positive charge not only secures the bonding with particular oxygen atom(s) of triphosphate and increases the positive charge on P^G^, but additionally polarizes the whole catalytic pocket.

##### 3.2.2.4. Does the [Ser/Thr]^K+1^ – Asp^WB^ pair accept a proton from W_cat_?

In their comprehensive review on enzymatic mechanisms of phosphate transfer, Cleland and Hengge wrote: “The problem for an ATPase … is thus to position a water molecule so that it *is* in a position to attack the γ-phosphorus. This requires steric restraints as well as organized hydrogen bonding networks. And more specifically, there must be a path for one proton of the attacking water molecule to reach a suitable acceptor “ (quoted from [60]). In P-loop NTPases, the proton from W_cat_ is commonly thought to be taken up either by γ-phosphate in TRAFAC class NTPases [67, 133] or by W_cat_-coordinating, „catalytic“ Glu or Asp residues in other classes of P-loop NTPases [8, 54, 99, 102, 114, 116, 125, 129, 190–192]. Still, our comparative structure analysis showed that such Glu/Asp residues, present in all classes of NTPases except the TRAFAC class, are non-homologous. In SIMIBI NTPases, W_cat_-coordinating Glu or Asp residues are at the C-tip of their WB+1 β-strands (Fig. 4A-B). In most classes of ASCE ATPases, the “catalytic” Glu^D+1^ follows Asp^WB^, see Fig. 5, and [14, 16, 32]. In F_1_/RecA ATPases, the “catalytic” Glu residue is located at the C-tip of the WB+1 β-strand, see Fig. 6 and [8, 14, 19]. Also, those kinases that require deprotonation of the second substrate have usually an Asp^WB^-Leu^D+1^-[Asp/Glu]^D+2^ motif where the D+2 residue interacts with the prospective nucleophilic group, see Fig. 4D and sequence alignments in [9]. Such an absence of homology is unusual for catalytically critical residues.

More importantly, neither γ-phosphate, nor W_cat_-coordinating Glu/Asp residues can hold the proton from W_cat_. Indeed, the inverse solvent isotope effect on the GAP-activated hydrolysis by the Ras GTPase upon H/D substitution [193] indicates that deprotonation is likely to occur before the rate-limiting step of bond cleavage [60]. The bond cleavage time by P-loop NTPases is in the millisecond range (100 ms at 260°K in a Ras GTPase activated by the RasGAP [194]). During this time, the proton must be “detained” in such a way as to prevent its return to OH^−^_cat_ and the reversal of the reaction. In general, the ability to keep the proton is seen as a prerequisite for the completion of enzymic reactions with participation of deprotonated nucleophiles, as discussed e.g. for the eukaryotic cAMP-dependent protein kinase [195, 196].

The common belief of proton trapping from water (pK_a_=14.0) either by a “catalytic” Glu or Asp residue in a polar environment (with expected pK_a_ in the range of 2.0-5.0) or by γ-phosphate (with pK < 3.0 [197, 198]) is at odds with the basic rules of proton transfer as formulated by Eigen [199, 200]. After Eigen, if the donor and acceptor of the proton are connected by a “hydrogen bridge”, proton transfer between them is “practically unhindered *provided the difference pK_acceptor_ – pK_donor_ is positive*” (quoted from [200]). In the case of γ-phosphate and W_cat_-coordinating Glu/Asp residues, the respective difference is strongly negative so the proton will promptly (in picoseconds) return to W_cat_.

It was speculated that the W_cat_-coordinating groups may facilitate its catalytic deprotonation by decreasing the proton affinity of W_cat_, see e.g. [201]. Our structural analysis of TS-like structures showed that positively charged side chains of Lys or Arg, which could significantly lower the pK value of W_cat_, interact with it only in a few classes of P-loop NTPases, e.g. in some AAA+ ATPases, see Section 2.2.2.2 and Table 1. Even in these cases, their positive charges can hardly decrease the apparent pK of W_cat_ by >10 pH units, which is needed for sustainable protonation of a “catalytic” Glu/Asp residue or γ-phosphate. Hence, neither “catalytic” acidic residues, nor γ-phosphate can trap a proton from W_cat_ per se.

As early as in 2004, some of these problems were recognized by Frick and colleagues [202, 203] who calculated the pK values for ionizable residues of the SF2 class Hepatitis C virus (HCV) NS3 helicase by using the MCCE software [204]. Frick and colleagues wrote about supposedly W_cat_-coordinating Glu291^D+1^ and its preceding Asp290^WB^: “to function as a catalytic base, the pKa of Glu291 would need to be much higher than that of a typical Glu in a protein. However, electrostatic analysis of all HCV helicase structures reveals that neither Glu291, nor any nearby Glu, has an abnormally high pKa. In contrast, Asp290 has a pKa as high as 10 in some structures and as low as 3 in others. Interestingly, in structures in the open conformation (such as 8OHM), the pKa of Asp290 is low, and in the closed conformation (ex. 1A1V), the pKa of Asp290 is higher than 7, suggesting that Asp290 picks up a proton (like a catalytic base) when the protein changes from the open to the closed conformation. Thus, Asp290 could serve as a catalytic base…” (quoted from [203]). Frick and colleagues made their calculations with crystal structures of helicases that contained neither Mg-ATP nor its analogs. Therefore, they were unaware of the exact positions of Asp290^D^ and Glu291^D+1^ relative to the bound substrate and could only guess by analogy with known structures of related P-loop NTPases. The two subsequently resolved ADP:AlF_4_^−^-containing, TS-like crystal structures of HCV NS3 helicase (PDB IDs 3KQL [205] and 5E4F [115]) show a typical for SF2 helicases arrangement of Asp290^WB^ and Glu291^D+1^ (see Fig. 5C). Furthermore, both structures show short 2.4 Å H-bonds between Ser211^K+1^ and Asp290^WB^, as seen in Fig. 5C.

Unlike “catalytic”, W_cat_-coordinating Glu/Asp residues, Asp^WB^ is strictly conserved in P-loop NTPases. Equally strictly conserved is the [Ser/Thr]^K+1^ residue [9, 10, 12, 206]. Both these residues can translocate protons. Therefore, building on our comparative structure analysis and following Frick and colleagues who suggested Asp290^WB^ as a catalytic base in the HCV NH3 helicase [202, 203], we propose here that the buried, H-bonded pair of strictly conserved [Ser/Thr]^K+1^ and Asp^WB^ serves as a universal module that accepts the proton from W_cat_ and holds it as long as needed in P-loop NTPases of all classes.

It can be countered that the [Ser/Thr]^K+1^ – Asp^WB^ pair does not interact directly with W_cat_. However, direct tracing of intra-protein displacements of protons in energy converting enzymes and chemical models (see [199, 200, 207–219] for reviews) showed that fast proton transfer over a distance of up to 20 Å can be mediated by water bridges, provided that the distance between the groups involved is ≤ 3.0 Å, in accordance with Eigen [200]. Thereby it does not matter that water molecules are equally poor proton acceptors (with pK_b_ of 0.0) and proton donors (with pK_a_ of 14.0) because protons pass water by so-called von Grotthuss mechanism [220–222]. According to the modern version of this relay mechanism, the bridging molecule receives an external proton simultaneously with the transfer of its own proton to the next carrier at picoseconds [221, 222]. Apart from water molecules, the side chains of serine, threonine, tyrosine, and neutral histidine are suitable for proton transfer by the von Grotthuss mechanism: they have a proton-accepting lone pair of electrons and also their own proton to transfer on [223]. In addition, protons are transferred very fast between carboxy groups when they are bridged by single water molecules [199, 200, 213, 217]. Zundel and colleagues showed that such systems are highly polarized [207, 224], which lowers the activation barriers for proton transfer.

The best-studied examples of such proton-conducting chains are known from the photochemical reaction center (PRC) of α-proteobacterium *Rhodobacter sphaeroides* (as shown in Fig. 10A and described in its caption) and bacteriorhodopsin of an archaeon *Halobacterium salinarum* (as shown in Fig. 10B and described in its caption).

**Figure 10.**
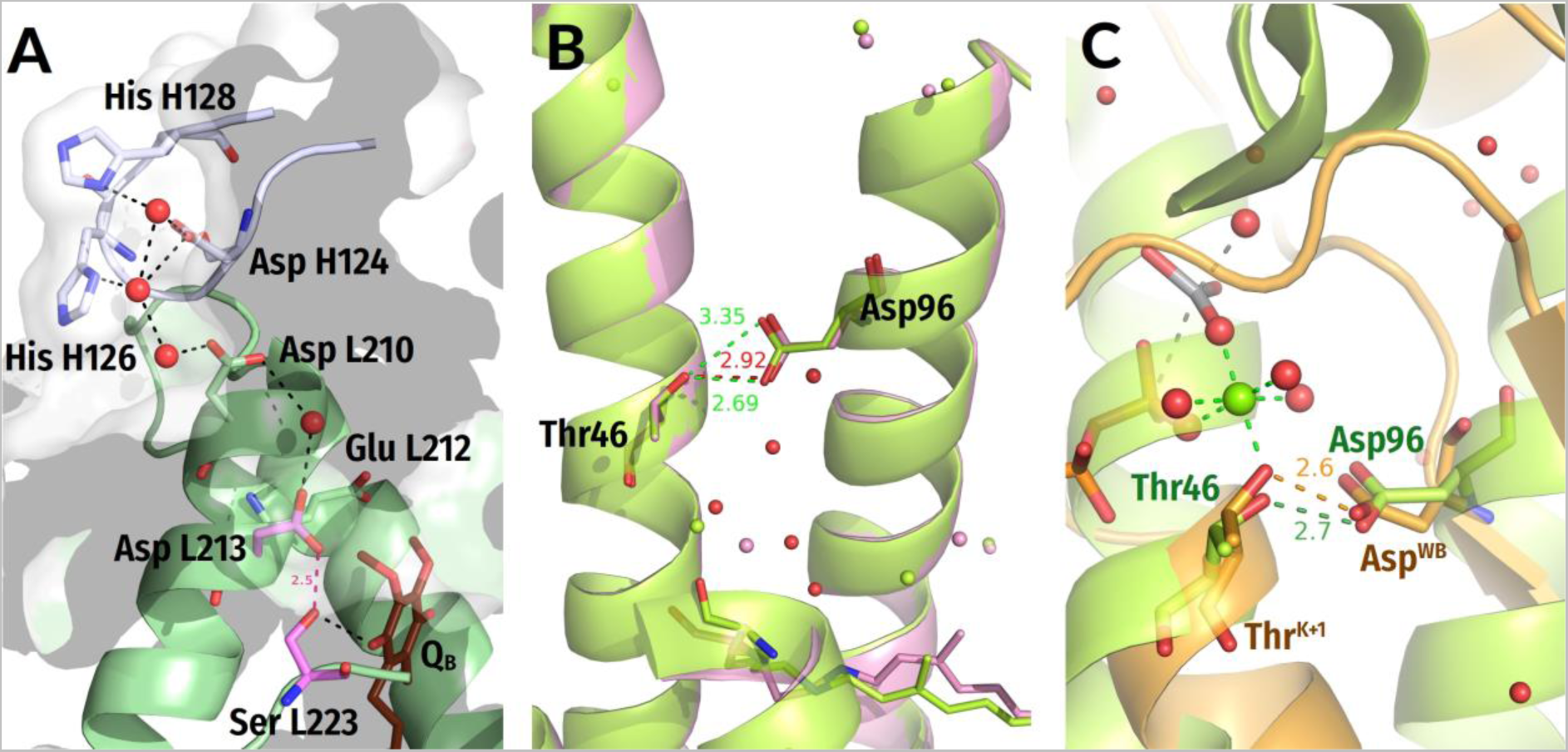
Proton traps in the photochemical reaction center and bacteriorhodopsin as compared with the Thr^K+1^-Asp^WB^ pair of P-loop NTPase. Distances are given in ångströms. **A**, a high-pK proton trap in the photosynthetic reaction center (PRC) from *Rhodobacter sphaeroides.* Shown is the PRC in the charge-separated state (PDB ID 1DV3 [225]). Here, a proton path connects the protein surface with the buried binding site of the secondary ubiquinone acceptor Q_B_. The proton is translocated over more than 20 Å via amino acid residues of the two PRC subunits, L and H. The proton pathway begins at the two His residues on the surface (His-H126 and His-H128); they are thought to have pK values of about 7.0 and to harvest protons which are ejected onto the photosynthetic membrane surface by the ATP synthase. From these histidine residues, the proton goes, via water-bridged carboxy groups of Asp H-124 and Asp-L210, to the Q_B_ -binding pocket. Proton transfer from the surface to the more hydrophobic interior of the protein becomes possible after the Q_B_ molecule docks to the Glu-L212 – Asp-L213 – Ser-L223 cluster [226] and turns it into a buried, high-affinity proton acceptor (a *membrane proton trap* according to Mitchell [227]), with a functional pK of about 9.5 – 10.0 [228–232]. This high functional pK is apparently due to the expulsion of water upon Q_B_ binding, the negative charge on the Glu-L212 – Asp-L213 pair, and the H-bond between Asp-L213 and Ser-L223, through which the proton passes to Q_B_. The functional pK value of the trap increases to > 12.0 after the appearance of an electron at Q_B_ and the formation of the Q_B_^-·^ anion radical, which attracts further protons into the buried catalytic site [212]. Mutations of either Glu-L212, or Asp-L213, or Ser-L223 block proton transfer from the surface [228, 233, 234]. For further details, see [216, 225, 235–237]. **B,** a high-pK proton trap in bacteriorhodopsin (BR), a membrane protein that pumps a proton across the membrane in response to the photoexcitation of its retinal pigment, see [209, 210, 238, 239] for reviews. Shown is the superposition of femtosecond X-ray laser-captured structures of bacteriorhodopsin from [240] in a closed resting state (PDB ID 6G7H, light-green) and in an “open” state 8.3 ms after illumination (PDB ID 6G7L, pink). Additional red-colored water molecules were taken from a high-resolution crystal structure of the V49A bacteriorhodopsin mutant that was crystalized in an “open” state (PDB ID 1P8U, [241]). In the BR, the key proton carrier Asp-96 has pK of ∼12.0 owing to the absence of water in the vicinity and a H-bond with Thr-46 of the nearby α-helix. Replacement of Thr46 by a valine decreased the apparent pK of Asp96 by approximately two pH units [242], which may characterize the contribution of the H-bond between Thr46 and Asp96 to the unusually high apparent pK of the latter. The photoisomerization of the retinal twists the α-helices and lets water molecules in the space between them, see the pink structure [209, 218, 219, 243-246, 247. The pK value of Asp-96 shifts to 7.1, which allows its deprotonation and proton transfer, via a transiently formed water chain, to the Schiff base of retinal at about 10 Å, see {Luecke, 2000 #3390, 248, 249]. Later, the cleft between helices closes again, the pK of Asp-96 returns to 12.0, and it is re-protonated from the surface [209, 210, 239, 244]. **C,** the Thr46-Asp96 pair of bacteriorhodopsin it its resting closed state (the green structure from panel 10B, PDB ID 6G7H) superimposed with the Thr^K+1^—Asp^WB^ pair from the transition-state like, VO_4_^-^-containing structure of myosin shown in Fig. 3D, orange (PDB ID 1VOM [95]).

The two proton transfer routes shown in Fig. 10A-B were reconstructed from direct electrometric tracking of flash-induced proton displacements [209, 250–254], kinetic IR and UV/Vis spectroscopy data [209, 210, 230, 231, 248, 251, 255, 256], kinetic ESR measurements [238, 243, 245, 257], and comparative structure analyses [218, 226, 236, 244]. These are the two best-understood cases of intra-protein proton transfer by far.

In the case of PRC (Figs. 10A), the buried proton acceptor must have a functional pK value greater than 9.0 to compensate for the desolvation penalty of about three pH units, which must be “paid” for delivery of a proton from a pK-neutral “antenna” group at the protein surface into the hydrophobic membrane [258, 259]. However, if such a buried group is equilibrated with the protein surface, its pK value can hardly exceed the ambient pH, which is usually neutral, because the groups with even higher pK values are already protonated. Hence, the strong proton acceptor shown in Fig. 10A is not fully equilibrated with the surrounding solution. In this case, it is better to speak about the *apparent/functional* pKa or high *proton affinity*. Such very strong proton acceptors usually emerge only transiently [232, 260].

The proton traps in Fig. 10A, B include H-bonded [Ser/Thr] – Asp pairs (modules). These examples show that proper H-bonding of a protein-buried Asp residue with a nearby Ser/Thr residue in a hydrophobic environment can generate a trap with a high proton affinity; such traps can, in principle, accept protons even from water. Specifically, in the absence of pH-buffers and protons from the ATP synthase, the protons that pass through the PRC to compensate the negative charge at the Q_B_ site (Fig. 10A) appear to detach from surface water molecules, presumably polarized by surface charges [256].

Our suggestion that a similarly H-bonded [Ser/Thr]^K+1^ – Asp^WB^ pair accepts the proton from W_cat_ in P-loop NTPases of different classes challenges the current ideas on catalytic proton transfer in P-loop NTPases. Therefore, we further substantiate the suggested mechanism below and illustrate it in Fig. 11 and 12:

1. In TS-like structures of P-loop NTPases of various classes (except the TRAFAC class), the “catalytic” Glu/Asp residues were found to connect W_cat_ with ligands of Mg^2+^ in positions #3 or #6, see Fig. 1F, 4-6, 7A. Notably, the six ligands of Mg^2+^ form a regular octahedron with edges 2.9-3.0 Å long, so that the ligands # 3 and #6 are on a H^+^-transfer distance from [Ser/Thr]^K+1^ (Fig. 11). The short H-bond between [Ser/Thr]^K+1^ and Asp^WB^ completes the proton-conducting pathway that connects W_cat_ with Asp^WB^. The proton pathways from W_cat_ to Asp^WB^, which resemble proton translocation systems of PRC and BR (cf Fig. 10), are shown by the red dashed lines for P-loop NTPases of different classes in Fig. 4-6 and by dashed arrows in Fig. 11.
2. The pKa of an aspartate residue in water is about 4.0, much lower than that of water (14.0). However, unlike the “catalytic” Glu/Asp residues or γ-phosphate surrounded by charged residues (Fig. 1E, 1F, 3-6), Asp^WB^ is in a nonpolar environment and its functional pK is likely to be high when the catalytic site is closed. Asp^WB^ is in the middle of an αβα sandwich, on the interface between the β-pleated sheet and the α_1_-helix; such interfaces are stabilized by hydrophobic interactions [185]. In addition, Asp^WB^ is preceded by four hydrophobic residues of the Walker B motif (Fig. 1); the adjacent β strands, as well as the α_1_-helix, also contain many hydrophobic residues, see the sequence alignments in [9, 10, 12, 206]. Upon constriction of the catalytic pocket and expulsion of eventually present water molecules, the hydrophobic environment should elevate the proton affinity of the H-bonded Asp^WB^, as it happens with similarly H-bonded Asp96 which has a functional pKa of ∼ 12.0 in a hydrophobic environment of the ground-state BR. Fig. 10C shows that the structure of the Ser186^K+1^ – Asp454^WB^ pair of myosin overlaps nicely with Thr46 – Asp96 pair of BR.
3. In contrast, the pK of [Ser/Thr]^K+1^, which is about 13.0 in water, is likely to be reduced when [Ser/Thr]^K+1^ serves as a Mg^2+^ ligand. Coordination of a Zn^2+^ ion by a serine side chain is known to decrease the pK value of the latter up to 5.5 yielding a serine anion (alkoxide) at neutral pH, see [60] and references therein. The impact of a Mg^2+^ ion should be weaker; still, within a closed/constricted catalytic site, the low dielectric permittivity would enhance electrostatic interactions. [Ser/Thr]^K+1^ is the most deeply buried of the Mg^2+^ ligands (see Fig. 1-6, 11), so it should be the most sensitive to electrostatic effects. As a result, the functional pK value of [Ser/Thr]^K+1^ may dramatically decrease upon constriction of the catalytic site.
4. The found shortening of H-bonds in the TS-like structures (Fig. 9) enables estimation of the difference in functional pK values of [Ser/Thr]^K+1^ and Asp^WB^ in the TS. It is well established, based on ample experimental evidence, that hydrogen bonds “generally shorten as ΔpKa, the difference in the donor and acceptor pKa values, decreases” (quoted from [261]). Specifically, Herschlag and colleagues observed, on various systems, that the Δp*K*a decreases linearly from 20 to 0 with the decrease in the O—H••••O distance from 2.9 Å to 2.4 Å, with a slope of 0.02 Å/p*K*a unit, [261, 262]. In NDP:AlF_4_^-^-containing structures with constricted catalytic site, the length of the H-bonds between [Ser/Thr]^K+1^ and Asp^WB^ varies around 2.5 Å (Fig. 1E-F, 3, 4A-B, 5C-D, 6A, 9), which corresponds to ΔpK < 3.0 and indicates a low-barrier hydrogen bond [261, 262]. Hence, [Ser/Thr]^K+1^ and Asp^WB^ may have comparably high proton affinities in a constricted catalytic site.
5. Limbach and colleagues combined low-temperature UV-Vis and ^1^H/^13^C NMR spectroscopy (UVNMR) to study the effect of solvent polarity on the proton equilibrium between phenols and carboxylic acids [263–265], which system can be viewed as a model of the [Ser/Thr]^K+1^—Asp^WB^ H-bonded pair. These authors have shown that proton relocates from the hydroxy group to the carboxyl with decrease in polarity.
6. In the octahedral coordination shell of Mg^2+^, the O^1G^ atom of γ-phosphate is the ligand opposite to [Ser/Thr]^K+1^ (Figure 11). Therefore, the stimulator-induced rotation of γ-phosphate, by moving O^1G^ in *<u>any</u>*direction (as shown by dashed purple arrows in Fig. 11), would inevitably increase the distance between O^1G^ and the hydroxyl of [Ser/Thr]^K+1^. Pulling away the negatively charged O^1G^ will increase the cumulative positive charge at [Ser/Thr]^K+1^ prompting the relocation of its proton to Asp^WB^, e.g. in response to a thermal fluctuation [265] (Fig. 12A).
7. We suggest that the resulting Mg^2+^-coordinated Ser/Thr anion (alkoxide), used as a proton acceptor from water by many enzymes [60, 164, 266, 267], withdraws the proton from W_cat_ (or the sugar moiety in some kinases) via proton pathways shown in Figs. 4-6, 7A, 11, 12B; this proton transfer is additionally driven by strong local electric field (see Fig. 8 and Section 2.2.4). The resulting state where both [Ser/Thr]^K+1^ and Asp^WB^ are protonated corresponds to the ground state of the Thr46-Asp96 pair in the BR (see the light-green structures in Fig. 10B-C).
8. The formed anionic nucleophile (e.g. **OH^—^** ) stabilized and polarized by its ligands, is attracted by the electrophilic P^G^ atom (Fig. 12B). The proton affinity of the anionic nucleophile decreases as it gets closer to P^G^, so that proton return from the [Ser/Thr]^K+1^— Asp^WB^ couple becomes increasingly unfavorable, eventually satisfying the Eigen’s condition for proton transfer and making it complete.
9. In diverse P-loop NTPases, mutations of Asp^WB^ or [Ser/Thr]^K+1^ diminished the enzyme activity dramatically, see e.g. [203, 268–275]. More specifically, mutations of Asp^WB^ to Asn retarded the activated hydrolysis without affecting the NTP binding [203, 268–272]. In the case of *E.coli* F_1_-ATPase, the mutation even increased the affinity for ATP [270]. The Asp^WB^ to Asn mutation mimics the charge state of a protonated Asp^WB^. Hence, the protonation of Asp^WB^ is unlikely to distort the catalytic pocket and be the cause of the universal catalytic incompetence of the Asp^WB^ to Asn mutants. We attribute this incompetence to the inability of Asn^WB^ to trap a proton from [Ser/Thr]^K+1^.

**Figure 11.**
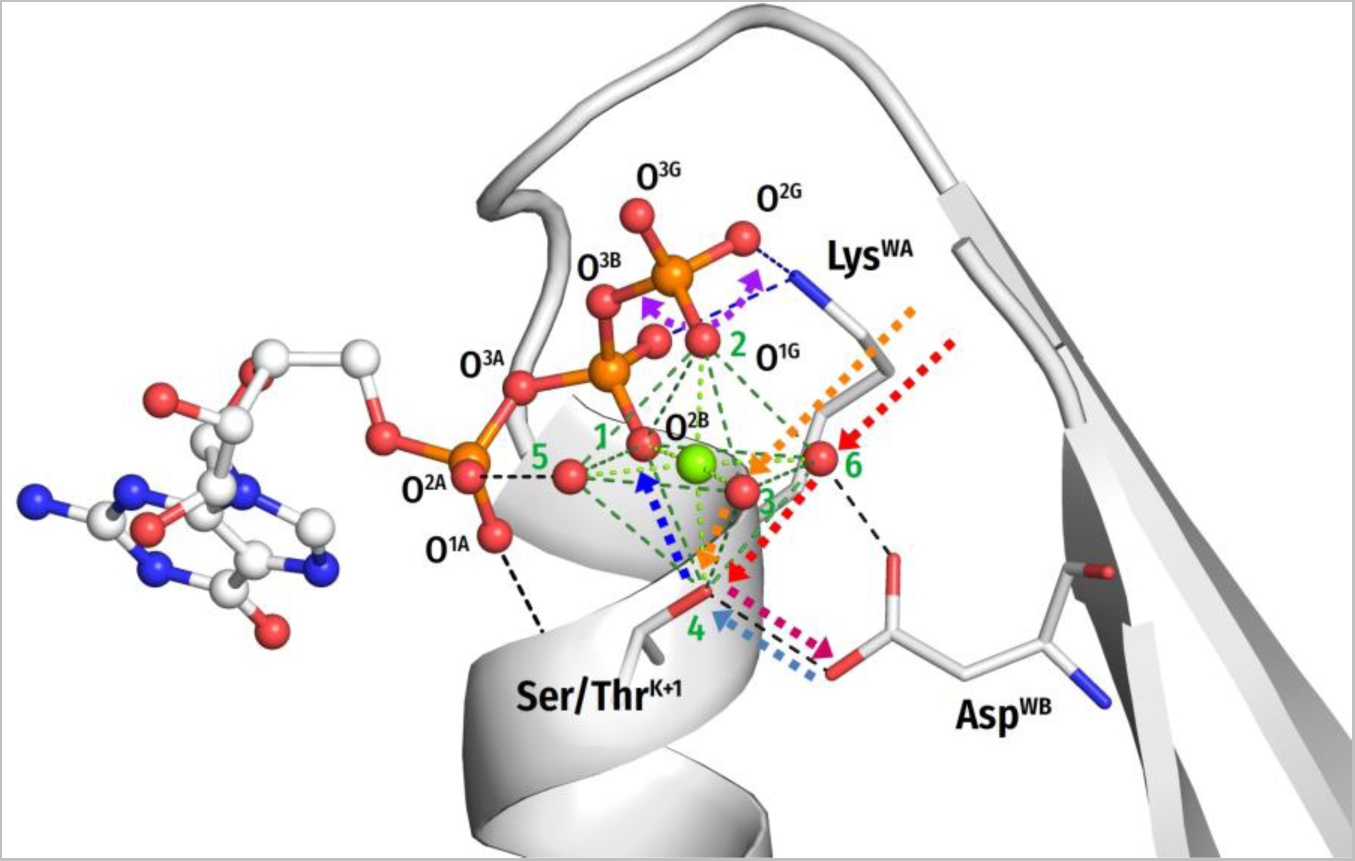
Schematic presentation of tentative proton routes along the edges of the octahedral coordination shell of Mg^2+^ ion. The Mg^2+^ ligand #3 is assumed to be a water molecule. Proton entry points via ligand #6 or ligand #3 are shown by red and orange arrows, respectively. Protonic connection between [S/T]^K+1^ and Asp^WB^ are shown by magenta and light blue arrows, the route from [S/T]^K+1^ to O^2B^ of β-phosphate is shown as a dark blue arrow. The movement of the O^1G^ atom as a result of γ-phosphate twist is shown by purple arrows.

**Figure 12.**
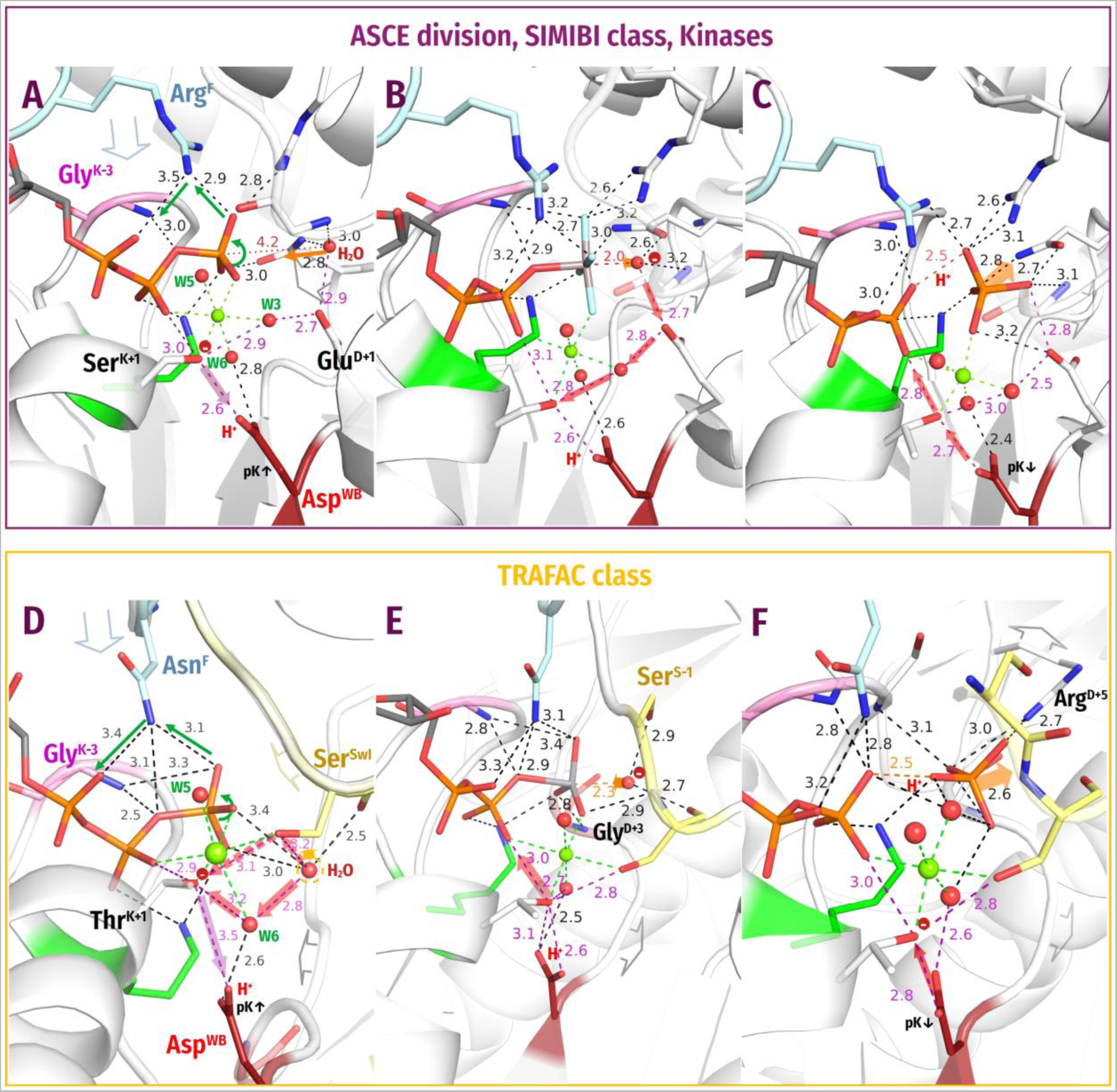
Two tentative mechanisms of proton transfer from W_cat_ to Asp^WB^ in P-loop NTPases. The shade of the red arrows of proton transfers varies through panels A-C and D-F to emphasize that the kinetic coupling of these steps with each other and reactions in the catalytic site could vary depending on the thermodynamics of particular enzymes. Other colors as in Fig. 3-6. <u>Top (**A-C**): Tentative mechanism of proton transfer via the **W**</u>**<u>_cat_</u>**<u>-coordinating Glu/Asp residue in all classes of P-loop NTPases but TRAFAC class enzymes</u>, as illustrated by structures of the SF1 helicase Pif1. **A**, In the pre-catalytic state, which is exemplified by a structure with a bound non-hydrolyzable ATP analog ANP (PDB ID 6HPH [276]), the binding of ATP and closure of the catalytic site increases the proton affinity of Asp^WB^. Activation-induced constriction of the catalytic site and twist of γ-phosphate prompt the proton redistribution from [Ser/Thr]^K+1^ to Asp^WB^; **B**, in the TS, which is exemplified by a structure with a bound TS analog ADP:AlF_4_^−^ (PDB ID 5O6B [192]), a proton from W_cat_ repletes the vacancy at the anionic [Ser/Thr]^K+1^ by going via the W_cat_-coordinating Glu^D+1^ and W3; the resulting **OH^-^_cat_** attacks γ-phosphate; **C**, in the post-TS, which is exemplified by a structure with ADP:MgF_4_^2-^ bound (PDB ID 6S3I [277]), where we replaced the H_2_PO ^2-^ mimic MgF_4_^2-^ by H_2_PO_4_ ^2-^, the proton goes from Asp^WB^, via Ser^K+1^, to β-phosphate to compensate its negative charge (see the main text). Bottom (**D-F**): Tentative mechanism of proton transfer from W_cat_ to Asp^WB^ in TRAFAC class <u>NTPases</u>, as illustrated by structures of myosin. **D**, For the pre-catalytic state we used the ATP-soaked crystal structure of myosin II (PDB ID 1FMW [139]) where the Switch I loop was taken from the structure of Myosin V complexed with ADP-BeF_3_ (PDB ID 1W7J [278]) and superimposed via Lys^WA^ and Mg^2+^. In this state, the binding of ATP and closing of the catalytic site increases the proton affinity of Asp^WB^. Constriction of the catalytic site upon activation and twist of γ-phosphate prompt the proton redistribution from Thr^K+1^ to Asp^WB^ (purple arrow). The anionic Thr^K+1^ accepts a proton from W6 that, in turn, takes a proton from W_12_ (red arrows, cf. Fig. 7B-C); alternatively, the proton route from W_12_ to Thr^K+1^ may involve the conserved Ser^Sw1^ (dashed red arrows) The resulting **OH^—^_cat_** is brought into the apical position by residues of Switch I (colored yellow) and WB-crest; **E**, in the transition state, which is exemplified by a structure with a bound TS analog ADP:VO_4_^−^ (PDB ID 1VOM [95]), **OH^—^_cat_** attacks P^G^. In this constricted transition state, the protonic connection between **OH^—^_cat_** and the Mg^2+^-coordinating water molecules appears to be broken. In the subsequent stage of the catalytic transition, the proton goes from [S/T]^WA^ to β-phosphate and compensates for the large negative charge that builds up upon the detachment of γ-phosphate (the red arrow); **F**, in the post-transition state, which is exemplified by a structure with bound ADP and H_2_PO_4_ ^2-^ (PDB ID 4PJK [52]), the negative charge on β-phosphate is compensated by a hydrogen bond between β- and γ-phosphate. Asp^WB^ repletes Thr^K+1^ concomitantly with the opening of the catalytic pocket (see also the main text).

In TRAFAC class NTPases, the H-bond between [Ser/Thr]^K+1^ and Asp^WB^ is as short in the TS-like structures as in other classes of P-loop NTPases, see (Fig. 1F, 3) and Table S1. However, these NTPases have no “catalytic” W_cat_-coordinating Glu/Asp residues. Instead, they all have a strictly conserved [Thr/Ser]^SwI^ residue that coordinates Mg^2+^ by its side chain as ligand #3 and, in the same time, stabilizes W_cat_ with its **CO** group in the TS-like configuration (Fig. 1F, 3). In this configuration, the side chain oxygen of [Ser/Thr] ^SwI^ is exactly 2.9 Å away from the side chain oxygen of [Ser/Thr]^K+1^, as they are neighboring ligands in the octahedral coordination shell of Mg^2+^ (see Fig. 11). In the absence of a bound TS-analog, however, [Thr/Ser]^SwI^, as well as the whole Switch I loop, are highly mobile with water sporadically being involved as the Mg^2+^ ligand # 3 instead of [Thr/Ser]^SwI^, see Fig. 7B-C, S5B and [140, 279]. It is tempting to speculate that the side chain of mobile [Thr/Ser]^SwI^ may be involved in a von Grotthuss-type proton transfer from the would-be W_cat_ to [Ser/Thr]^K+1^ in the pre-TS state. Kumawat and colleagues, studying the small Rho GTPase by MD simulations, have shown that seven water molecules, on average, come within 4 Å distance from Thr37^SwI^ in a closed, GTP bound state [280]. This amount should be sufficient to transfer a proton from the would-be W_cat_ to anionic [Ser/Thr]^SwI^ over a distance of just 4– 5 Å concomitantly with the constriction of the catalytic site. Alternatively, a W_12_ or W_13_ water molecule can give its proton to [Ser/Thr]^SwI^ and then move by 1-2 Å into the apical position turning into **OH^—^_cat_**. For myosin, where W_12_ is “visible” owing to extra bonds with residues of Switch I (Fig. 7D), we could reconstruct two tentative/alternative proton route(s), as shown by solid and dashed red arrows in Fig. 12D. Similar proton pathways can be proposed for Ras-like GTPases based on the “open” structure shown in Fig. 7C.

In TS-like structures, as already mentioned, [Ser/Thr]^SwI^ is fixed in a distinct position where it coordinates both the W_cat_ and Mg^2+^; here the distance from its hydroxyl group to W_cat_ is about 4.5 Å (Fig. 3A-D), there appear to be no water molecules in between. It is tempting to speculate, that this configuration may hamper the unwanted return of a proton from Asp^WB^ to OH^—^_cat_.

##### 3.2.2.5. Charge compensation at β-phosphate

According to Herschlag and colleagues [62, 64], the activation barrier of NTP hydrolysis is due to the strong negative charge that develops at β-phosphate as γ-phosphate breaks away; this charge should be compensated upon catalysis. The post-TS structures of the RhoA/RhoA-GAP complex (PDB ID 6R3V [201]) and activated myosin (PDB IDs 4PFP, 4PJK [52]) with NDP and detached P_i_ (Fig. 12F), as well as the structure of SF1 helicase Pif1 with a H_2_PO_4_^2-^ mimic ADP:MgF_4_^2-^ bound (Fig. 12C) show that the negative charge of β-phosphate is compensated by the joint action of Mg^2+^, Lys^WA^, Arg/Lys/Asn fingers and **HN**^K-3^. At this stage, the H-bond between **HN**^K-3^ and O^2G^ is lost so that **HN**^K-3^ is fully involved in compensating for the negative charge at β-phosphate. In addition, the structures show a short H-bond of 2.4-2.5 Å between the oxygen atoms of β-phosphate and P_i_ (see Fig. F12C, F and [52, 201, 277, 281]); this H-bond apparently also compensates for the negative charge at β-phosphate.

The proton for the H-bond between β-phosphate and P_i_ is thought to stem from W_cat_ [52, 201, 281] and should be somehow transferred to β-phosphate. In the octahedral coordination shell of Mg^2+^, [Ser/Thr]^K+1^ is 2.9 Å away from β-phosphate, so that its proton can directly pass on and compensate the negative charge (as shown in Fig. 11, 12C, 12E). The proton vacancy on [Ser/Thr]^K+1^ could be then refilled by a proton from Asp^WB^ (Fig. 11, 12C, 12F), thus restoring the initial protonic configuration. Since the octahedral arrangement of Mg^2+^ ligands (Fig. 11) is similar in all P-loop NTPases, the proton can relocate from Asp^WB^ to β-phosphate, via [Ser/Thr]^K+1^, in all of them.

#### 3.2.3. Evidence for transient protonation of Asp^WB^ from infrared spectroscopy data

In PRC and BR shown in Fig. 10, the protonation of Asp/Glu residues was directly followed in the infrared (IR) spectral range [210, 230, 231, 239, 248, 255, 282]. In general, Asp and Glu residues are unique because the ν(C=O) vibration of their protonated carboxyl groups absorbs in the 1710-1760 cm^-1^ spectral region that is free from overlap with absorption of other protein components [283]. Specifically, the protonated Asp-96 of BR has an absorption maximum at 1741 cm^-1^ [248], whereas the protonated GluL212-AspL213-Ser223 complex of PRC absorbs at 1728-1725 cm^-1^ [230, 231].

Application of the IR-spectroscopy to P-loop NTPases is complicated by the transient nature of Asp^WB^ protonation – proton passes through Asp^WB^ on the time scale of the NTPase turnover. Nevertheless, IR measurements in this spectral range were performed by Kim and colleagues who studied human Eg5, a kinesin-like motor protein of TRAFAC class, where Asp265^WB^ and Thr112^K+1^ are connected by a short H-bond of 2.59 Å (PDB ID 3HQD [284]). In their steady state experiments [285], Kim and colleagues investigated the action of monastrol, an allosteric inhibitor that binds some 12 Å away from the catalytic site [286] and increases the reversals in this enzyme by hampering the release of H_2_PO_4_^2-^ [287]. In the presence of monastrol, an absorption maximum at 1726-1722 cm^-1^ was recorded and attributed to the protonation of a carboxylic group [285]. These data are consistent with proton trapping at Asp265^WB^ when the overall equilibrium of ATP hydrolysis shifts to the left, as it happens in the presence of monastrol [287].

In kinetic experiments, the FTIR spectra of human Eg5 were monitored in real time and in response to the photorelease of caged ATP [288]. In the absence of microtubules, the hydrolysis by Eg5 proceeded at a time scale of seconds, which facilitated the IR measurements. A sharp absorption maximum at 1743 cm^-1^ appeared at approx. 3 s and then decayed, whereas H_2_PO_4_^2-^, as measured at 1049 cm^-1^, appeared at approximately 5 s and reached its maximum at 10 s (see Fig. 1 in [288]). According to the views of the time, the authors attributed the maximum at 1743 nm to an “organized water cluster undergoing protonation” (quoted from [288]). This attribution, however, seems far-fetched. In proteins, protonated water clusters absorb at higher wave numbers of 1900 - 1800 cm^-1^ and show very broad continuum bands [282]. It is also unlikely that a protonated water cluster with pK around zero could have a lifetime of seconds at neutral pH of the experiment. It is tempting to suggest that the well-defined, sharp transient maximum at 1743 cm^-1^ was in fact due to the transient protonation of Asp265^WB^ next to Thr112^K+1^ in human Eg5 (cf with the maximum at 1741 cm^-1^ of Thr46-bound, protonated Asp96 in BR [248]).

#### 3.2.4. Re-assignment of functions in and around the Walker motifs

In the framework proposed here, the Walker B motif serves as a trap for the proton from W_cat_, whereas the coordination shell of the Mg^2+^ ion functions as an octahedral proton transfer hub with almost all Mg^2+^ ligands involved (Fig. 11). The proton is transferred from W_cat_ to Asp^WB^ in two steps: first, the proton shifts from [Ser/Thr]^K+1^ to Asp^WB^ in response to the constriction of the catalytic site and stimulatory interaction, and then a proton from W_cat_ fills the proton vacancy at [Ser/Thr]^K+1^, as shown in Fig. 12. Totally, one proton relocates from W_cat_ to Asp^WB^. Still, the existence of separate proton transfer steps, as shown in Fig. 11, 12 by different shades, provides flexibility; the mechanism could be adapted to the dissociative, associative, or concerted mechanism of hydrolysis. For instance, if the reaction is concerted, the proton from W_cat_ can pass directly through the Mg^2+^-coordinating ligands to β-phosphate, with the proton vacancy at [Ser/Thr]^K+1^ being filled by Asp^WB^ later, concomitantly with the loosening of the catalytic site and decrease in proton affinity of Asp^WB^, as shown in Fig. 12D-F.

The Asp^WB^ residue is almost strictly conserved throughout P-loop NTPases, see Table 1 and [9, 10, 12, 32, 107]. Only in several cases its function is performed by Glu, see [32] for details. Some of such cases are shown in Fig. S7 and described in its extended caption. The tentative interplay between Glu^WB^, Asp^E+1^ and Glu^E+4^ residues in Vir/PilT-like ATPases is considered separately in the Supplementary File 1 together with other ambiguous cases.

Notably, the cause for the strict conservation of the entire Walker B motif has remained obscure so far; to our best knowledge, no function has been attributed to the motif as a whole. The here suggested scheme invokes Asp^WB^ as the common terminal acceptor of a proton from W_cat_. Other four hydrophobic residues of the same motif serve as “hydrophobic protonic insulators”. They may be needed to increase the proton affinity of Asp^WB^ in the constricted catalytic site and to prevent eventual unwanted proton escape from Asp^WB^.

In our model, [Ser/Thr]^K+1^ also acquires new key functions as a catalytic nucleophilic alkoxide and a von Grottuss-type proton carrier, which may explain its strict conservation, see Table 1. A few known exceptions are shown in Fig. S8 and described in its extended caption.

Most common outliers are glycine residues which substitute for [Ser/Thr]^K+1^ in several distinct families of P-loop nucleotide monophosphate kinases [9]; in these cases, a water molecule serves as the Mg^2+^ ligand #4, see Fig. S8B. On the one hand, the successful involvement of a water molecule as the 4^th^ ligand of Mg^2+^ in these kinases indicates that the conservation of [Ser/Thr]^K+1^ in all other classes of P-loop NTPases may be unrelated to its function as a Mg^2+^ ligand. On the other hand, these multiple losses of [Ser/Thr]^K+1^ provide additional support for here proposed mechanism. It is in these P-loop kinases that the attacking nucleophile – the anionic phosphate moiety of a nucleotide monophosphate – needs no deprotonation and, therefore, does not need a catalytic alkoxide. As noted earlier, kinases do not interact with separate activator proteins; they are activated by binding the second substrate, which causes covering of the catalytic site by the Lid domain and the insertion of stimulatory finger(s) (Fig. 4C-D, S8B). Therefore, there is always some danger that the insertion of the finger(s) will stimulate a nucleophilic attack on γ-phosphate not by the anionic phosphate group of the second substrate but by a haphazard water molecule, leading to a futile ATP hydrolysis. Within our proposed scheme, an unwanted ATP hydrolysis can be prevented by replacing [Ser/Thr]^K+1^ with a residue incapable of accepting a proton from water. And that is what independently happened in several lineages of nucleotide monophosphate kinases, see Fig. S8B and multiple alignments in [9]. In particular, all the human adenylate kinases have glycine residues in the K+1 position except adenylate kinase 6, which has a threonine residue [289]. And it is for this kinase that both kinase and ATPase activities have been shown [290]. Hence, the consistent loss of [Ser/Thr]^K+1^ – independently in several families of nucleotide monophosphate kinases [9] – finds a plausible explanation.

Neither of the exceptions in Fig. S7 and S8 calls into question the suggested mechanism where the buried [Ser/Thr]^K+1^ – [Asp/Glu]^WB^ module turns into a deep proton trap after the constriction of the catalytic site in most P-loop NTPases.

Why then the strictly conserved, H-bonded [Ser/Thr]^K+1^ – Asp^WB^ pair has not established itself as a proton acceptor from W_cat_ so far? The theory of catalysis by P-loop NTPases was developed initially and intensively for small GTPases of TRAFAC class, such as G_α_-proteins and Ras-like GTPases. Their W_cat_ molecules are stabilized not by a “catalytic” Glu/Asp, but by **CO**^SwI^, Gly^D+3^ and Gln^D+4^ (Fig. 1F, 3C), so that Gln^D+4^ was initially suggested as the catalytic base. Warshel and colleagues challenged this view arguing that a Gln residue with pK < −2.0 is unlikely to accept a proton from water [67]. Among possible alternatives, they considered Asp57^WB^ of Ras GTPase but discarded it because its mutation did not affect the slow “intrinsic” activity of the lone GTPase. Instead, the γ-phosphate group was proposed as a universal proton acceptor for W_cat_ [67] and has been considered as such ever since in TRAFAC class NTPases. Although just a year later it had been shown that mutations of Asp^WB^ slowed by orders of magnitude the fast activated hydrolysis by Ras/RasGAP complexes [272], the role of γ-phosphate as the proton acceptor was not revisited. After Cleland and Hengge noted that the direct proton transfer from W_cat_ to γ-phosphate is not possible “as the geometry does not permit such a four center reaction” [60], the involvement of an additional water molecule between W_cat_ and γ-phosphate was hypothesized for TRAFAC class NTPases. In the case of other NTPase classes, the W_cat_-coordinating Glu (Asp) residues were considered as “catalytic” bases. Unfortunately, the seminal work by Frick and colleagues on the suitability of buried Asp^WB^ as a proton acceptor from W_cat_ [202, 203], mentioned above, has not changed the common view. The inability either of γ-phosphate or “catalytic” Glu/Asp residues to serve as proton traps because of their low pK values has been overlooked.

Our suggestion that the proton affinity of Asp^WB^ depends on the length of its H-bond with [Ser/Thr]^K+1^ explains why mutations of Asp57^WB^ affect the fast RasGAP-activated GTP hydrolysis [272], but have no impact on the slow intrinsic hydrolysis by lone Ras GTPases [67]. In lone GTPases, the difference between the time of proton equilibration over the whole protein (microseconds) and the catalysis time (minutes/hours) is so large that the mere presence of a dedicated proton acceptor for W_cat_ is unlikely, see also [62]. Since the catalytic site is fully exposed in the absence of a GAP, W_cat_ eventually losses a proton to the neutral bulk solution. Analogously, some group with a pK value slightly higher than pH provides a proton for β-phosphate and then is repleted from the solution. Anyway, the terminal acceptor/donor of the proton is the neutral bulk solution so that the overall reaction is slow (as discussed elsewhere in relation to proton transfer in PRC [256]). Hence, the intrinsic hydrolysis in the absence of the GAP is insensitive to mutations of Asp^WB^ because this residue is unlikely to be specifically involved. In the exposed catalytic site of a lone Ras GTPase, the pK value of Asp^WB^ is expected to be ≤ 5.0 (which corresponds to the 2.6-2.7 Å length of the [Ser/Thr]^K+1^ – Asp^WB^ H-bond in the structures that contain NTPs or their analogs, see Fig. 9 and [261]), so that Ser17^K+1^ is protonated (see the neutron crystal structure of Ras, PDB ID 4RSG, [291]) and cannot accept a proton from W_cat_. Both [Ser/Thr]^K+1^ and Asp^WB^ can only be transiently visited by the “catalytic” proton as it redistributes over the protein on its way into the bulk solution. Such a mechanism of slow, solution-controlled hydrolysis may also operate, even in the presence of an activator, in those P-loop NTPases where Asp^WB^ and/or [Ser/Thr]^K+1^ were mutated [203, 268–275].

In a RasGAP-stimulated wild-type complex, the situation differs. Within the constricted catalytic site, the shortening of the H-bond with Ser17^K+1^ up to 2.4-2.5 Å would increase the proton affinity of Asp57^WB^ (see Fig. 9 and [261]), turning it into a deep trap for a proton from Ser17^K+1^. The anionic alkoxide of Ser17^K+1^ is thermodynamically much more favorable proton acceptor from W_cat_ than the pH-neutral solution. The same proton relays to β-phosphate when the latter is ready to accept it; there is no need to wait for a proton from the solution. As a result, hydrolysis is 10^5^ times faster than in the case of the lone Ras GTPase. Expectedly, the Asp57^WB^ to Asn mutation slowed the rate of the activated hydrolysis by Ras/RasGAP to the “intrinsic” level of hydrolysis observed in the absence of GAP [272].

By no way we state that γ-phosphate is banned for the “catalytic” proton from W_cat_. Distance measurements indicate that the catalytic proton can pass to [Ser/Thr]^K+1^ and Asp^WB^ also through the oxygen atom(s) of γ-phosphate in NTPases of all classes, which, to some extent, justifies numerous QM/MM models of water-mediated proton transfers from W_cat_ to γ-phosphate. These proton routes, however, involve “uphill” proton transfer from water to enzyme-bound triphosphate moiety with pK < 3.0 [197, 198]. Hence, these routes are thermodynamically very unfavorable and, therefore, very slow. Unfortunately, the QM modeling does not capture thermodynamic obstacles of this kind. It is this hindrance that may have prompted repeated independent emergence of γ-phosphate-bypassing proton pathways from W_cat_ to the [Ser/Thr]^K+1^ – Asp^WB^ pair, which we identified in TS-like structures of diverse P-loop NTPases, see Fig. 3-7, 11, 12.

We turned towards the strictly conserved Asp^WB^ and [Ser/Thr]^K+1^ as proton acceptors from W_cat_ after we realized that the “catalytic” Glu/Asp residues are provided by topologically diverse protein segments in different classes of P-loop NTPases see Fig. 1F, 4-6, 7A, Table 1, Section 2.2. and [9, 10, 32, 292]. Our analysis shows that proton pathways via “catalytic” Asp or Glu residues, emerged independently in kinases, SIMIBI NTPases, “medium-size” ASCE ATPases, and large F_1_/RecA-like ATPases, i.e. four times, at least.

This observation was unusual since catalytic bases are usually conserved within enzyme families. At the same time, the discovered amino acid residues that link W_cat_ to the coordination shell of Mg^2+^, although non-homologous, strongly resemble each other as well as the proton path in the PRC (cf Fig. 4-6, 7A with Fig. 10A), which suggested that they constitute proton pathways. In the scheme proposed here, these non-homologous, “catalytic” Asp/Glu residues, as well as [Ser/Thr]^SwI^ of TRAFAC class NTPases, are regarded as intermediate proton carriers, whereas thermodynamics of catalysis are determined by the difference in proton affinities between W_cat_ (or its analogs in kinases) and somewhat distantly located, but evolutionary conserved [Ser/Thr]^K+1^ – Asp^WB^ module, which serves as a *catalytic base*.

#### 3.2.5. Minimal mechanistic model of NTP hydrolysis by P-loop NTPases

Building on the comparative structure analysis of over 3100 catalytic sites presented here and in the accompanying article [41], as well as on the available experimental and theoretical data, we describe the activated catalysis typical for P-loop NTPases by a simple mechanistic model depicted in Fig. 13. We show the minimalistic version of the model that includes ubiquitous Walker A and Walker B motifs, Mg^2+^-NTP, a simple stimulator, such as a K^+^ ion or a Lys/Asn residue, and a few water molecules one of which serves as W_cat_. The model, however, can be easily expanded/adapted to fit distinct NTPase families by adding further stimulatory interactions, auxiliary and W_cat_-coordinating residues, as well as by replacing W_cat_ by other nucleophiles.

**Figure 13.**
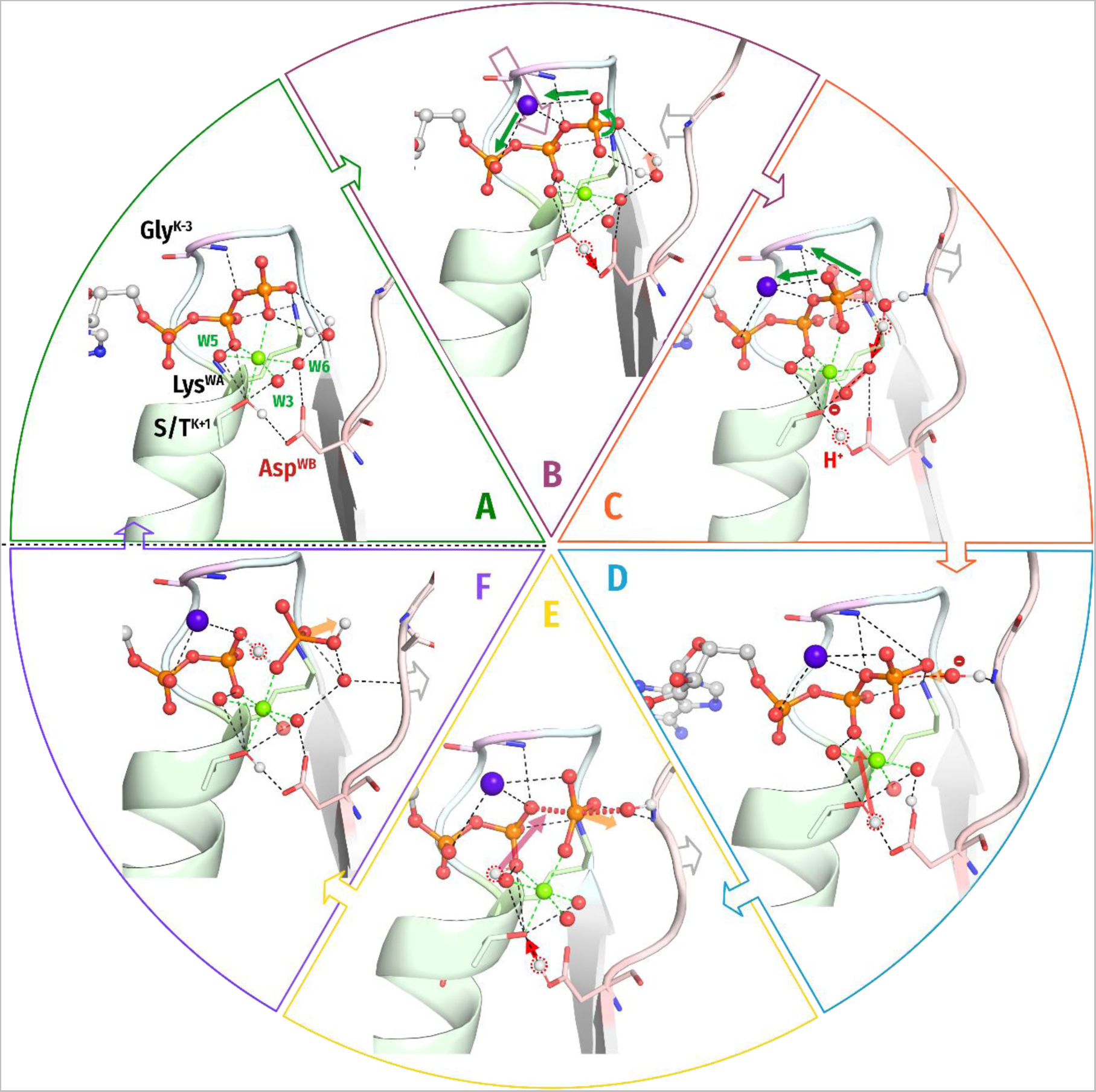
The common scheme of stimulated hydrolysis in P-loop NTPases. Empty purple arrow, movement of the stimulator (shown as a purple sphere); gray arrow, movement of the WB-crest; green arrows, twist and planarization of γ-phosphate; red arrows, proton displacements; orange arrows, detachment of Pi. The following crystal structures were used as templates: panels **A, B** – PDB ID 1FMW [139]; panels **C-E** – PDB ID 1VOM [95]; panel **F** – PDB ID 4PFP [52]. See the main text for a detailed explanation. A) A Mg-NTP complex binds to the Walker A and Walker B motifs; the binding energy is used to bring the NTP molecule into an elongated conformation with eclipsed β- and γ-phosphates and to close the catalytic site by surrounding the triphosphate chain by positively charged groups that are provided by the P-loop, WB-crest and, only in TRAFAC NTPases, Switch I loop. The accompanying protein conformational changes can power useful mechanical work. The catalytic site is further stabilized by the H-bond between Asp^WB^ and [Ser/Thr]^K+1^. The H-bond length is about 2.6-2.7 Å (Fig. 9); in this state Asp^WB^ is negatively charged. B) An exergonic interaction between the activating partner (another protein domain and/or an RNA/DNA molecule) and the WB-crest (i) shields and constricts the catalytic site, (ii) moves the WB-crest residues closer to the γ-phosphate, (iii) inserts the stimulator(s) next to the phosphate chain. The constriction of the catalytic site shortens the H-bond between [Ser/Thr]^K+1^ and Asp^WB^ to 2.4-2.5 Å, turning Asp^WB^ into a potent proton trap. In most cases, (one of) the stimulator(s) links the O^2A^ and O^3G^ atoms of the triphosphate (Fig. 1E, F, 3B-D, 4A-D, 5A-C, 6A) and twists γ-phosphate counterclockwise; the rotated γ-phosphate is stabilized by a new H-bond between O^2G^ and **HN**^K-3^. In other cases, the stimulators drag only γ-phosphate and, supposedly, twists it in some direction, see Fig. 3A, 5D, 6B-D. The interaction of stimulators with γ-phosphate (i) increases the electrophilicity of the P^G^ atom, (ii) weakens the O^3B^–P^G^ bond, (iii) promotes the transition of γ-phosphate to a more planar conformation, and (iv) inevitably affects the coordination of the Mg^2+^ ion by displacing the O^1G^ atom. The increase of local positive charge at [Ser/Thr]^K+1^ – after O^1G^ is moved aside by the stimulator – promotes the relocation of proton from [Ser/Thr]^K+1^ to Asp^WB^. C) The anionic [Ser/Thr]^K+1^ alkoxide withdraws a proton from the polarized W_cat_ molecule via intermediate proton carriers. Here we depicted the simplest proton route as envisioned for TRAFAC NTPases (see Fig. 12D-F and Section 3.2.2.4). More complex proton routes via W_cat_-coordinating Glu/Asp residues, as found in other classes of NTPases, are indicated by red dashed lines in Fig. 4-6 and differently shaded red arrows in Fig. 11, 12A-C. D) The resulting **OH^—^_cat_**, stabilized/polarized by its ligands, attacks the P^G^ atom. Although the simplified diagram in Fig. 13D shows only one stabilizing interaction of **OH^—^_cat_** with the **HN** group of the WB-crest residue, several ligands are usually involved in the stabilization, see Fig. 1E-F, 3-6. During this step, the proton stays on Asp^WB^. The formation of a covalent bond between **OH^—^_cat_** and P^G^ increases the planarization of γ-phosphate; its oxygen atoms repel the β-phosphate oxygen atoms, resulting in a lengthening of the O^3B^–P^G^ bond. With the inversion of γ-phosphate, increase in the O^3B^–P^G^ distance, and γ-phosphate moving away from β-phosphate, **HN**^K-3^ detaches from γ-phosphate and, together with Lys^WA^, Mg^2+^ and the stimulator, neutralizes the negative charge appearing on the O^3B^ atom, thereby lowering the activation barrier. In addition, the negative charge on β-phosphate attracts a proton from [Ser/Thr]^K+1^. E) The proton that comes from [Ser/Thr]^K+1^ forms a new short H-bond between β- and γ- phosphate [52, 201, 281], which further stabilizes the negative charge on the β-phosphate. The proton at Asp^WB^ relocates to [Ser/Thr]^K+1^. F) The H-bond between β- and γ-phosphate gradually dissociates as H_2_PO_4_ ^2-^ leaves the catalytic site. The departure of H_2_PO_4_^2-^ is an exergonic reaction that may be coupled to conformational changes, detachment of the activating partner from the WB-crest, and useful mechanical work.

According to the model, the catalytic transition proceeds in the following steps:

The simplified diagram in Figure 13 does not show the activation partner, as they are quite different in different families of P-loop NTPases. Accordingly, the exothermic interactions of the activator with the amino acid residues of the NTPase domain, which apparently result in the constriction of the catalytic site, are also not depicted. Some of them, however, are shown in Fig. S9, where the proposed general scheme is applied to myosin. Despite several sets of excellent crystal structures available [52, 91, 139, 293–297], the mechanism of ATP hydrolysis by myosin remains unclear [166]. Using the entire set of myosin crystal structures and the scheme in Fig. 13, we tried to reconstruct the catalytic cycle of myosin, as shown in Fig. S9 and described in its caption.

### 3.3. Conclusions and Outlook

Here and in the accompanying article [41] we applied the tools of evolutionary biophysics to elucidate the mechanisms of catalysis by P-loop NTPases. This approach allowed us not only to find support for previous MD simulation data on twisting γ-phosphate by the stimulatory moiety in P-loop GTPases [40, 163], but also, rather unexpectedly, to suggest a common catalytic mechanism for P-loop NTPases. The key elements of the proposed mechanism are the activation-induced, oppositely directed changes in proton affinity of H-bonded [Ser/Thr]^K+1^ and Asp^WB^, which convert the latter into a deep trap for a proton from [Ser/Thr]^K+1^ that, in turn, withdraws a proton from W_cat_ and converts it into a strong nucleophile OH^—^ . The suggested mechanism rationalizes the strict evolutionary conservation of [Ser/Thr]^K+1^ and Asp^WB^, neither of which has been ascribed a specific catalytic function so far.

Indeed, Walker A and Walker B motifs — together — contain only two strictly conserved residues capable of proton transfer, namely [Ser/Thr]^K+1^ and Asp^WB^. It is amazing that this H-bonded pair has not been considered in relation to proton transfer from W_cat_ up to now.

As argued elsewhere, the ability to provide strong acids and bases timely is important for enzyme catalysis [232]. Specifically, enzymes were suggested to generate strong bases or acids – precisely when required – by transiently altering the length of relevant H-bonds [261, 262, 298–304]. In the case of P-loop NTPases, the mechanism is quite simple: the free energies of (i) substrate binding and (ii) interaction with the activation partner are used to close and constrict the catalytic site, which shortens the H-bond between [Ser/Thr]^K+1^ and Asp^WB^ and levels their functional pK values. Eventually, after γ-phosphate is rotated by the stimulator, proton relocates from [Ser/Thr]^K+1^ to Asp^WB^ yielding a serine alkoxide as a strong nucleophile for W_cat_.

The still prevalent notion that chemically different catalytic bases (phosphates vs glutamates) are used in different classes of apparently homologous P-loop NTPases is bizarre, to say the least. Furthermore, neither phosphates nor glutamates are common as catalytic bases in the other families of phosphate transferases. In one of the most comprehensive reviews on their mechanisms, Cleland and Hengge even pointed out the oddness of anticipated catalytic bases in P-loop ATPases: “… more specifically, there must be a path for one proton of the attacking water molecule to reach a suitable acceptor. ATPases appear not to use general bases such as the aspartates usually found in kinase active sites “ (quoted from [60]). Our message is that P-loop NTPases are no exception, they do use aspartates as catalytic bases just like most other phosphate transferases.

The proposed mechanism of proton transfer from W_cat_ to Asp^WB^ via Mg^2+^-coordinating ligands brings the P-loop NTPases into the general context of other Mg-dependent hydrolases and transferases [57, 58, 60, 136, 266, 305, 306]. The full range of theoretical approaches developed for such enzymes can now be applied to P-loop NTPases. In particular, the relationship between changes in the Mg^2+^ coordination shell, proton affinity of Mg^2+^ ligands, and catalysis is the subject of QM/MM modeling in this field, see e.g. [307]. Also, Nemukhin and colleagues, have recently suggested proton transfer via the Mg^2+^ coordination shell in adenylate cyclase [306], which corroborates with the here proposed mechanism of proton transfer in P-loop NTPases. Therefore, we think that a new generation of QM/MM models of P-loop NTPases, which will include the entire Mg^2+^ coordination shell as well as proton transfer networks around Asp^WB^, will help to quantitatively describe their catalytic mechanisms.

Our tentative identification of the anionic [Ser/Thr]^K+1^ alkoxide as the proton acceptor from W_cat_ brings P-loop NTPases into another broad context of enzymes that generate a strong nucleophile by stripping a Ser/Thr residue of its proton [60, 164, 266, 267]. These are numerous families of serine proteases where, within apparently similar, but non-homologous “catalytic triads”, proton is transferred from the catalytic Ser to proton-accepting Asp via a histidine residue that serves as a von Grotthuss-type proton carrier. It is less known that the proton-accepting Asp residue of serine proteases is usually H-bonded to another conserved Thr/Ser residue [299, 308, 309]. Also, the catalytic Ser/Thr of eukaryotic protein kinases gives its proton to the conserved Asp [196, 310, 311], which, in turn, is also H-bonded to another conserved Ser/Thr residue.

In essence, we argue that P-loop NTPases use the same (bio)chemical strategies to produce a strong nucleophile as many other enzymes do; there is no reason to consider them different or special in this respect.

Notably, serine proteases, serine-threonine kinases, Mg-dependent hydrolases and transferases, as well as P-loop NTPases considered here, form the largest known enzyme families. Many of them use H-bonded [Asp/Glu]–[Ser/Thr] functional modules in different roles, which feature they share with membrane-embedded PRC and BR, see Fig. 10 and [247]. We believe that further search for structural features common to distinct enzyme families is a very promising task.

Last but not least, protonation of Asp and Glu residues can be traced in real time by IR spectroscopy [210, 230, 231, 239, 248, 255, 282, 285, 288]. Therefore, we believe that the proton shuttling between [Ser/Thr]^K+1^ and Asp^WB^ in various P-loop NTPases is worthy of being traced using photoactivated, caged ATP/GTP substrates and modern IR spectroscopy techniques (see, e.g., [39, 194, 282, 312]). Also, the position of a hydrogen atom within a short H-bond can be identified by complementary X-ray and neutron crystallographic structure determination [302–304]. Investigation of P-loop NTPases by these methods will shed new light on enzyme mechanisms.

## 4. Methods

### Manual comparative structure analysis

For each class of P-loop NTPases, representative structures were manually selected from the Protein Data Bank (PDB) at www.rcsb.org [71, 72] based on literature data and structure availability.

### Cumulative computational structure analysis

As for the analysis in the accompanying paper [41], structures were selected among those PDB entries that matched the following criteria: (1)assigned to InterPro record IPR027417; (2) contained an ATP/GTP molecule, or non-hydrolyzable analog of NTP, or a transition-state analog; (3) contained at least one Mg^2+^, Mn^2+^ or Ca^2+^ ion; (4) resolution of 5 Å or higher, if applicable. Proteins were assigned to major classes of P-loop NTPases according to membership in Pfam families (ftp://ftp.ebi.ac.uk/pub/databases/Pfam/mappings/pdb_pfam_mapping.txt) [9, 10], see the details in the accompanying paper [41]. This search yielded 1474 PDB structures with 3666 catalytic sites in them.

We checked the reliability of each catalytic site by assessing the integrity of the NTP analog/presence of γ-phosphate mimic, presence of a P-loop motif Lys residue within 5Å of the β-phosphate. This analysis yielded 3136 complexes in 1383 structures with various substrates: ATP and GTP, their nonhydrolyzable analogs, and ADP/GDP molecules associated with γ-phosphate-mimicking moieties, see Fig. 14 in the accompanying paper [41].

Also as a quality control, we have measured distances from [Ser/Thr]^K+1^ to Mg^2+^, to ensure the correct binding of the Mg^2+^ and general reliability of the structure resolution at the binding site (i.e, very long distance would indicate a disturbed catalytic site or resolution at the site that is insufficient for our purposes of comparative analysis), and from [Asp/Glu]^WB^ to Mg^2+^, to identify cases of direct coordination of Mg^2+^ by the acidic residue (short distances) or disassembled binding sites (long distances).In addition, we examined the presence of interactions of phosphate chain with NH^K-3^ group and positively charged moieties in those sites that passed the quality control (the details of analysis are described in the accompanying paper [41] and depicted in Fig. 14 of it.)

Putative [Asp/Glu]^WB^ residue was identified as follows: distances from all Asp and Glu residues to [Ser/Thr]^K+1^ were measured, and the closest residue that was preceded by at least three non-ionizable residues (Glu, Asp, Ser, Thr, Tyr, Lys, Arg and His were considered as ionizable) was chosen as the partner of [Ser/Thr]^K+1^. If this failed, a closest Asp/Glu was selected without hydrophobicity check (data available in Table S1). Residues located further than 5Å were not considered. In those few cases where [Asp/Glu]^WB^ residue is indicated in the Table S1 as “absent at the threshold distance”, the catalytic site is likely to be fully “open”. We have not checked all these cases manually.

### Visualization

Structure superposition, manual distance measurements, manual inspection and structures visualization were performed by Mol* Viewer [313] and PyMol v2.5.0 [314].

### Data availability

Descriptions of each binding site are available in Table S1. Scripts used to generate and annotate the data and quickly visualize selected sites listed in Table S1 are available from github.com/servalli/pyploop.

## Acknowledgements

Very fruitful discussions with Drs. D.A. Cherepanov, M.Y. Galperin, D.N. Frick, K. Gerwert, A. V. Golovin, Y.Kalaidzidis, E.V. Koonin, A. Lupas, B.H. Meier, N. Voskoboynikova, T. Wiegand and M. Zereal are highly appreciated. We also are thankful to Alexander Mulkidzhanyan for his help during the initial stage of the project. The research was supported by DFG, DAAD, and the Osnabrueck University (the EvoCell Program and Open Access Publishing Fund).

## [7] Supplementary Materials

### Supplementary Figures

**Supplementary File 1.** Consideration of tentative activation mechanisms in STAND, KAP, VirD/PilT and FtsK-HerA classes of P-loop ATPases

**Supplementary Table 1.** Results of the computational analysis of all available structures of the P-loop proteins in complex with NTPs and NTP-like molecules. The Microsoft Excel table contains the list with characteristics of all analyzed structures, together with key functional residues of the Walker A and Walker B motifs, Arg, Lys, and Asn fingers, as well as distances from (1) the respective atoms of NTPs/their analogs to the K-3 residues and Arg/Lys fingers, as well as (2) from [Asp/Glu]^WB^ to [Ser/Thr]^K+1^. Each row contains the data for one catalytic site in one structure. Catalytic sites containing “properly bound” NDP:AlF_4_^-^ complexes that we deemed to be reliable TS-analogs (see Table S3 in the accompanying paper [41] for detailed description of AlF_4_^-^ and Mg^2+^ binding in such structures) are marked with “y” or “*” in column “site rel”; they are colored green. The sites with “unproperly bound” NDP:AlF_4_^-^ complexes are colored pink. All columns present in the Sheet A of the Table S1 (data) are described in the Sheet B.

**Supplementary Table 2.** Analysis of the bonding pattern within the catalytic sites of the RhoA GTPase and F_1_-ATPase as a function of the TS analog bound.

## Supplementary Figures to the manuscript

**Figure S1.**
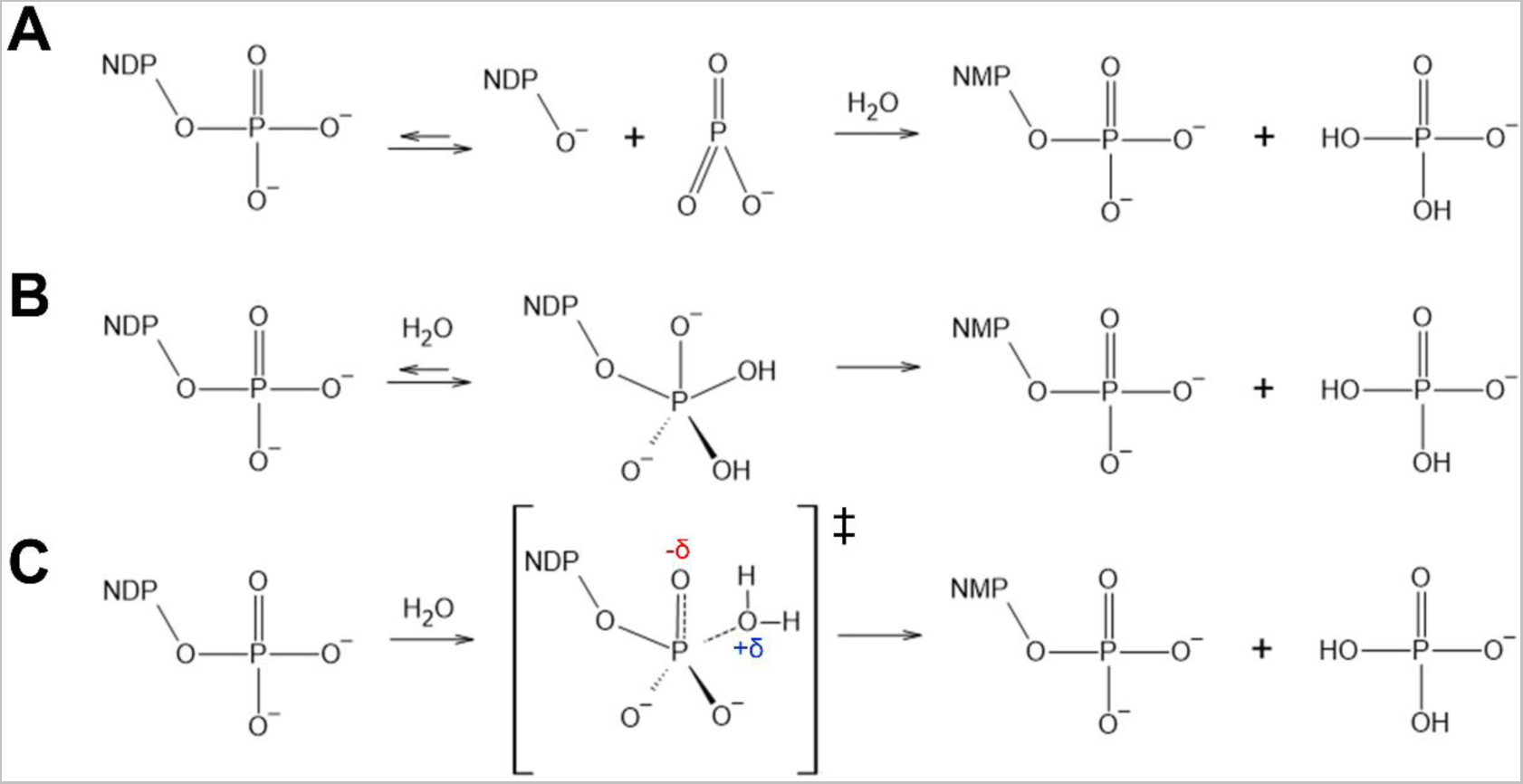
Tentative mechanisms of NTP hydrolysis. Dissociative (**A**), associative (**B**), and concerted pathways (**C**). See the main text for further details and references.

**Figure S2.**
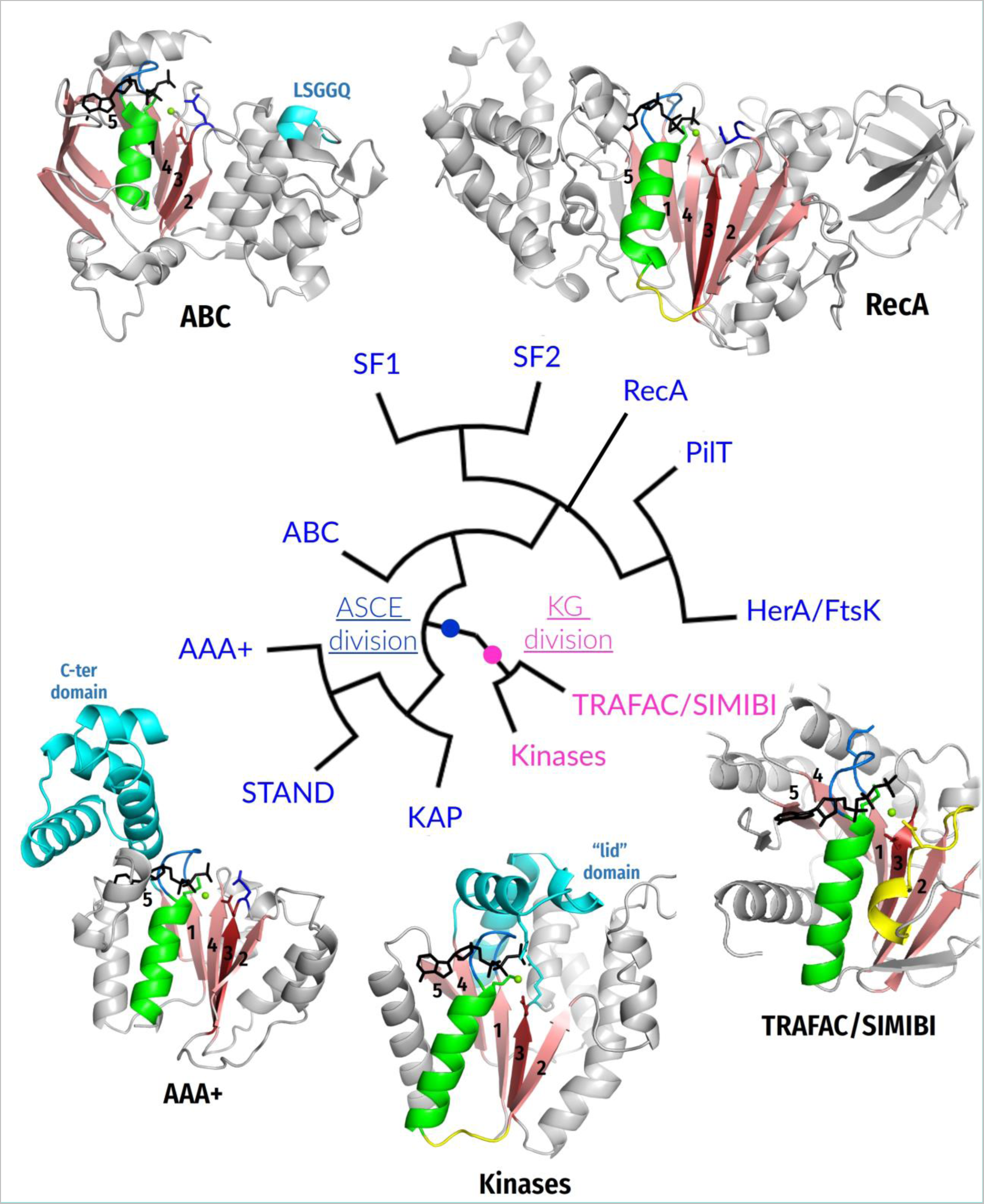
Cladogram of higher-order relationships between major divisions/classes of P-loop NTPases with depicted typical structures. The cladogram (as modified from [1]) shows the two major divisions: the <u>k</u>inase-<u>G</u>TPase (KG) division and the ASCE (<u>a</u>dditional <u>s</u>trand, <u>c</u>atalytic <u>E</u>) division. The β-strands forming the cores of P-loop domains are numbered in a traditional way [1, 2]. These β-strands are colored pink, the P-loop is shown in blue, the following α_1_-helix is shown in green, the K-loop/Switch I in TRAFAC class NTPases as well as the corresponding loops in other NTPases are colored in yellow, NTP analogs are shown as black sticks, Mg^2+^ ions are shown as green spheres, the rest of proteins is shown as gray cartoons. The reference residues of Walker A and Walker B motifs (lysine (K) and aspartate (D), respectively), as well as the catalytic glutamate (E) in ASCE NTPases are shown as sticks. The following reference crystal structures are depicted: ABC – antibacterial peptide ABC transporter McjD of *Escherichia coli*, PDB ID 5EG1 [3]; RecA/F_1_ –F_1_-ATPase of *Caldalaklibacillus thermarum*, PDB ID 5HKK [4]; AAA+ –clamp loader γ-subunit *E. coli*, PDB ID 1NJG [5]; Kinases –thymidylate kinase of *E. coli*, PDB ID 4TMK [6]; TRAFAC/SIMIBI – nitrogenase ATPase subunit of *Azotobacter vinelandii*, PDB ID 4WZB [7].

**Figure S3.**
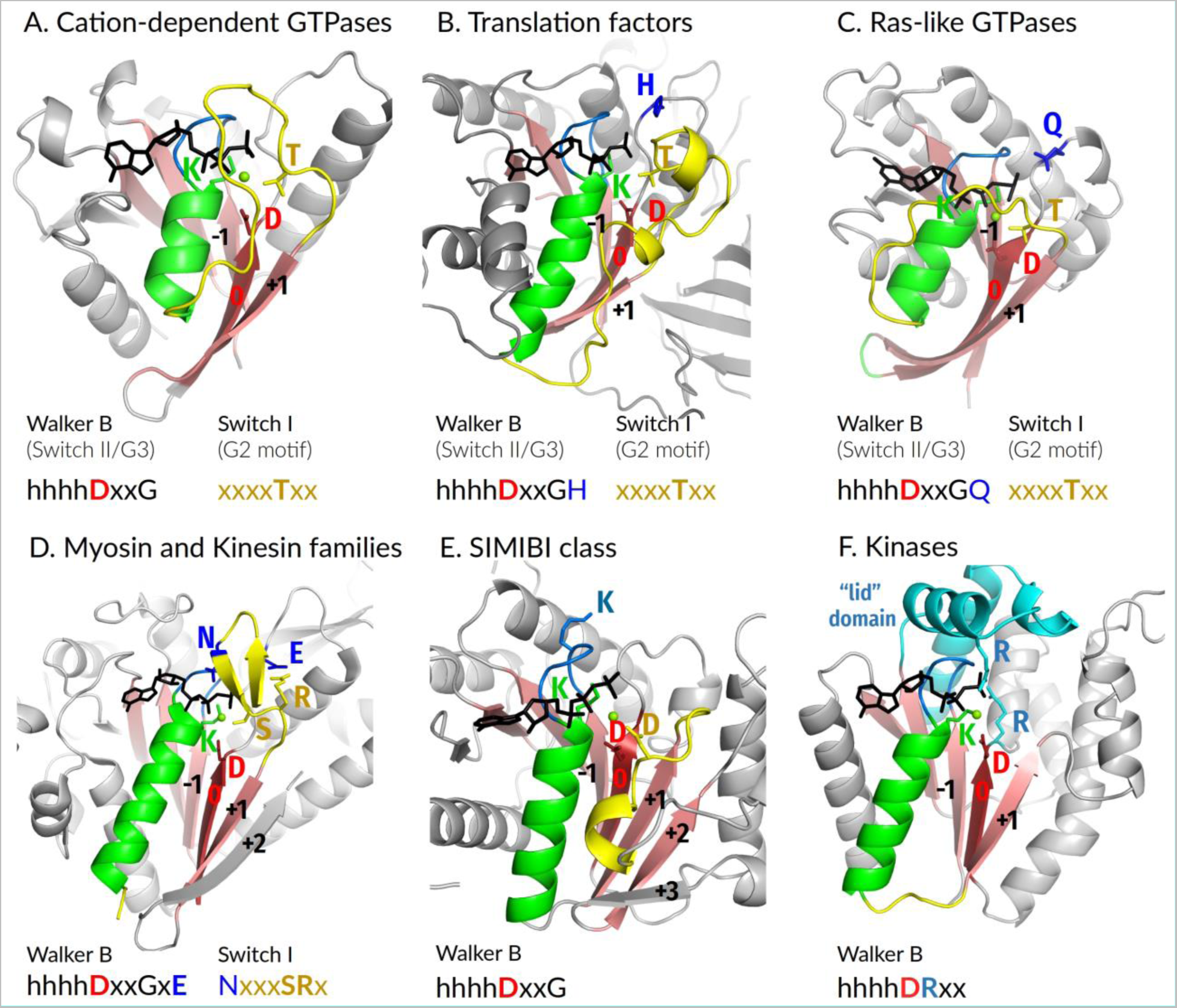
P-loop domain organization and conserved motifs in P-loop NTPases of the Kinase-GTPase division. The β-strands forming the cores of P-loop domains are numbered relative to the Walker B motif-containing WB-strand as suggested in the main text. Color code as in Fig. S2. The following crystal structures are depicted: A, MnmE GTPase of *E. coli*, PDB ID 2GJ8 [8]; B, Elongation factor EF-Tu of *Thermus thermophilus* HB8, PDB ID 4V5L [9]; C, Human Ras-GTPase, PDB ID 1WQ1, [10]; D, Myosin II of *Dictyostelium discoideum*, PDB ID 1W9L; E, nitrogenase ATPase subunit of *Azotobacter vinelandii*, PDB ID 4WZB [7]; F, thymidylate kinase of *E. coli*, PDB ID 4TMK [6].

**Figure S4.**
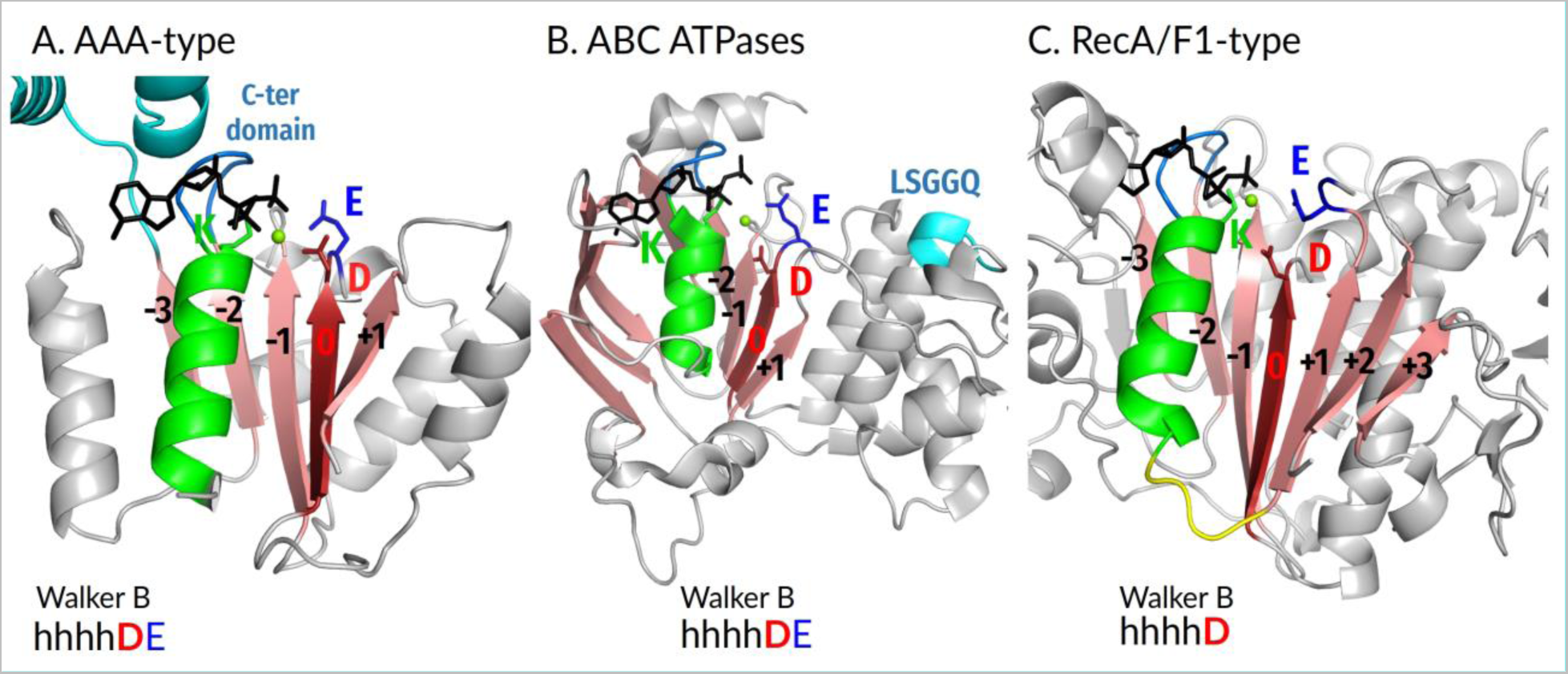
P-loop domain organization and conserved motifs in P-loop NTPases of the ASCE division. The β-strands forming the cores of P-loop domains are numbered relative to the Walker B motif-containing WB-strand as suggested in the main text. Color code as in Fig. S2. The following crystal structures are depicted: A, AAA+ Clamp Loader Gamma Subunit of *E. coli*, PDB ID 1NJG [5]; B, antibacterial peptide ABC transporter McjD of *E.coli*, PDB ID 5EG1 [3]; C, F_1_-ATPase of *Caldalaklibacillus thermarum*, PDB ID 5HKK [4].

**Figure S5.**
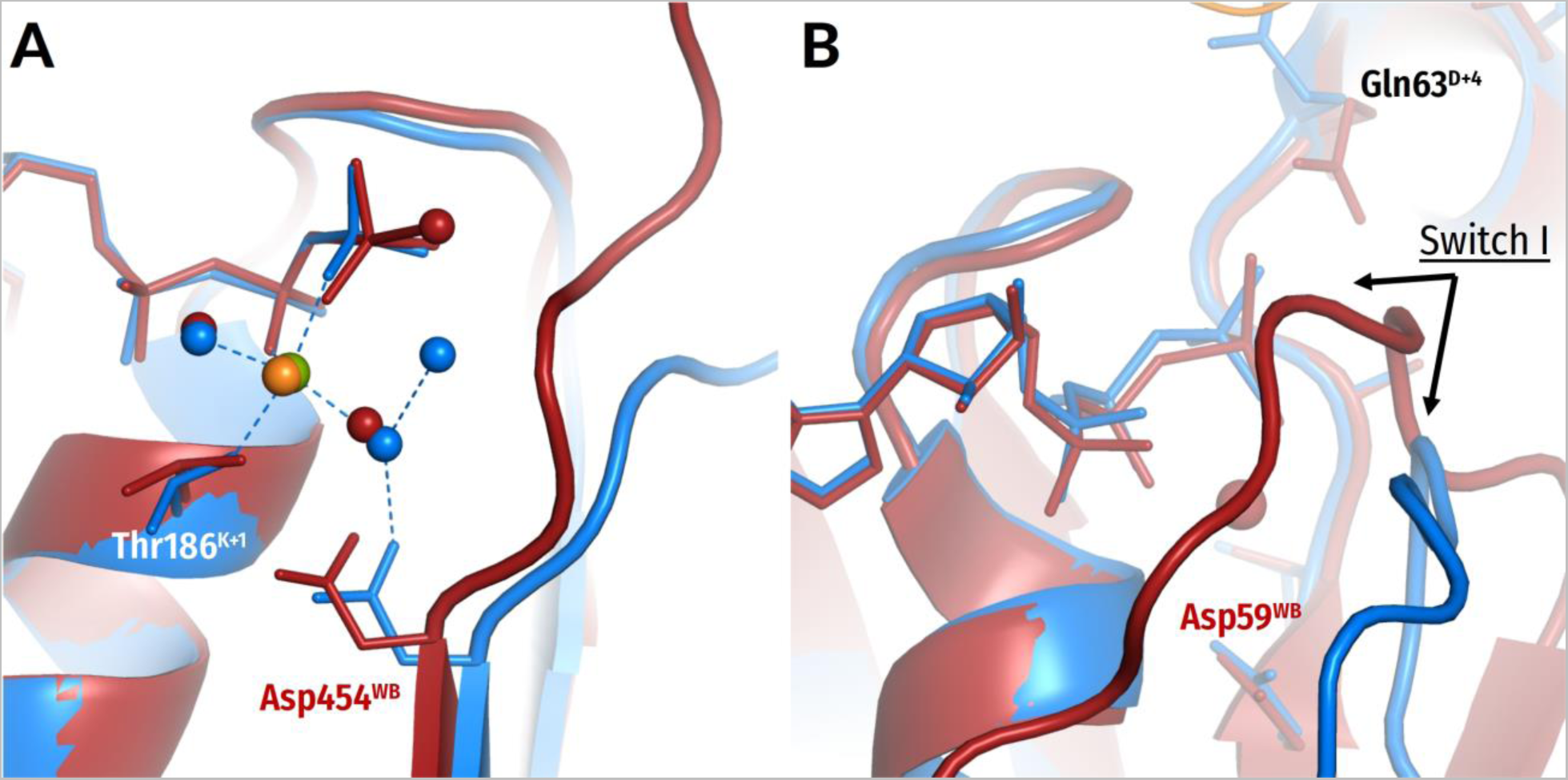
Structures of same P-loop NTPases with transition state analogs and substrate/substrate analogs bound, respectively. The structures with bound substrate or substrate analogs are shown in blue with their Mg^2+^ ions in green; the structures with transition state analogs bound are shown in dark red with their Mg^2+^ ions in orange. Cations and water molecules are shown as spheres. The following crystal structures are depicted**: A,** Myosin II from *Dictyostelium discoideum* in complex with a bound ATP molecule (blue, PDB ID 1FMW [11]) and with a transition state analog ADP:VO_4_^-^ bound (dark red, PDB ID 1VOM [12]); **B,** Ras-like GTPase RhoA with a bound non-hydrolyzable substrate analog GNP (blue, PDB 6V6M, [13]) and a complex of Rho with RhoGAP and a transition state analog GDP:MgF_3_^—^ (dark red, ODB 1OW3, [14]).

**Fig. S6.**
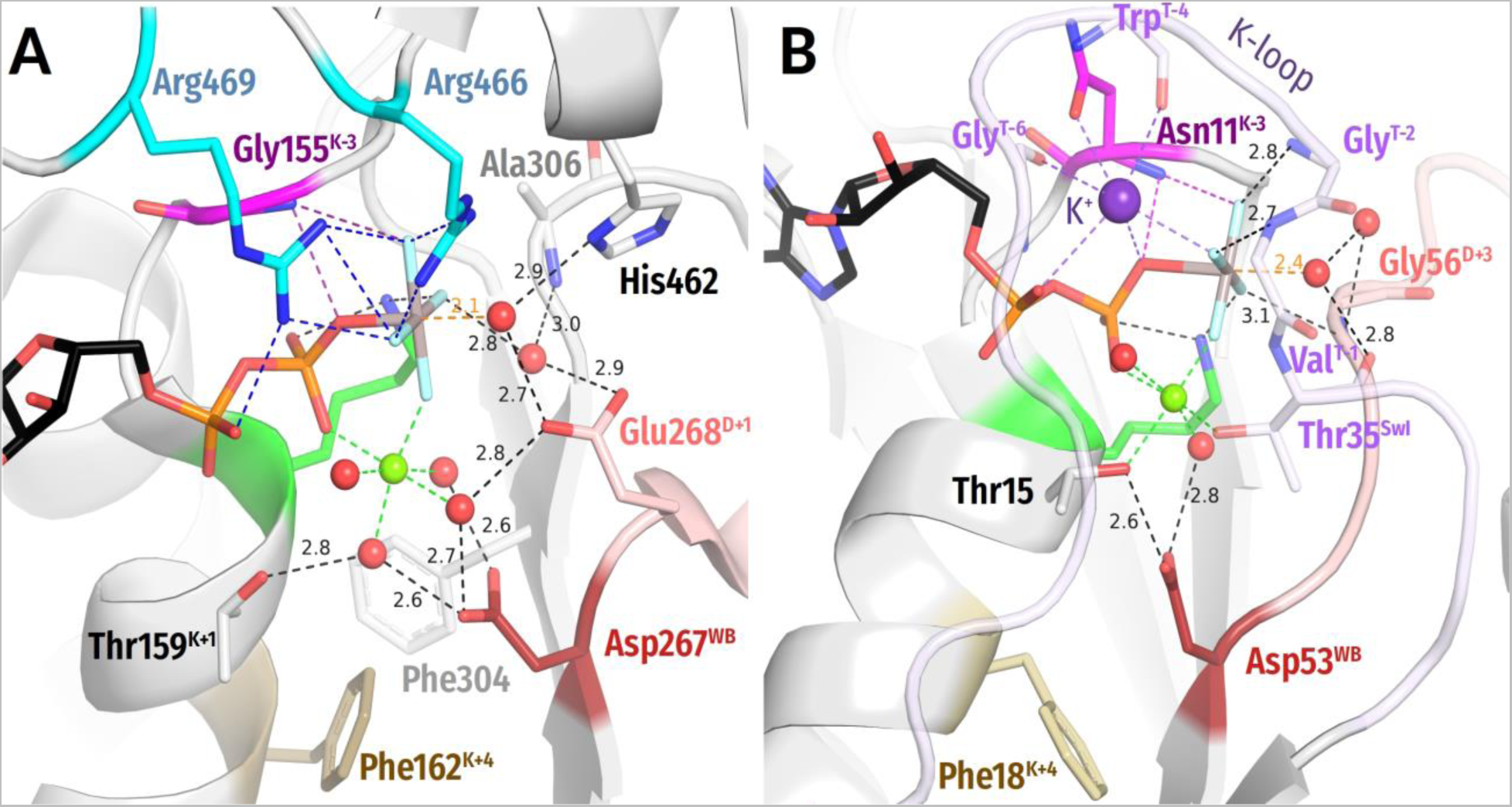
Interference of Phe^K+4^ with the Asp^WB^ — [Ser/Thr]^K+1^ bonding. **A.** The only detected structure with properly bound AlF_4_^—^ and, still, long distance between Asp^WB^ and [Ser/Thr]^K+1^ is the mitochondrial yeast DEAD box protein Mss116p where this distance is about 4 Å, see Panel A and (PDB 3I62 [15]). The structure analysis has shown that the formation of the H-bond between Asp^WB^ and [Ser/Thr]^K+1^ is prevented by the Phe162^K+4^ residue. It interacts with Phe304 of the WB-1 strand so that these two residues intercalate between Walker A and Walker B motifs and prevent the formation of the H-bond (Panel A). Remarkably, the Thr^K+1^ is not directly coordinating Mg^2+^, pointing to a possibility of a crystallization artefact or an inactive configuration of the binding site with the Phe^K+4^ acting as a regulatory switch. **B.** A phenylalanine residue is not unique in K+4 position, it is also found in the well-studied K^+^- stimulated GTPase FeoB. In its GDP:AlF_4_^−^-containing structure (PDB ID 7BWV), the homologous Phe18^K+1^ is, however, in a different rotameric position and do not interfere with the Asp^WB^ – [Ser/Thr]^K+1^ bond (Panel B). Notably, in this structure the residue is not involved in a π-π interaction with another phenylalanine residue that would orient it towards the Asp^WB^ —Thr^K+1^ bond (cf Panel A).

**Figure S7.**
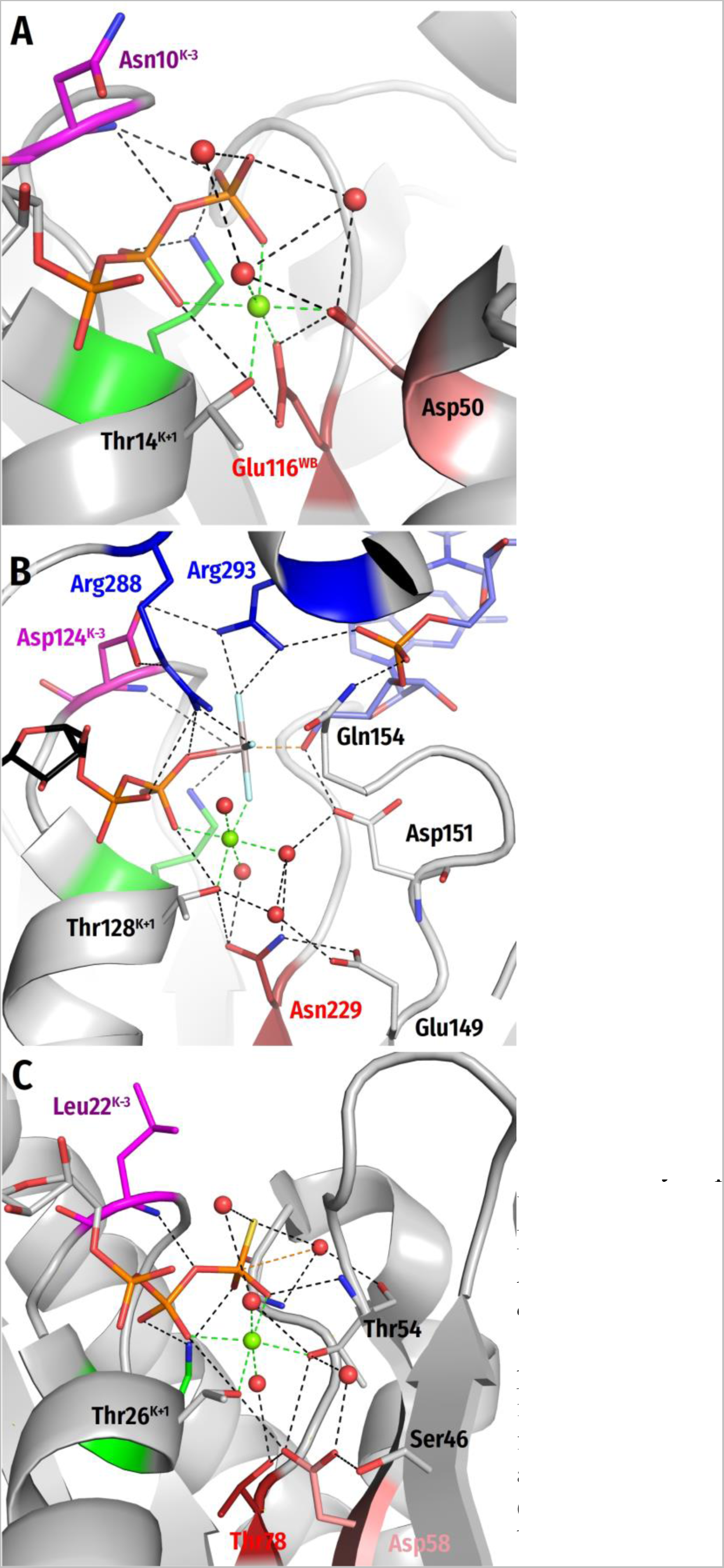
Deviations from a typical Walker B motif. Color code as in Fig. 1E,F. A. In a few cases, which are considered in detail in [16], the long side chain of Glu^WB^ reaches Mg^2+^ and directly coordinates it, forming also a H-bond with [Ser/Thr]^K+1^, as shown in Panel A for the Dethiobiotin synthetase (BioD) from *Helicobacter pylori* complexed with GTP (PDB 3QXJ [17]). In this enzyme, the γ-phosphate group is transferred on the carboxy group of 7,8-diaminononanoate, which is likely to be deprotonated at neutral pH, so that no proton trapping is needed. In a few more conventional cases, the Glu^WB^ residue, similarly to Asp^WB^, does not reach Mg^2+^, interacts with Mg^2+^-coordinating water and makes an H-bond with [Ser/Thr]^K+1^, see [16]. B. An Asn229 residue links W3, W6 and Thr131^K+1^ in the wild-type polynucleotide kinase of *Caenorhabditis elegans,* see Panel B for its structure with bound ssRNA dinucleotide GC and ADP:AlF_4_^-^ (PDB ID 4OI1 and [16, 18]). In this case, however, the adjacent WB+1 strand has a ^149^ELD^151^ triad at its C-cap. In the TS-like AlF_4_^-^- containing structure, the Glu149 residue is connected via a bridging water molecule to Thr131^K+1^ and to W3, whereas Asp151 makes a short H-bond (2.48 Å) with the hydroxyl of the RNA ribose. Deprotonation of this hydroxyl yields the nucleophile that attacks the γ-phosphate. We believe that the functions of Asp^WB^ are divided between Asn229 and Glu149 in this polynucleotide kinase. The Asn229^WB^ residue serves as a structural linker to the Walker A motif, whereas Glu149 serves as a trap for the proton that is taken by Asp151 from W_cat_. While the same Asn-Glu-Asp triad is found in other Metazoa enzymes, Asp substitutes for Asn229 and Asn substitutes for Glu149 in *Saccharomyces cerevisiae*, *Schizosaccharomyces pombe*, and *Candida albicans.* Hence, in evolutionarily primary primitive microorganisms, a single Asp^WB^ appears to interact with W6 and Thr^K+1^ and to obtain a proton from the catalytic Asp residue at the C-cap of the adjacent WB+1 strand; in the *C. elegans* structure (PDB ID 4OI1 [18]), Asn229 and Asp 151 are linked by W6. The “primordial” dyad of aspartates is functionally similar but structurally distinct from the Asp^WB^L^D+1^Asp^D+2^ aspartate dyads found in some other kinase families, see Fig. 4D and [19]. Generally, the overall diversity of proton routes within kinases [19] might reflect the diversity of second substrates in these enzymes, see Section 2.2. C. A natural mutation in an otherwise typical small TRAFAC class GTPase MglA changed Asp^WB^ to Thr. The structure of MglA GTPase of *Myxococcus xanthus* with a bound non-hydrolizable analog of GTP is shown (PDB ID: 6IZW and [20]). The enzyme retained its function owing to the appearance of an Asp residue at the N-terminus of the adjacent antiparallel WB+1 strand [16]. The resulting topology of the catalytic site is typical for P-loop NTPases.

**Fig. S8.**
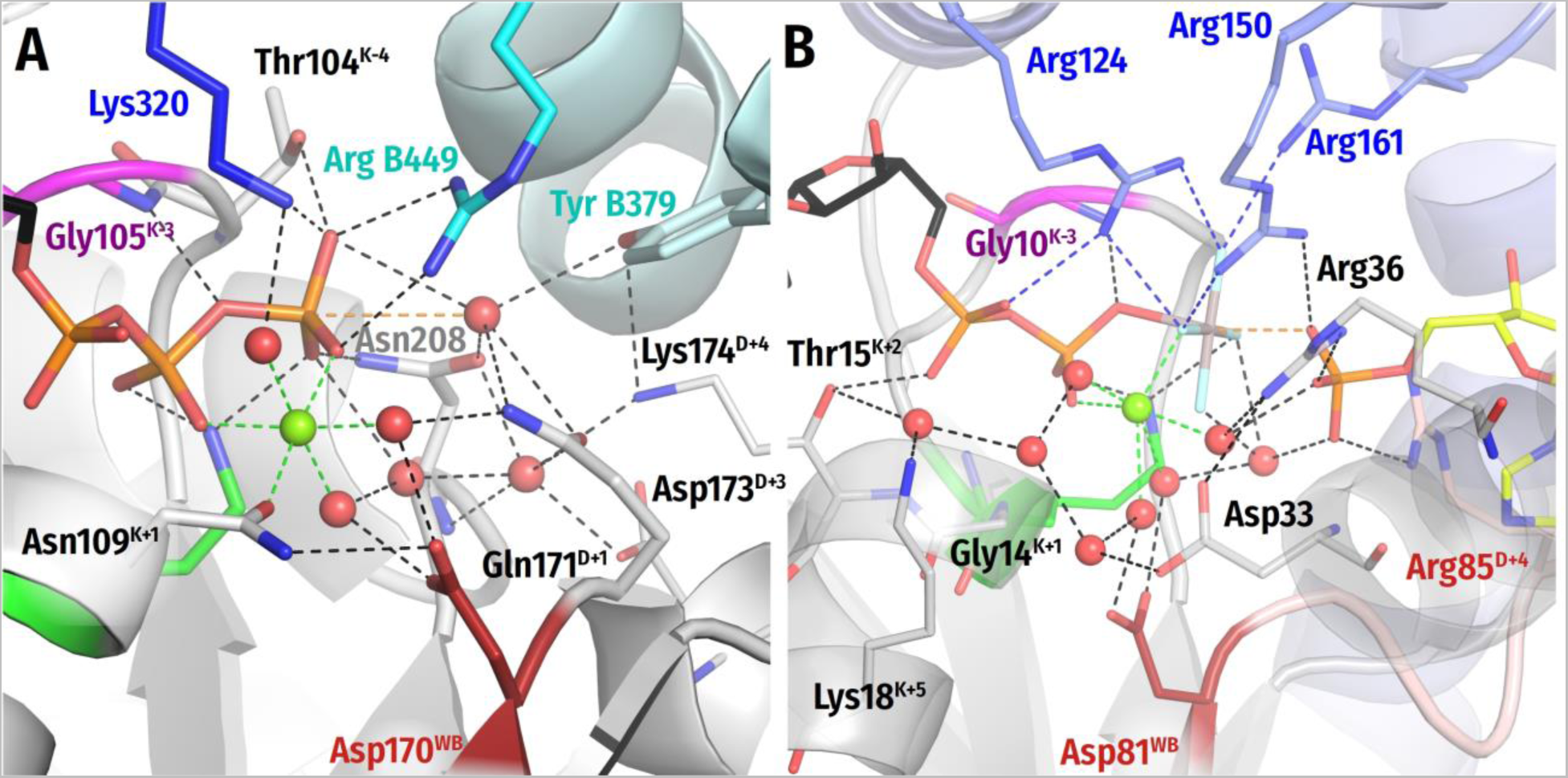
Residues other than [Ser/Thr] in the K+1 position of the Walker A motif. Colors as in Fig. 3-6 in the main text. Lys^WA^ shown in green, Asp^WB^ in dark red, adjacent monomer and its residues in cyan, Arg and Lys finger from the same polypeptide chain in blue. A, an asparagine residue replaces [Ser/Thr]^K+1^ in such distantly related AAA+ ATPases as torsinA of metazoa and DnaC helicase loader of some enterobacteria including *E. coli* [16, 21, 22]. In addition, the potentially W_cat_-stabilizing “catalytic” [Glu/Asp]^D+1^ – ubiquitous in AAA+ ATPases [23] – is replaced by Asn^D+1^ in torsinA (but not in DnaC). The ATPase activity of both these proteins are very low, so it is not yet clear if they are true ATPases or simply work as switches that change their conformation depending on whether ATP or ADP is bound in the catalytic site [21, 22, 24, 25]. Shown is the high-resolution ATP-containing structure of human torsinA in complex with its activator LULL1 (PDB ID 5J1S [26]). It can be seen that Asn109^K+1^ makes the canonic H-bond with Asp170^WB^ which could increase the proton affinity of the latter. While Asn171^D+1^ cannot transfer a proton from W_cat_, a water chain – similar to one observed in TRAFAC class NTPases (Fig. 7B, 12D-F) – connects W_23_ with Asp170^WB^ via the Mg^2+^-coordinating W6 molecule. In AAA+ NTPases, a Lys/Arg residue always interacts with γ-phosphate and, supposedly, W_cat_ (see section 2.2.2.2, Fig. 5A,B, and [27, 28]). In torsinA, this is the Arg449 residue of LULL1. In the absence of negatively charged Asp/Glu^D+1^, the interaction of a positively charged Arg449 with W_cat_, by dramatically decreasing its proton affinity, may trigger proton transfer from W_cat_, via W6, to Asp170^WD^. Subsequently, the proton can move to β-phosphate via W6. Hence, the structure is compatible with involvement of Asp170^WB^ as a proton trap in torsinA and could explain its, albeit slow, ATPase activity. B, Adenylate kinase of *Aquifex aeolicus* ([PDB 3SR0] [29]), see the main text.

**Figure S9.**
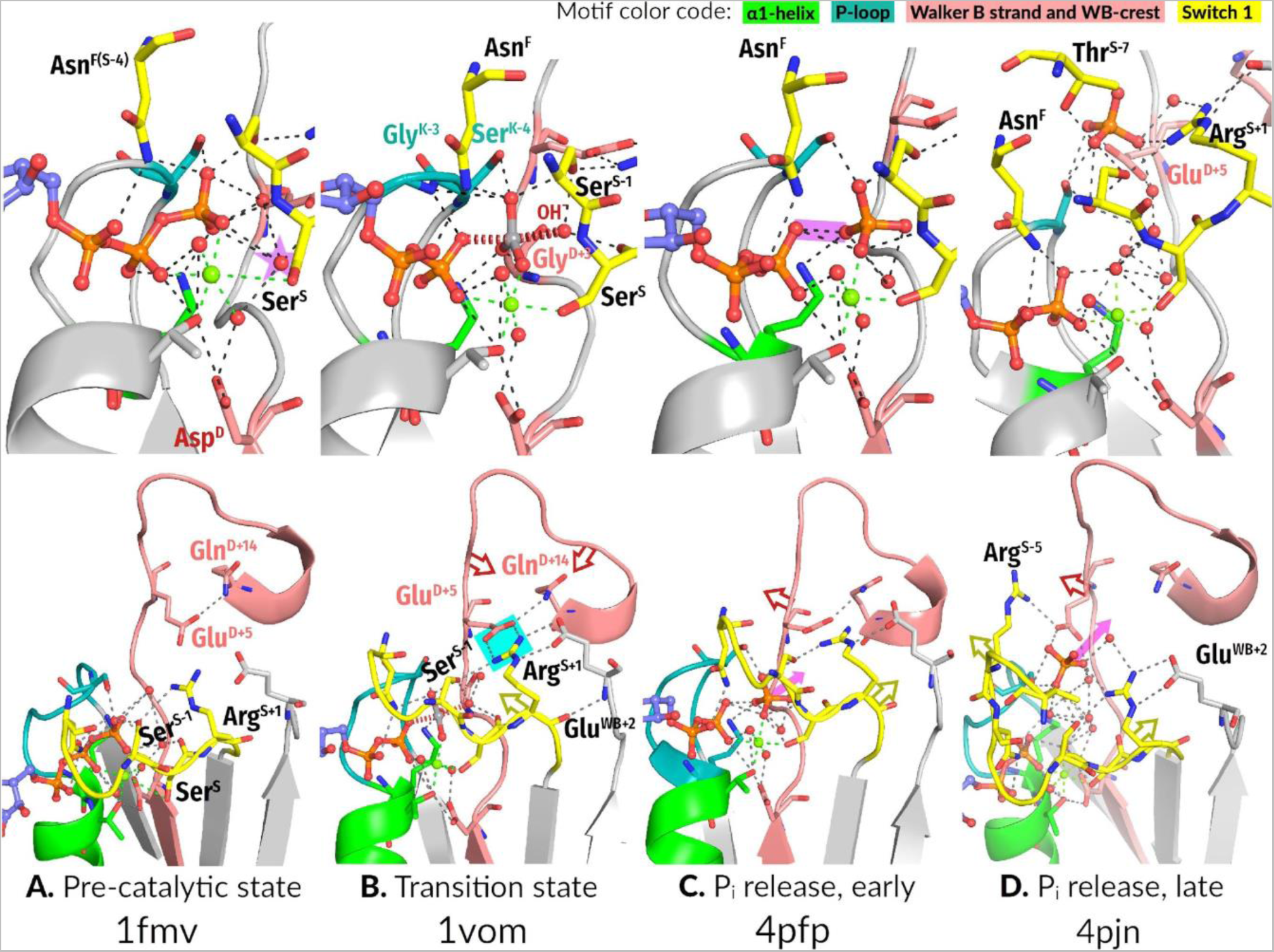
Catalytic cycle of myosin as inferred from the available set of crystal structures. The top and bottom rows show similar structures differently zoomed. Given that myosin is one of the few P-loop NTPases with a set of high-quality crystal structures including those mimicking the pre-transition, transition, and several post-transition states (see also Supplementary Table 1), we tentatively reconstruct the catalytic cycle of myosin as follows: **A)** (top) In the substrate-bound state (ATP-soaked myosin crystals, PDB ID 1FMW [11]), the catalytic pocket is not constricted, and the proton pathway from the would-to-be W_cat_ in the W_12_ position (highlighted with a star) to Asp^WB^ (via W6) is clearly seen. (bottom) In this state, Glu^D+5^ of WB-crest and Arg^S+1^ of Switch I motif are remote from each other, and each residue is only interacting with its neighbors in the polypeptide chain. Glu^D+5^ is forming an H-bond with Gln^D+14^, while Arg^S+1^ is in contact with Ser^S-1^ (bottom). **B)** (top) In the ADP:VO_4_^-^-containing structure, which mimics the transition state (PDB 1VOM [12]), the site is constricted, and W_12_, after giving its proton via W6 to Asp^WB^, is pushed upwards into the attack position of **OH^-^_cat_**, here mimicked by the oxygen atom of VO_4_^-^ . In this position, **OH^-^_cat_** is stabilized by the side chain of Ser^S-1^, **CO** of Ser^S^, and **HN** of Gly^D+3^. Also in this state, the Asn finger comes closer to the phosphate chain and interacts with α-phosphate and one of the oxygen atoms of VO_4_^-^, another oxygen atom of VO_4_^-^ interacts with Gly^K-3^ and with the side chain of Ser^K-4^. The WB-crest is brought closer to the catalytic site: an oxygen atom of VO_4_^-^ is coordinated by Gly^D+3^. Here, the distance between **OH^-^_cat_** and W6 is 4.8 Å, which, supposedly, precludes the proton at Asp^WB^ from returning to W_cat_. The proton, however, can pass to β- phosphate; all the tentative proton transfer steps on the way to β-phosphate are of 2.9 Å. (bottom) An extensive salt bridge network is formed between WB-crest, Switch I and the adjoining domain. Arg^S+1^ interacts with Ser^S-1^ and Glu^D+5^ (bonds highlighted in bright cyan), whereas subsequent residues of the WB-crest are contacting the adjoining protein domain. **C and D**) In the two subsequent post-transition states (mimicked by differently P_i_-soaked ADP-containing myosin structures with PDB IDs 4PFP and 4PJN [30]), catalytic site opens up: the release of H_2_PO_4_^2-^ is coupled with breaking of salt bridges between WB-crest and Switch I. Hence the exergonic release of P_i_ drives the opening of the catalytic pocket, which, in turn, is coupled to the mechanical movement. The Asn finger no longer interacts with the outgoing H_2_PO_4_ ^2-^ moiety. In the early post-transition state the proton is already engaged in the H-bond between O^3B^ and O^3G^ (C, top, the H-bond is highlighted in purple), while in the late post-transition state (D), hydrogen-bonded networks are further rearranged: the outgoing H_2_PO_4_^2-^ no longer contacts residues involved in P_γ_ coordination during the previous steps (D, bottom); H_2_PO_4_ ^2-^ gets directly engages with Glu^D+5^ and Arg^S+1^ pulling apart the WB-crest and Switch I (D, bottom).

## Supplementary File 1 to the manuscript

### Classes of P-loop NTPases with ambiguous mechanisms of stimulation and Wcat stabilization

For several large families and even classes of P-loop NTPases, there are no structures with full-fledged catalytic sites containing TS-analogs. Only assumptions can be made on their stimulatory moieties and environments of W_cat_.

### Kinase-GTPase Division

***Septins and septin-like proteins* of the TRAFAC class** appear to be stimulated by His residues, which are reciprocally inserted into the catalytic sites upon dimerization of the NTPase domains; the dimerization, supposedly, is induced by the interaction with the activating partner(s) [1, 2]. In septins, the NE2 group of the stimulatory His residue appears to enter the AG site and to link the O^2A^ and O^3G^ atoms [2]. For septins, no TS-like structures are available. Still, in the GNP-containing structure of human septin 12 (PDB ID 6MQ9 [3]) the water molecules next to γ-phosphate interact with NH^D+3^ and residues of the Switch I motif.

### ASCE Division

**STAND (*<u>s</u>ignal transduction <u>A</u>TPases with <u>n</u>umerous <u>d</u>omains)* ATPases** unite apoptotic ATPases (animal apoptosis regulators CED4/Apaf-1, plant disease resistance proteins, and bacterial AfsR-like transcription regulators) and the ATPases of the NACHT superfamily, which consists of the animal disease response ATPases such as CARD4, the NAIP proteins and other ATPases [4]. The Walker B motif of these ATPases contains two consecutive acidic residues [4]. In addition to the P-loop domain, most proteins of the STAND class carry a unique HETHS domain - the third helical domain, which is present only in this class of proteins [4]. Since some proteins of the STAND class have a conserved Arg residue within the P-loop domain, analogous to the Arg finger of AAA+ ATPases, it was suggested that their activation mechanism is similar to that of AAA+ ATPases [4]. However, available structures of Apaf-1 (Apoptotic protease activating factor 1) suggest a direct involvement of the HETHS domain in the ATP hydrolysis. Structures of Apaf-1 are available for the two states of the protein: the ADP-bound (no Mg^2+^), “inactive” state (PDB ID 3SFZ [5]), and the “active” state, with Mg-ATP and cytochrome *c* bound (PDB ID 3JBT [6]). It is important to note, that Apaf-1 does not have ATPase activity, so that “active” and “inactive” states differ in the ability of the protein to oligomerize into an apoptosome. In the ATP-bound state, a Tyr residue forms an H-bond with the γ-phosphate (Fig. SF1_1B). In the ADP-bound state, a His residue from the HETHS domain attains a position next to α and β-phosphates and from the same side as Arg/Lys fingers in other P-loop NTPases (Fig. SF1_1B). This His residue is conserved in the majority of STAND proteins [4]. Specifically, a His residue is seen in the similar position in the ADP-containing STAND protein with a tetratricopeptide repeat sensor PH0952 from *Pyrococcus horikoshii* (see Fig. SF1_1C, PDB ID 6MFV [7]). STAND proteins appear to be special; unlike other P-loop NTPases, they convert from a closed form occluding an ADP molecule to an ATP-bound open form prone to multimerize [7]. Hence, Apaf-1 and its homologs cannot serve as a reliable representative for the whole class, especially considering the diversity of domain architectures and the presence of a conserved Arg residue in the P-loop domains of some families. Still, it is tempting to speculate, that the His residue, which is conserved within the STAND class, may be the activating moiety in those enzymes that are capable of hydrolyzing ATP, similarly to aforementioned septins. Structures of STAND ATPases with TS analogs bound are needed for clarification of the activation mechanism and the positioning of **W_cat_**.

**Figure SF1_1.**
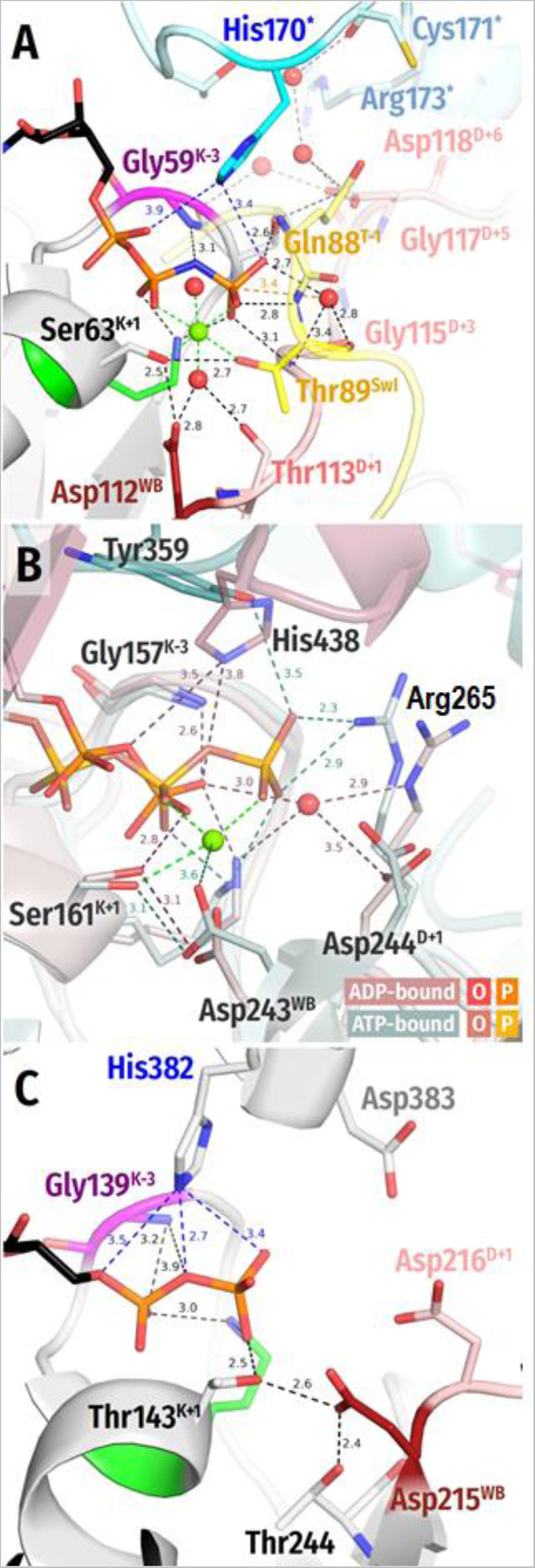
Tentative His fingers in septins and STAND ATPases. A. Structure of human septin 12 (PDB ID 6MQ9 [3]) complexed with GNP. Protein surrounding NTP-binding site is shown as a cartoon, functionally relevant residues are shown as sticks, water molecules are shown as red spheres, Mg^2+^ as a green sphere. P-loop lysine is shown in green, K-3 residue is shown in magenta. Asp^WB^ is shown in dark red and following residues in light red. Switch 1 residues are shown in yellow, residues belonging to adjacent monomer are highlighted in cyan and marked with an asterisk. All distances are given in ångströms. **B.** Superimposed structures of murine apoptotic peptidase activating factor 1 (Apaf-1): <u>In shades of pink:</u> ADP-bound form (PDB 3SFZ [5]). Water molecule is shown as a red sphere. HETHS domain is shown in a darker shade of pink. The distances are highlighted in dark pink. <u>In shades of teal:</u> dATP-bound form (PDB ID 3JBT [6]). Mg ion is shown as a green sphere. HETHS domain is shown in a darker shade of teal. The distances are highlighted in dark teal. **C.** Structure of the STAND protein with a tetratricopeptide repeat sensor PH0952 from *Pyrococcus horikoshii* (PDB ID 6MFV [7]). Colors as in panel A.

**VirB/PilT-like** class unites ATPases of the type IVa pili, GspE proteins of the type II secretion system, FlaI proteins of the archaeal flagellar system, and VirB11 proteins of the type IV secretion system [8–13]. In addition to the C-terminal nucleotide-binding P-loop domain, PilT-like ATPases have an N-terminal two-layer α/β sandwich domain, similar to the well-known ligand-binding PAS domain [14]. In the hexameric enzymes, the PAS-like domain provides the Arg finger to the AG site of the bound NTP molecule of the same subunit; an additional Arg finger can interact with γ-phosphate, see the structure in Fig. SF1_2A. In this structure, Ser^K+1^ forms a H-bond with Glu^E+4^ (highlighted in yellow), which, as shown on the panel, is connected to Glu^WB^ by a complex system of water bridges (highlighted in cyan). They resemble those in the PRC (see Fig. 10A in the main text), so that Glu^WB^ is likely to serve as a proton trap in PilT-like NTPases. The Asp^E+1^ residue on the C-cap is far from γ-phosphate. In the absence of TS-like structures, it cannot be fully excluded that Asp^E+1^ comes closer to the ATP molecule upon activation so that Asp^E+1^ interacts with W_cat_. Alternatively, Glu^E+4^ of the WB-crest can well perform this function and transmit a proton to the buried Glu^WB^.

**Fig. SF1_2.**
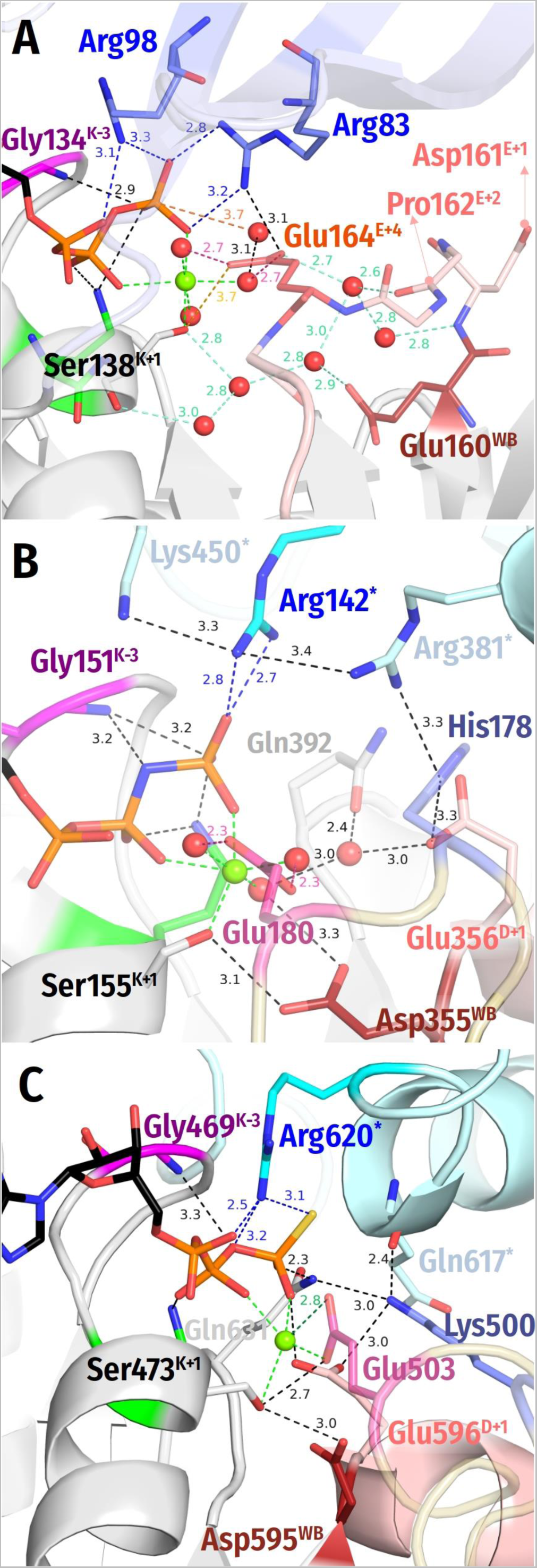
Nucleotide-binding sites of P-loop NTPases of VirB/PilT and FtsK-HerA classes. A. Structure of the twitching motility pilus retraction ATPase PilT4 from *Geobacter metallireducens,* PDB ID 6OJX [13]). Protein surrounding the NTP-binding site is shown as a cartoon, functionally relevant residues are shown as sticks, water molecules are shown as red spheres, Mg^2+^ as a green sphere. The P-loop lysine is shown in green, the K-3 residue is shown in magenta. Glu^WB^ is shown in dark red and the following residues in pale red. The PAS domain is shown in pale blue and its Arg fingers are shown in blue. All distances are indicated in Å. **B.** Crystal structure of the HerA hexameric DNA translocase from *Sulfolobus solfataricus* (PDB 4D2I[15]). Residues belonging to adjacent monomer are highlighted in cyan and marked with an asterisk. The additional Glu residue is shown in pink and the loop harboring it is shown in beige. Other colors as in panel A. **C.** Cryo-EM structure of FtsK from *Pseudomonas aeruginosa* PAO1 with resolution of 3.7 Å [16]. Colors as in panel B.

**FtsK-HerA** pumping ATPases are related to, but not monophyletic with the PilT-like ATPases. The ***HerA ATPases*** are present in all archaea and some bacteria [17] and participate in the repair of double strand DNA breaks [15]. The structure of HerA hexamer from *Sufolobus solfataricus*, in complex with AMP-PNP, a non-hydrolysable analogue of ATP, shows two Arg and one Lys residue pointing towards the triphosphate chain. The Glu355^WB^ residue interacts with Ser155^K+1^ and W6, whereas Glu356^D+1^ points towards the anticipated position of **W_cat_** [15], see Fig. SF1_2B. An additional class-specific Glu residue (Glu180 in the HerA hexamer from *Sufolobus solfataricus*, colored pink), similarly to Glu^E+4^ in PilT4 coordinates W3and W5, Mg^2+^ ligands #3 and #5.

The titular ***FtsK ATPases*** are motor protein that translocate double-stranded DNA during chromosome segregation. A recently reported cryo-EM structure of FtsK of *Pseudomonas aeruginosa* PAO1 with resolution of 3.7 Å indicates that an Arg residue analogous to the arginine finger of the AAA+ superfamily interacts with the NTP-binding site of the adjacent subunit Fig. SF1_2C [16]. The additional Glu residue is also present, but, contrary to HerA, it interacts with Ser^K+1^, similarly to Glu^D+4^ in PilT-like class ATPases, cf Fig. SF1_2A. The Asp^WB^ is also close, at only 3 Å away of Ser^K+1^. It is followed by Glu^D+1^ that points towards γ-phosphate and may stabilize W_cat_ in the TS.

Unless TS-like structures of FtsK, HerA, and VirB/PilT-like ATPases with their constricted catalytic sites are obtained, the functions and interactions of their additional Glu residues will remain obscure. Hence, the TS-like, constricted site in any of these ATPases may resemble either that of HerA and FtsK ATPases (where [Asp/Glu]^WB^ interacts with Ser^K+1^, and the additional Glu interacts with ligands of Mg^2+^) or that of PilT-like ATPases (where the additional Glu makes a H-bond with Ser^K+1^). It cannot be fully excluded that two carboxylic groups reach the [Ser/Thr]^K+1^ residue in the TS in the case of these NTPases.

**KAP ATPases** named after Kidins220/ARMS (mammalian neuronal membrane proteins) and PifA (F-plasmid protein involved in phage T7 exclusion), supposedly, use a AAA-type activation mechanism [18]. The KAP family proteins lack the Arg finger within the P-loop domain proper but carry a conserved arginine in the C-terminal helical segment, which could potentially function as an activating moiety, similar to Sensor 2 of the AAA+ ATPases [18]. Still, in the absence of structures of KAP proteins, both the activation mechanism and the **W_cat_** positioning in enzymes of this class remain unclear.

